# Formation and growth of co-culture tumour spheroids: new compartment-based mathematical models and experiments

**DOI:** 10.1101/2022.12.21.521515

**Authors:** Ryan J. Murphy, Gency Gunasingh, Nikolas K. Haass, Matthew J. Simpson

## Abstract

Co-culture tumour spheroid experiments are routinely performed to investigate cancer progression and test anti-cancer therapies. Therefore, methods to quantitatively characterise and interpret coculture spheroid growth are of great interest. However, co-culture spheroid growth is complex. Multiple biological processes occur on overlapping timescales and different cell types within the spheroid may have different characteristics, such as differing proliferation rates or responses to nutrient availability. At present there is no standard, widely-accepted mathematical model of such complex spatio-temporal growth processes. Typical approaches to analyse these experiments focus on the late-time temporal evolution of spheroid size and overlook early-time spheroid formation, spheroid structure and geometry. Here, using a range of ordinary differential equation-based mathematical models and parameter estimation, we interpret new co-culture experimental data. We provide new biological insights about spheroid formation, growth, and structure. As part of this analysis we connect Greenspan’s seminal mathematical model to co-culture data for the first time. Furthermore, we generalise a class of compartment-based spheroid mathematical models that have previously been restricted to one population so they can be applied to multiple populations. As special cases of the general model, we explore multiple natural two population extensions to Greenspan’s seminal model and reveal biological mechanisms that can describe the internal dynamics of growing co-culture spheroids and those that cannot. This mathematical and statistical modelling-based framework is well-suited to analyse spheroids grown with multiple different cell types and the new class of mathematical models provide opportunities for further mathematical and biological insights.

## 1 Introduction

Tumour spheroids are a key experimental tool to study avascular cancer progression and to develop cancer therapies [1, 2, 3, 4]. Spheroids bridge the gap between two-dimensional *in vitro* experiments and three-dimensional *in vivo* experiments. In comparison to two-dimensional experiments, three-dimensional spheroid experiments can capture realistic geometric limitations and spatial structure, such as those that arise due to differences in oxygen partial pressure at the periphery and centre of growing spheroids [5, 6, 7]. Spheroid experiments can be performed with a single cell type, referred to as a monoculture experiment, or performed with a mixture of two or more cell types, referred to as a co-culture experiment. Previously we have performed monoculture spheroid experiments and characterised the temporal evolution of spheroid size and structure using mathematical modelling and parameter estimation [5, 6, 7]. These studies are based on Greenspan’s seminal compartment-based mathematical model of avascular tumour growth [8]. Greenspan’s model captures the growth of a monoculture spheroid from an initial exponential growth phase, where each cell within in the spheroid can proliferate, to a limiting structure and size where proliferation at the periphery is thought to balance loss from the necrotic core [5, 6, 7, 8]. Furthermore, this model is relatively simple and all mechanisms and parameters in the model are biologically interpretable. This ordinary differential equation-based model is derived by coupling conservation of volume arguments with algebraic constraints (Section 4, Supplementary S5.1.1). Conservation of volume arguments incorporate biological mechanisms such as cell proliferation and cell death. Constraints, obtained by considering nutrient and waste mechanisms, define the boundaries of proliferating and necrotic regions. Using this framework, we have shown that transient and limiting spheroid structure can be independent of initial spheroid size [5]. We have also quantitatively compared a number of experimental designs to identify design choices that produce reliable biological insight and consistent parameter estimates [6]. Experimental design choices in this current work are informed by that study. Furthermore, we have revealed growth and adaptation mechanisms of spheroids to time-dependent oxygen availability [7]. As part of this study we aim to connect Greenspan’s monoculture model to co-culture data for the first time and generalise Greenspan’s model to multiple populations for the first time.

In this study, we focus on more complicated co-culture tumour spheroid experiments. These co-culture experiments are routinely performed to capture some of the complexity of *in situ* tumours, in particular the growth and interactions of multiple cell types that may be cancerous or healthy [9, 10, 11, 12]. These co-culture experiments are also routinely performed for convenience when a cancer cell line does not form an approximately spherical structure without the addition of another cell type, such as fibroblasts [11, 12]. We choose to perform experiments with human melanoma cancer cell lines in co-culture with fibroblasts. Using melanoma cell lines builds on our previous monoculture studies [5, 6, 7, 13] and using fibroblasts enhances spheroid formation and compactness. Each cell type within a co-culture spheroid may exhibit different characteristics, for example different proliferation rates or different responses to nutrient availability. Further, within each spheroid multiple biological processes occur on overlapping timescales. Therefore, characterising and interpreting co-culture spheroid growth is challenging. When taking a purely experimental approach quantitative differences and similarities in co-culture spheroid growth, and the mechanisms giving rise to these similarities and differences, are unclear. For example, how does the addition of fibroblasts influence the formation of melanoma spheroids and the structure of growing spheroids? Furthermore, what are the biological mechanisms driving internal dynamics of growing co-culture spheroids?

To address challenges interpreting spheroid formation and internal dynamics in growing co-culture spheroids, we perform two experiments and interpret the data using mathematical modelling and parameter estimation. Experiment 1 focuses on two-dimensional brightfield images taken from above each spheroid (Fig 1A). These images capture spheroid formation and growth of overall spheroid size. We aim to explore these new co-culture spheroid formation and growth data using an ordinary differential equation-based biphasic mathematical model [14]. Using the biphasic model allows us to explore *in vitro* spheroid formation, where cells placed in a well migrate and adhere over a timescale of days to form a compact solid mass. These mechanisms are not captured in Greenspan’s model. Furthermore, this approach extends traditional analysis of overall spheroid size that overlook formation [15, 16, 17]. Taking an extremely simple approach of approximating the early time growth as a straight line provides additional insight into the proliferation rate. However, images from Experiment 1 do not allow us to visualise the internal structure of growing spheroids.

**Figure 1:**
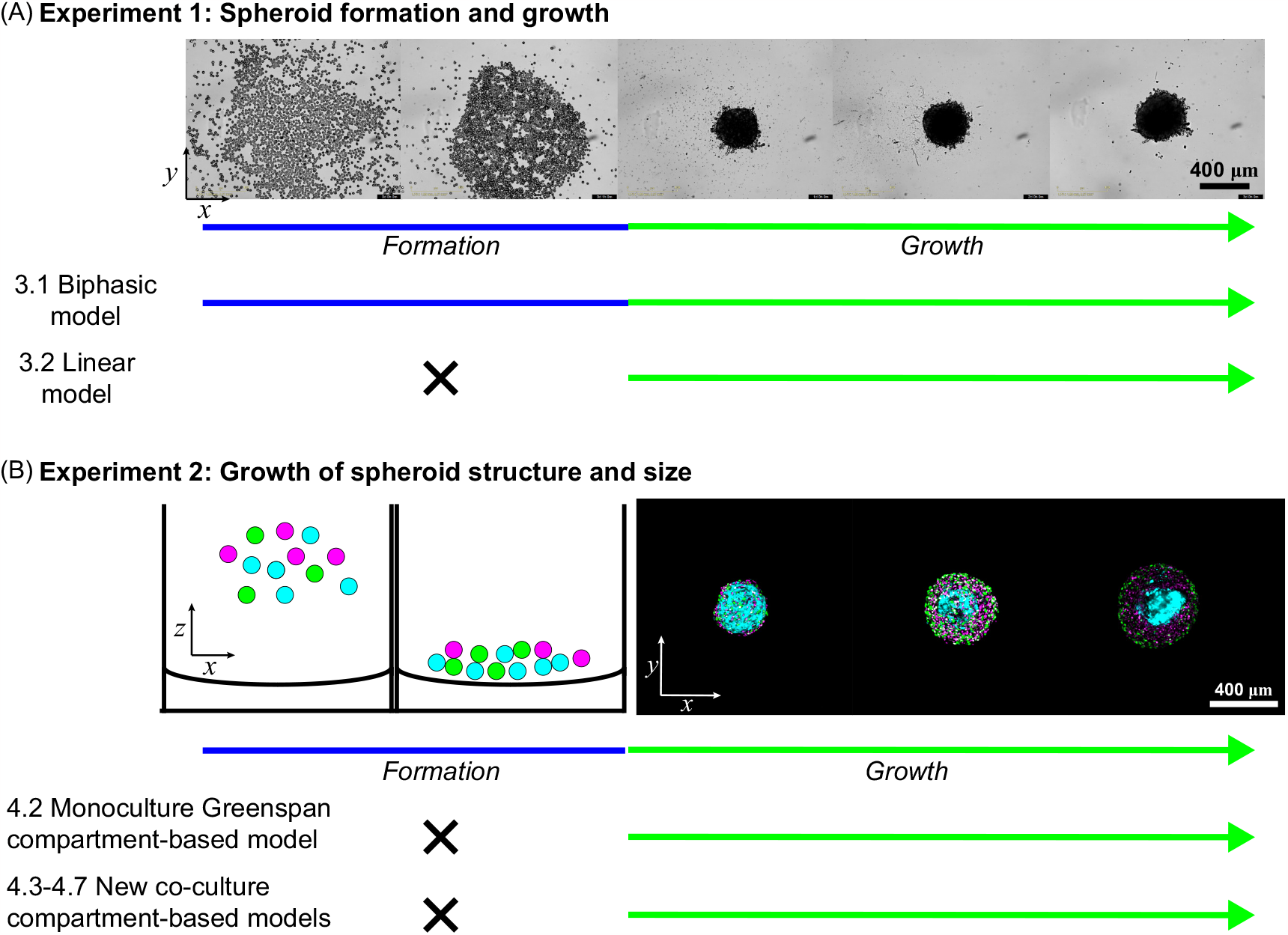
Workflow for characterising co-culture tumour spheroid growth using mathematical modelling and parameter estimation. (A) Experiment 1: Brightfield images of spheroid formation and overall size. Brightfield images are taken from above each co-culture melanoma fibroblast spheroid (M50:F50) at days 0, 1/24, 1, 2, and 3 after seeding. Images from Experiment 1 are used to estimate the projected area covered by cells, *R*(*t*). Data from Experiment 1 is interpreted with the biphasic model (§3.1) and linear model (§3.2). (B) Experiment 2: Growth of spheroid structure and size. Confocal images of the equatorial plane of a co-culture melanoma fibroblast spheroid (M25:F75) at days 2, 6, and 10 after seeding. Melanoma cells are transduced with FUCCI and colours represent FUCCI signals observed in experimental images: cells in gap 1 (G1) phase fluoresce red, shown in magenta for clarity; and cells in synthesis, gap 2, and mitotic (S/G2/M) phases fluoresce green shown in magenta and green. Fibroblasts are strained and shown in cyan. The internal dynamics and overall size of growing co-culture spheroids are analysed using the monoculture Greenspan model (§4.2) and a number of new co-culture models that are natural extensions of Greenspan’s model (§4.3-§4.6). Labelling of model corresponds to the sections in the manuscript where the model is described.

Experiment 2 uses confocal microscopy to reveal the internal structure of growing co-culture spheroids, after the compact solid mass has formed (Fig 1B). This technique allows us to focus on the equatorial plane of each spheroid and generate three-dimensional renderings of spheroid structure. Melanoma cells are transduced with fluorescent ubiquitination-based cell cycle indicator (FUCCI) to visualise the position and cell cycle status of each melanoma cell within the spheroid [13, 18]. Fibroblasts are stained with a cell tracker to observe their position within the spheroid. To characterise and interpret the internal dynamics of growing co-culture spheroids we aim to connect Greenspan’s monoculture mathematical model to co-culture data for the first time. It is unclear before performing this analysis how much insight can be gained by connecting Greenspan’s monoculture model to co-culture data. For example, can we identify differences in parameter estimates between melanoma spheroids without fibroblasts and spheroids in co-culture with fibroblasts? Furthermore, we aim to produce a generalised class of monoculture compartment-based mathematical models to multiple populations for the first time. In doing so we aim to extend Greenspan’s model and a number of related compartment-based mathematical models to multiple populations. Therefore our work can be considered to be a genersalisation of models described by Burton [19], Deakin [20], McElwain and colleagues [21, 22], Adam and colleagues [23, 24, 25, 26, 27, 28], Landry [29] as well as those presented by Byrne [30, 31] Using the general compartment-based model, we aim to efficiently and systematically develop multiple natural extensions of Greenspan’s model to two populations. Exploring these new co-culture models we aim to reveal biological mechanisms that can describe our new co-culture experimental data and those that cannot. Other mathematical models describing tumour spheroid growth are reviewed in [32, 33, 34, 35]. For example, individual-based models that prescribe rules that govern the dynamics of each cell [36, 37, 38] or continuum models in the form of partial differential equations [39, 40]. However, these other models typically contain parameters that are challenging to estimate using data collected in this study [17].

This study is structured as follows. Section §2 outlines key experimental methods. Section §3 details the biphasic and linear mathematical models used to interpret data from Experiment 1 that focuses on spheroid formation and growth of overall spheroid size. Section §4 presents the general compartment-based mathematical model and multiple new two-population extensions of Greenspan’s model that allow us interpret data from Experiment 2 that focuses on internal dynamics of growing co-culture spheroids. Section §5 explains methods for parameter estimation and identifiability analysis. Results and discussion, and the conclusion are presented in §6 and §7, respectively. Supplementary material includes additional details of experimental methods; a summary of the experimental data; additional experimental images; additional mathematical modelling details; numerical methods; and additional results.

## 2 Experimental methods

We perform two co-culture experiments (Fig 1). Experiment 1 focuses on two-dimensional brightfield images taken from above each spheroid (Fig 1A) and focuses on spheroid formation and growth of overall spheroid size. Experiment 2 uses confocal microscopy to reveal the size, internal structure, and geometry of growing co-culture spheroids, after the compact solid mass has formed (Fig 1B). Throughout we mix one human melanoma cell line (WM983B or 1205Lu [13, 41, 42]) with human primary fibroblasts (QF1696 [43]) at four different initial compositions (M100:F0, M75:F25, M50:F50, M25:F75), where M*x*:F*y* is the initial proportion, measured as a percentage, of melanoma cells *x* [%] and fibroblasts *y* = 100 − *x* [%]). Both melanoma cell lines are transduced with fluorescent ubiquitination-based cell cycle indicator (FUCCI) constructs that allow us to visualise the cell-cycle status of each melanoma cell continuously throughout time without loss of signal [13, 18]. In experimental images from Experiment 2 FUCCI-transduced melanoma cells in gap 1 (G1) phase fluoresce red, shown in magenta for clarity; and FUCCI-transduced melanoma cells cells in synthesis, gap 2, and mitotic (S/G2/M) phases fluoresce green shown in magenta and green. Fibroblasts are stained and shown in cyan. Further details of experimental methods including cell culture; spheroid generation, culture, and experiments; imaging and image processing, see Supplementary S1.

## 3 Mathematical models to analyse spheroid formation and size

In Experiment 1 we observe and measure spheroid formation and growth of overall spheroid size, *R*(*t*), using brightfield images taken from above each spheroid. Here, to analyse and interpret measurements of *R*(*t*) we consider two models: (i) a biphasic model to explore formation and growth (Fig 2A); and (ii) a linear model to approximate early time growth dynamics after formation (Fig 2B).

**Figure 2:**
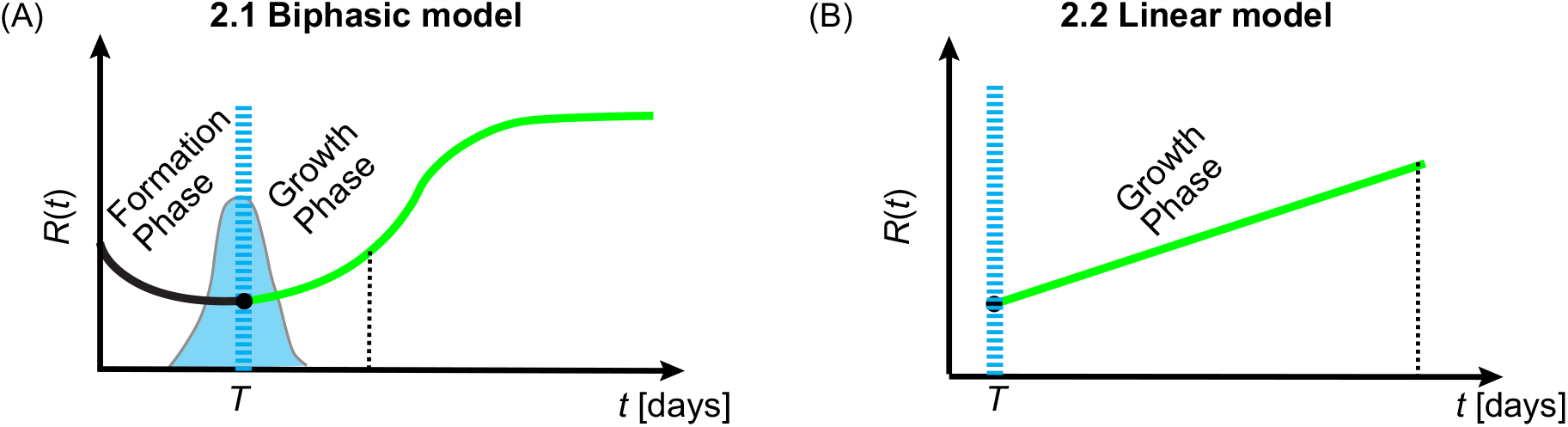
Mathematical models to analyse spheroid formation and size. (A) Biphasic model to explore formation and growth phases. (B) Linear model to approximate early time growth phase dynamics after formation. In (A,B) blue dashed vertical corresponds to the time, *T*, when the spheroid formed as a compact solid mass.

### 3.1 Biphasic model for spheroid formation and growth of overall size

We analyse three-dimensional spheroid formation using the biphasic model introduced in [14] (Fig 1C). In the first phase, cells that are placed in the well of a tissue culture plate migrate and adhere to form a compact solid mass (Fig 1A,B). The projected area covered by cells, as viewed from above, shrinks until the spheroid forms, for example compare the yellow masks from 4 hours to 48 hours in Figure S1. In the second phase, the spheroid grows as a three-dimensional compact mass. We describe each phase using a logistic-type model with distinct parameters,

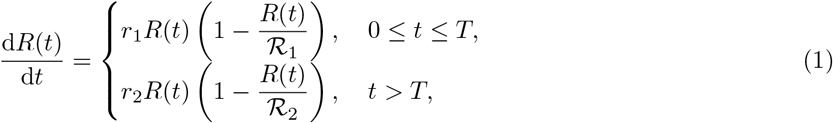

where *R*(*t*) [μm] describes the equivalent radius of the projected area covered by cells at time *t*, and *T* [days] is the formation time. In the second phase, *t > T* [days], *R*(*t*) is equivalent to the radius of a compact spheroid at time *t*. Further, *r*_1_ [day^−1^] is associated with the rate at which the spheroid forms; *ℛ*_1_ [μm] is the limiting radius of the spheroid in the first phase; *r*_2_ [day^−1^] is the growth rate of the spheroid as a compact mass; and *ℛ*_2_ [μm] is the long-time maximum spheroid radius. The initial spheroid size, *R*(0) [μm], satisfies *R*(0) *> R*_1_ to observe shrinking in the first phase. Hence, *R*(*t*) *> ℛ*_1_ for 0 ≤ *t* ≤ *T* . In the second phase, *t > T, R*(*t*) *→ ℛ*_2_ as *t → ∞*. This model is characterised by six parameters *θ* = (*R*(0), *r*_1_, *r*_2_, *ℛ*_1_, *ℛ*_2_, *T*) that we will estimate from our experimental data.

### 3.2 Linear model for growth of overall spheroid size

In contrast to the biphasic model in §3.1 that can be used to describe spheroid formation and growth, we now restrict our attention to growth after the spheroid has formed. For spheroids that do not appear to reach a long-time maximum size during the timescale of our experiments we simplify our analysis. We approximate early time exponential growth dynamics in the second phase of the model presented in Eq. (1) with a linear model (Fig 1D, Supplementary S3.1), i.e. exp(*λt*) = 1 + *λt* + *O*((*λt*)^2^) when *λt* is sufficiently small [44], to give

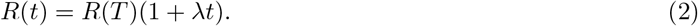

where *R*(*t*) [μm] is the radius of the spheroid at time *t, R*(*T*) [μm] is the radius of the spheroid at the end of formation that we treat as a constant parameter, and *λ* [day^−1^] is the growth rate. This model is characterised by two parameters *θ* = (*R*(*T*), *λ*) that we will estimate from our experimental data.

## 4 Compartment-based mathematical models to analyse internal dynamics of growing spheroids

In Experiment 2 we use confocal microscopy to capture the internal structure of growing co-culture spheroids after they have formed as a compact solid mass. This data comprises of radial measurements of overall spheroid size, *R*(*t*), radial measurements of the necrotic core, *R*_n_(*t*), and images showing the spatiotemporal evolution of melanoma cells and fibroblasts throughout the spheroid. Here, to interpret these data we seek to use and extend Greenspan’s monoculture compartment-based model [8]. To efficiently and systematically obtain multiple two population extensions of Greenspan’s model, we derive a general compartment-based spheroid model composed of *I* populations and *J* compartments. Greenspan’s seminal model is then a special case (Supplementary S5.1).

We derive the general compartment-based spheroid model from conservation of volume arguments subject to constraints that define the boundaries of the compartments. We refer to these constraints as *boundary constraints*. Boundary constraints, for example to define the size of proliferating region, may be prescribed [17] or arise by considering additional biological mechanisms, for example oxygen, nutrient, or waste mechanisms [8]. Conservation of volume arguments can incorporate biological mechanisms such as cell proliferation, cell death, and cell migration. In this framework, a population could refer to a cell type, for example melanoma or fibroblasts, or the state of the cell, for example living or dead. Here, when we refer to a population we intend the former, but the framework is general and either interpretation can be considered.

To introduce our general model we first explore a reduced version of Greenspan’s monoculture model (Fig 3A-C). We choose to focus on this two compartment model, comprising of a proliferating region at the periphery and a necrotic core, since we have measurements of *R*(*t*) and *R*_n_(*t*) and so that the multiple population extensions are simpler to interpret. Furthermore, this model is relatively simple and all mechanisms and parameters are biologically interpretable. This monoculture model has three parameters associated with cell proliferation rate, loss from the necrotic core, and an oxygen threshold for cell death that corresponds to the formation of the necrotic core. As the model is restricted to one population this model cannot be used to describe the internal dynamics of multiple populations within the spheroid.

**Figure 3:**
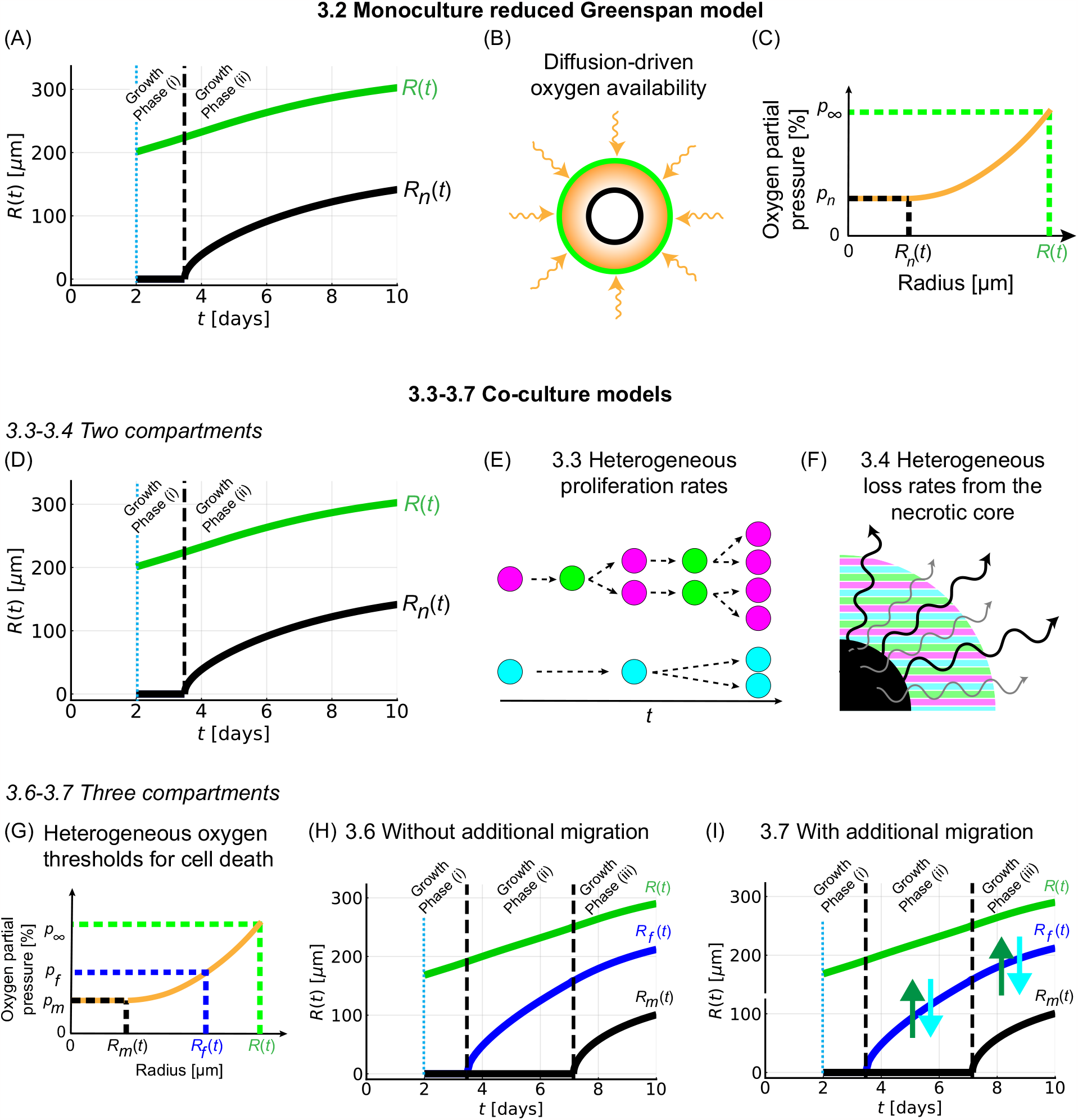
Compartment-based mathematical models to analyse spheroid structure. (A) Simulation of the monoculture reduced Greenspan model (Section 4.2) that is comprised of two compartments at later times: a proliferating region at the periphery and a necrotic core. (B-C) Oxygen diffuses within the spheroid and is consumed by living cells which gives rise to gradient in the oxygen partial pressure within the spheroid. Cells are assumed to die below an oxygen partial pressure threshold *p*_*n*_ [%]. (D-I) Four natural extensions of the monoculture reduced Greenspan model to two populations: (i) Co-culture Model 1 (Section 4.3) - a two-compartment co-culture model with heterogeneous proliferation rates shown in (D,E); (ii) Co-culture Model 2 (Section 4.4) - a two-compartment co-culture model with heterogeneous loss rates from the necrotic core shown in (D,F) (grey/black arrows indicate loss of necrotic matter from the necrotic core (black) from melanoma and fibroblast cells in various stages of degradation); (iii) Co-culture Model 3 (Section 4.5) - a three-compartment co-culture model with heterogeneous oxygen thresholds for cell death shown in (G,H); and (iv) Co-culture Model 4 (Section 4.6) - a three-compartment co-culture model with heterogeneous oxygen thresholds for cell death with additional cell migration (green/blue arrows for melanoma/fibroblasts, respectively) shown in (G,I). In (G) Melanoma cells and fibroblasts are assumed to die below an oxygen partial pressure thresholds *p*_m_ [%] and *p*_f_ [%], respectively. Two/three compartment models undergo two/three growth phases. Note that (A) and (D) look similar as they are both represent models with two compartments. Colours in (E-F) are described in Fig 1.

To explore mechanisms that could describe the internal dynamics of multiple populations within growing spheroids, we consider four natural extensions on the monoculture reduced Greenspan model. Each extension is developed by generalising one of the monoculture model parameters. The models are as follows: (i) a two-compartment co-culture model with heterogeneous proliferation rates (Fig 3D,E); (ii) a two-compartment co-culture model with heterogeneous loss rates from the necrotic core (Fig 3D,F); (iii) a three-compartment co-culture model with heterogeneous oxygen thresholds for cell death (Fig 3G,H); and (iv) a three-compartment co-culture model with heterogeneous oxygen thresholds for cell death with additional cell migration (Fig 3G,I).

### 4.1 General model with *I* populations and *J* compartments

We assume that the tumour spheroid is a compact mass maintained by surface tension or cell-cell adhesion, however the details of such mechanisms are not explicitly modelled nor crucial to model description [8]. We further assume that each compartment within the tumour spheroid is spherically symmetric, populations are well-mixed within each compartment, and all compartments are concentric. In Figure 4 we present a schematic of the general model with *I* populations and *J* compartments. In Table 1 we present model variables and descriptions. Compartment *j* denotes the region *R*_*j*_(*t*) *< r < R*_*j*−1_(*t*) for *j* = 1, 2, …, *J* − 1 and compartment *J* denotes the region 0 *< r < R*_*J*−1_(*t*). Boundary constraints, for example to define the size of proliferating region, may be prescribed [17] or arise by considering additional biological mechanisms, for example oxygen, nutrient, or waste mechanisms [8].

**Table 1:**
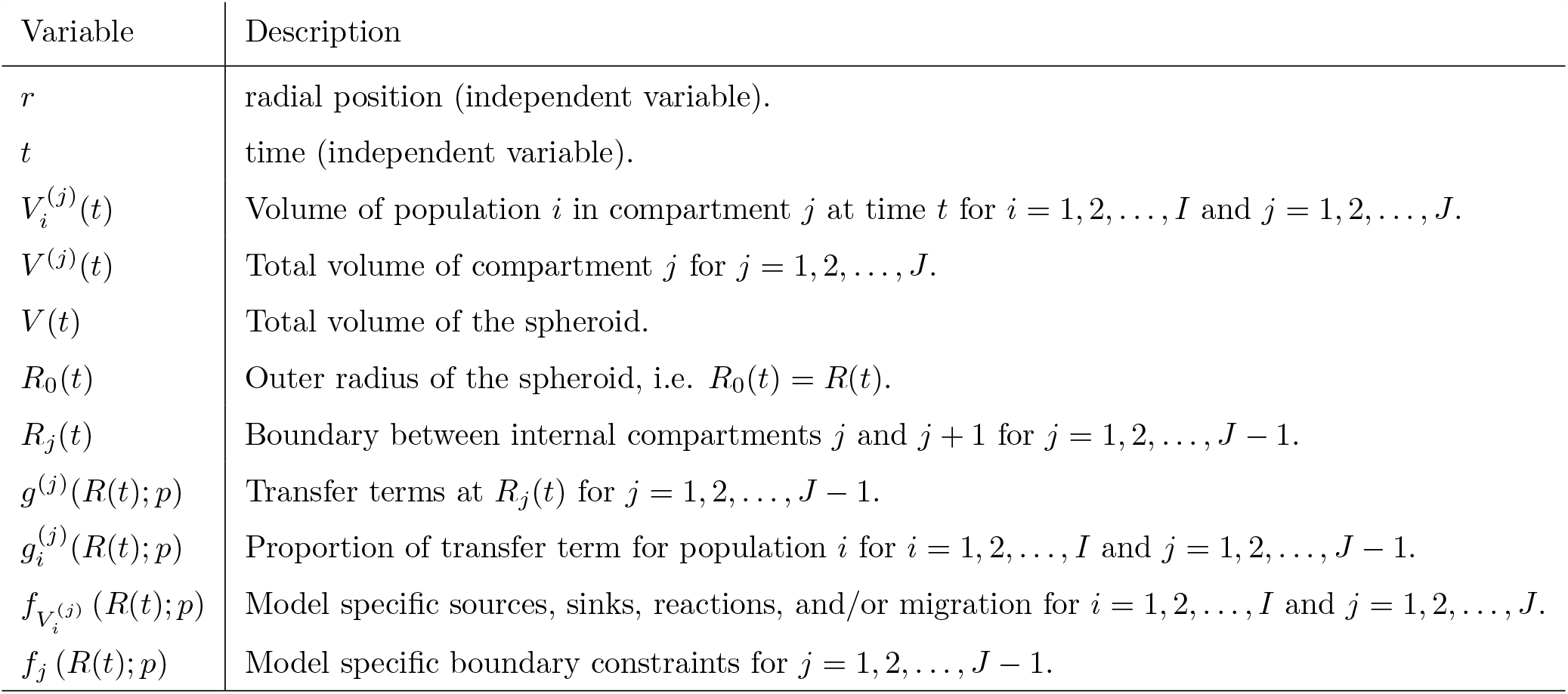
General compartment-based model variables and descriptions.

**Figure 4:**
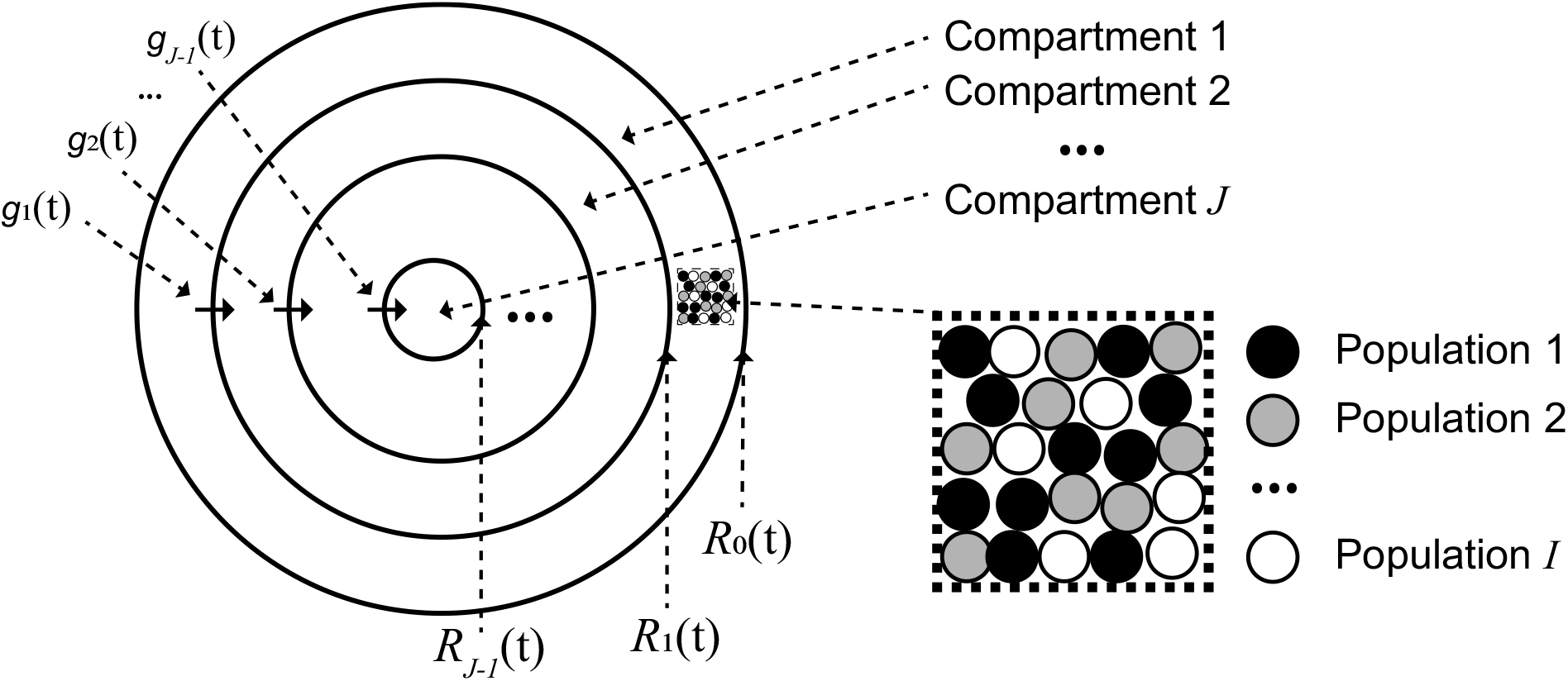
Schematic for general compartment-based tumour spheroid model with *I* populations and *J* compartments. The direction of arrows representing *g*_*i*_ for *i* = 1, …, *J* − 1 corresponds to the direction in a growing spheroid where d*R*_*j*_ (*t*)*/*d*t >* 0 for *j* = 0, 1, …, *J* − 1.

A general conservation of volume statement for *I* populations and *J* compartments, subject to boundary constraints (Eqs (3.6), is shown in Eqs (3.1)-(3.5). Equation (3.1) incorporates sources, sinks, reactions, and/or migration. Equation (3.1) also incorporates *transfer terms* that arise due to internal boundary movement and ensure conservation of volume. To interpret these transfer terms consider an illustrative example with a Greenspan-type model. Assume that at time *t* the spheroid is composed of two compartments: a necrotic core in 0 ≤ *r* ≤ *R*_n_(*t*), and a region *R*_n_(*t*) *< r* ≤ *R*(*t*) where cells are living and proliferating. Further, assume that over a time of duration Δ*t* the spheroid increases in volume due to proliferation at the periphery exceeding loss from the necrotic core, i.e. *R*(*t*) *< R*(*t*+Δ*t*), and due to a boundary constraint that arises from oxygen diffusion and consumption mechanisms *R*_n_(*t*) *< R*_n_(*t* + Δ*t*). In this situation, transfer terms capture cells in the region *R*_n_(*t*) *< r < R*_n_(*t* + Δ*t*) transferring from a compartment representing living cells at time *t* to a different compartment representing necrotic cells at time *t* +Δ*t*. Note that internal compartments change drastically in size while overall spheroid size can remain approximately constant, for example in experiments with time-dependent oxygen availability [14]. In general, as we assume a well-mixed population we set the transfer term for population *i* across the boundary *R*_*j*_(*t*) to be proportional to the volume fraction of population *i* in compartment *j*, i.e. we set 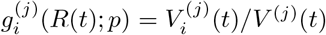 for *i* = 1, 2, …, *I* and *j* = 1, 2, …, *J* − 1. This choice of 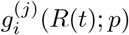 satisfies Eq (3.2) which ensures that the total transfer of volume across the boundary *R*_*j*_(*t*) is *g*^(*j*)^(*t*). Further, enforcing that each *g*^(*j*)^(*R*(*t*); *p*) is non-negative in Eq (3.2) ensures that transfer of volume across *R*_*j*_(*t*) is in the same direction for all populations.

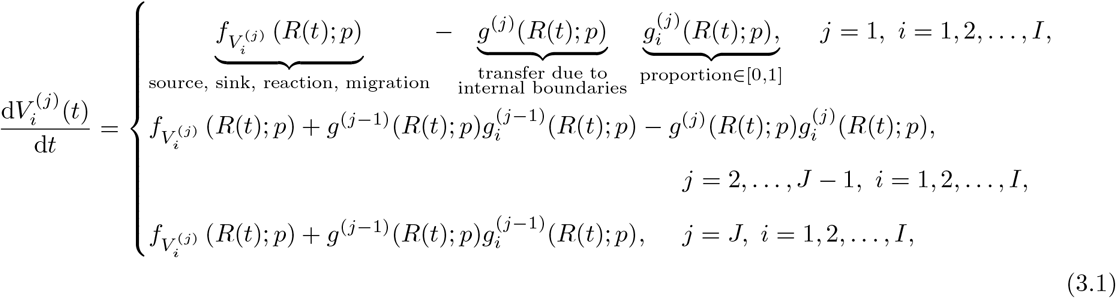

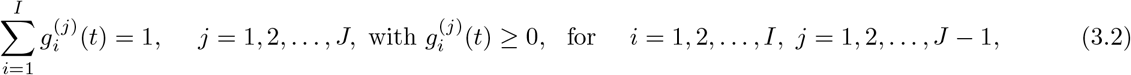

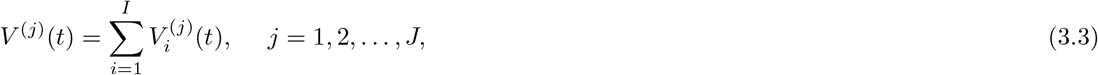

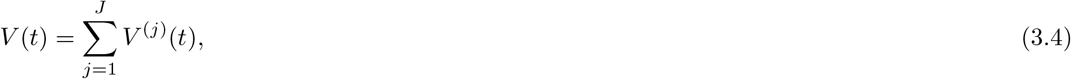

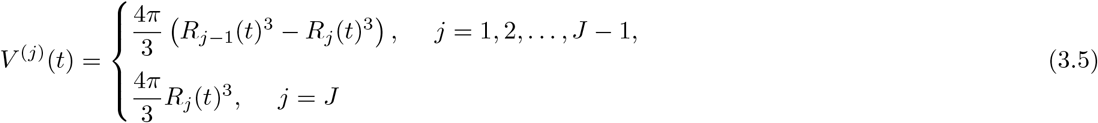

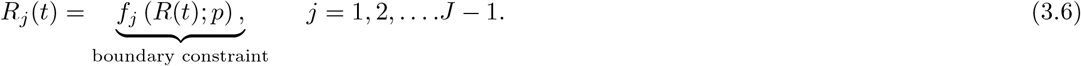

Equation (3.3) ensures there are no voids in each compartment by stating that the total volume of a compartment is equal to the sum of the volume of each cell population in that compartment. Equation (3.4) ensures there are no voids in the spheroid by stating that the total volume of the spheroid is equal to sum of the volumes of each compartment that make up the spheroid. Equation (3.5) enforces the assumption of spherical symmetry for each compartment within the spheroid making use of the fact that the volume of a sphere of radius *r* is 4*πr*^3^*/*3.

To solve Eq (3.1)-(3.6) we prescribe the following functions: (i) 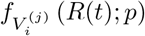 for *i* = 1, 2, …, *I* and *j* =1, 2, …, *J* that describe sources, sinks, reactions, and/or migration; and (ii) *f*_*j*_ (*R*(*t*); *p*) for *j* = 1, 2, …, *J*−1 that describe boundary constraints. These prescribed functions depend on *R*(*t*) and a vector of model parameters, *p*, that are also to be prescribed. Eqs (3.1), (3.3)-(3.6) form a system of *IJ* + 3*J* equations for *IJ* + 3*J* unknowns 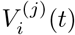 for *i* = 1, 2, …, *I* and *j* = 1, 2, …, *J*; *g*^(*j*)^(*t*) for *j* = 1, 2, …, *J* − 1; *V* ^(*j*)^(*t*) for *j* = 1, 2, …, *J*; *V* (*t*); and *R*_*j*_ for *j* = 0, 1, 2, …, *J* − 1).

It is useful to reformulate Eqs (3.1)-(3.6) to obtain expressions for the temporal evolution of overall spheroid size (Eq (4.2)) and to obtain expressions for the transfer terms (Eq (4.3)). To obtain these expressions we simplify Eqs (3.1)-(3.6) from a system of *IJ* + 3*J* to the following system of *IJ* + 2*J* − 1 equations for *IJ* +2*J*−1 unknowns 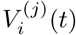 for *i* = 1, 2, …, *I* and *j* = 1, 2, …, *J*; *g*^(*j*)^(*t*) for *j* = 1, 2, …, *J*−1; and *R*_*j*_(*t*) for *j* = 0, 1, 2, …, *J* − 1) (see Supplementary S3.2 for further details).

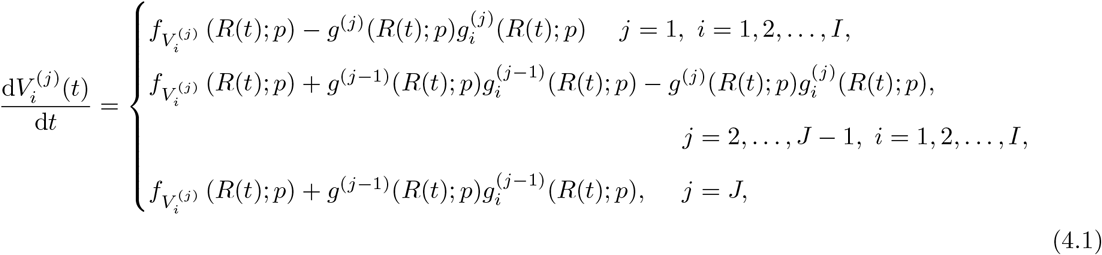

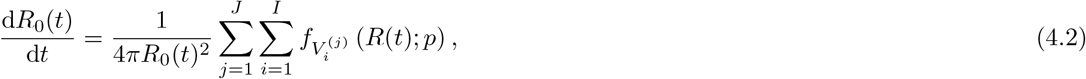

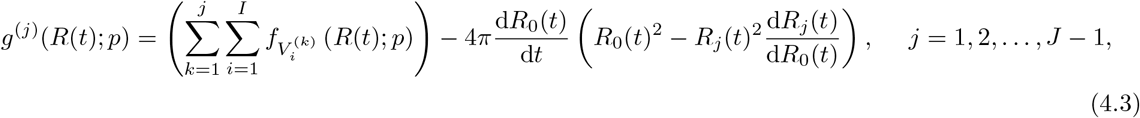

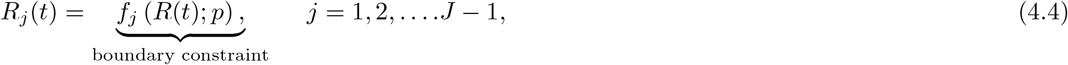

where it is assumed that an expression for d*R*_*j*_(*t*)*/*d*R*_0_(*t*) in terms of *R*_0_(*t*) for *j* = 1, 2, …, *J* − 1 can be derived from the prescribed functions in Eq (4.4). We then solve Eqs (4.1)-(4.4) numerically (Supplementary S4). Given 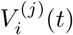 for *i* = 1, 2, …, *I* and *j* = 1, 2, …, *J* from the solution of Eq (4) we obtain *V* ^(*j*)^(*t*) for *j* = 1, 2, …, *J* and *V* (*t*) using Eqs (3.3)-(3.4), respectively. Motivated by our new experimental data, we will now describe four simplifications of our general model so that it applies to our particular experimental conditions.

### 4.2 Monoculture reduced Greenspan model

The monoculture reduced Greenspan model considers one population and two compartments (*I* = 1, *J* = 2), captures key characteristics of Greenspan’s model, and describes growth with two phases (Fig 3A-C). In phase (i) all cells within the spheroid can proliferate and the spheroid grows exponentially. In phase (ii) the spheroid is composed of two compartments: compartment one which is a proliferating region at the periphery, *R*_1_(*t*) *< r < R*(*t*), and compartment two which is the necrotic core, 0 ≤ *r* ≤ *R*_1_(*t*). At later times proliferation at the periphery balances mass loss from the necrotic core resulting in a limiting spheroid structure [5, 6, 7, 8]. For consistency with the literature we write *R*_1_(*t*) as *R*_n_(*t*) and *R*_0_(*t*) as *R*(*t*) [6, 7, 8]. We define *R*_n_(*t*) through a boundary constraint obtained by considering additional biological mechanisms, specifically oxygen diffusion within the spheroid and oxygen consumption by living cells (detailed in Supplementary S3.3) [7].

We assume that in the proliferating region the rate at which cell volume is produced by mitosis per unit volume of living cells is *s* [day^−1^]. In the necrotic region we assume that the rate at which cell volume is lost from the necrotic core per unit volume of necrotic material is 3*sγ* [day^−1^] where *γ* [-] is a dimensionless parameter and the three is included for mathematical convenience and consistency with the literature [8]. Therefore, we prescribe the following functions,

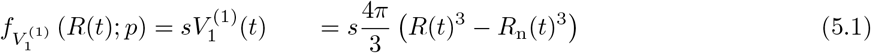

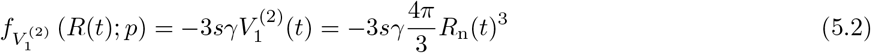

Substituting the prescribed functions (Eqs (5) and (S.14)) into general model Eqs (4.1-4.4) gives equations for the temporal evolution of the volume of living cells at the periphery (Eq (6.1)), the temporal evolution of the volume of the necrotic core (Eq (6.2)), the temporal evolution of overall spheroid size (Eq (6.3)), an equation for the rate of transfer of living cells to the necrotic core (Eq (6.4)), and the algebraic boundary constraint that arises from oxygen diffusion and consumption mechanisms (Eq (6.5))

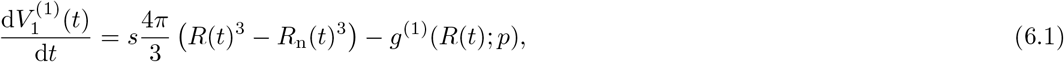

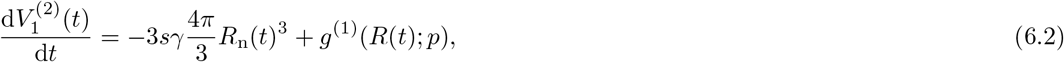

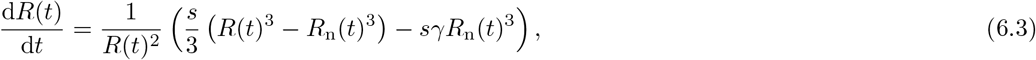

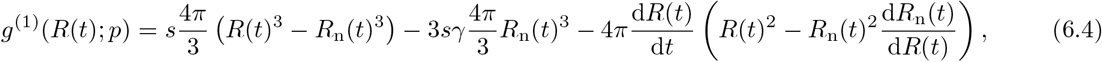

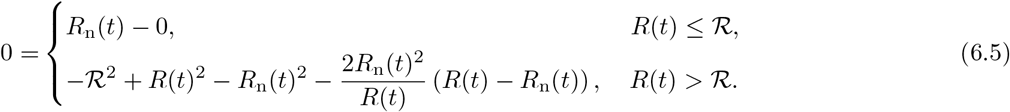

where *ℛ* is the radius of the spheroid when the necrotic first forms and is associated with the oxygen threshold *p*_n_ [%] below which cells die (Fig 3C, Supplementary S3.3).

In general Eqs (4.1-4.4) are coupled and must be solved simultaneously to obtain *R*(*t*) and *R*_n_(*t*). Here, for the special case of this monoculture model, Eqs (6.3) and (6.5) can be solved for *R*(*t*) and *R*_n_(*t*) independently of Eqs (6.1), (6.2), and (6.4). Solving only Eqs (6.3) and (6.5) is the standard formulation of a monoculture compartment-based Greenspan-type model. Therefore, for consistency with the literature we present the final model as only Eqs (6.3) and (6.5)

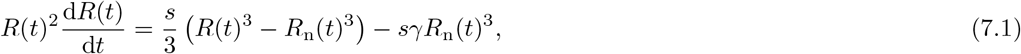

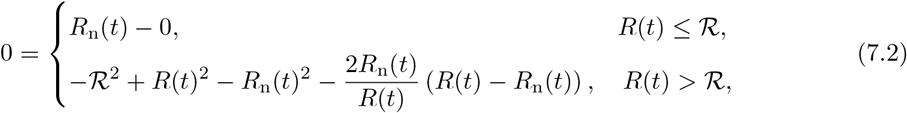

where *s* [day^−1^] is the rate at which cell volume is produced by mitosis per unit volume of living cells; *γ* [-] is a dimensionless parameter associated with loss of mass from the necrotic core; *ℛ* [μm] is radius of the spheroid when the necrotic region forms associated with the oxygen partial pressure threshold *p*_n_ [%]; *R*(*T*) [μm] is the initial radius of the spheroid at formation that we treat as a constant parameter; and we set *R*_n_(*T*) = 0 based on experimental observations. This model is characterised by four parameters *θ* = (*R*(*T*), *s, γ, ℛ*) that we will estimate from our experimental data.

### 4.3 Co-culture Model 1: Heterogeneous proliferation rates

Here we extend the monoculture reduced Greenspan model presented in §4.2 by generalising the rate at which cell volume is produced by mitosis per unit volume of living cells, *s* [day^−1^], to be two potentially distinct parameters (Fig 3D,E). This results in a model with two populations and two compartments (*I* = 2, *J* = 2). We let population one and two represent melanoma and fibroblast cells, respectively. Accordingly we update the notation and let *s*_m_ [day^−1^] and *s*_f_ [day^−1^] corresponding to the rates at which cell volume is produced by mitosis per unit volume of living cells for melanoma cells and fibroblasts, respectively (Figure 1I). We assume that the rate at which cell volume is lost from the necrotic core per unit volume of necrotic material is 3*s*_m_*γ* [day^−1^]. All other model assumptions and mechanisms are unchanged. The prescribed functions for Co-culture Model 1 are

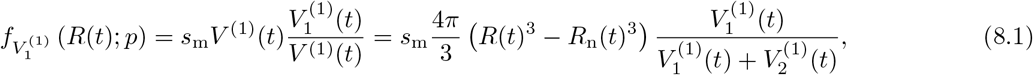

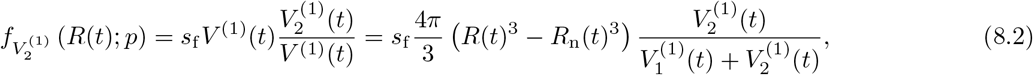

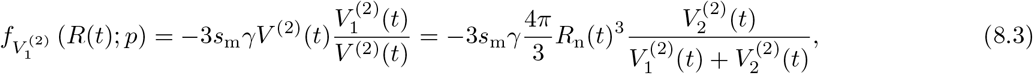

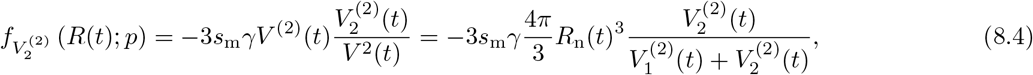

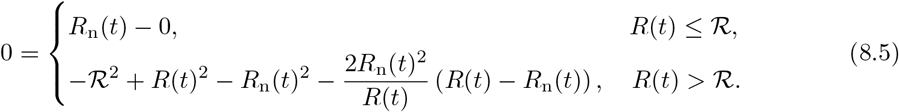

Substituting Eqs (8.1)-(8.5) into Eqs (4.1-4.4) gives equations for the temporal evolution the volume occupied by each population in each compartment (Eqs (S.15.1)-(S.15.4)), the temporal evolution of overall spheroid size (Eq (S15.5)), and an equation for the rate of transfer of living cells to the necrotic core (Eq (S.15.6).

We solve the system of Eqs (S.15.1)-(S.15.6) with Eqs (8.5) and (S.16) numerically. Eq (S.16) governs the time evolution of d*R*_n_(*t*)*/*d*R*(*t*), obtained by differentiating Eq (8.5) with respect to *R*(*t*). To specify initial conditions we introduce *V*_m_(*T*) = *V*_1_(*T*) as the volume of melanoma cells at spheroid formation, *t* = *T*, and treat this as a parameter. The volume of fibroblast cells at *t* = *T* is then *V*_2_(*T*) = *V*_f_ (*T*) = 4*πR*(*T*)^3^*/*3 − *V*_m_(*T*). This model is characterised by six parameters *θ* = (*R*(*T*), *V*_m_(*T*), *s*_m_, *s*_f_, *γ, R*).

### 4.4 Co-culture Model 2: Heterogeneous loss rates

Here we extend the monoculture reduced Greenspan model presented in §4.2 by generalising the dimensionless parameter *γ* associated with rate at which mass is lost from the necrotic core to be two potentially distinct parameters (Fig 3D,F). This results in a model with two populations and two compartments (*I* = 2, *J* = 2). We let population one and two represent melanoma and fibroblast cells, respectively. We assume that the rate at which cell volume is lost from the necrotic core per unit volume of necrotic material is 3*sγ*_*m*_ [day^−1^] for melanoma cells and is 3*sγ*_*m*_ [day^−1^] for fibroblasts. All other model assumptions and mechanisms are unchanged. The prescribed functions for Co-culture Model 2 are

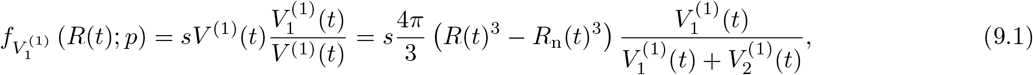

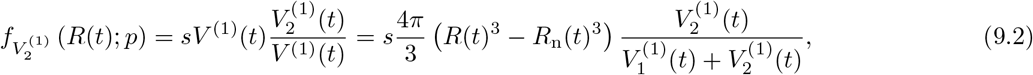

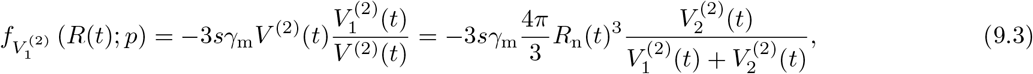

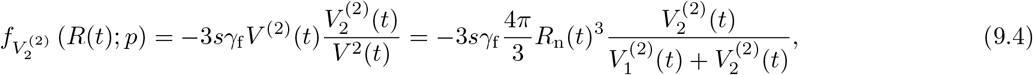

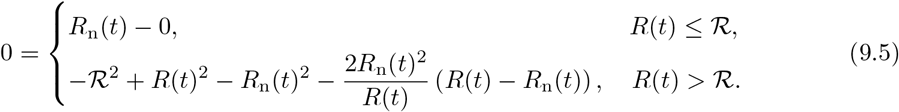

Substituting Eqs (9.1-9.5) into Eqs (4.1-4.4) gives equations for the temporal evolution of the volume occupied by each population in each compartment (Eqs (S.17.1)-(S.17.4)), the temporal evolution of overall spheroid size (Eq (S.17.5)), and an equation for the rate of transfer of living cells to the necrotic core (Eq (S.17.6)). We solve the system of Eqs (S.17.1)-(S.17.6) with Eqs (9.1 and (S.16) numerically. This model is characterised by six parameters *θ* = (*R*(*T*), *V*_m_(*T*), *s, γ*_m_, *γ*_f_, *ℛ*).

### 4.5 Co-culture Model 3: Heterogeneous cell death oxygen thresholds

Here we extend the monoculture reduced Greenspan model (§4.2) by generalising the single oxygen partial pressure threshold *p*_n_ [%] to two potentially distinct parameters *p*_m_ [%] and *p*_f_ [%] (Fig 3G,H). The new parameters *p*_m_ and *p*_f_ correspond to the oxygen partial pressure below which melanoma (population 1) and fibroblast (population 2) cells die, respectively (Figure 1K,L). We assume *p*_f_ ≥ *p*_m_ with the aim of reproducing experimental observations with regards to the position of fibroblasts within the spheroid. Generalising *p*_n_ corresponds to generalising the parameter *ℛ* (Eq S.13) to two potentially distinct parameters *ℛ*_m_ [μm] and *ℛ*_f_ [μm]. The new parameters *ℛ*_m_ and *ℛ*_f_ depend on *p*_m_ and *p*_f_, respectively.

The resulting model describes two populations with three compartments (*I* = 2, *J* = 3). In phase (i) all cells proliferate and the spheroid grows exponentially. In phase (ii), the in *R*_f_ (*t*) ≤ *r* ≤ *R*(*t*) melanoma cells and fibroblasts proliferate and in 0 ≤ *r* ≤ *R*_f_ (*t*) melanoma cells proliferate and fibroblasts are necrotic. In phase (iii), in compartment one, *R*_f_ (*t*) ≤ *r* ≤ *R*(*t*), melanoma cells and fibroblasts proliferate; in compartment two, *R*_m_(*t*) ≤ *r* ≤ *R*_f_ (*t*), melanoma cells proliferate while in fibroblasts are necrotic; and in compartment three, 0 ≤ *r* ≤ *R*_m_(*t*), melanoma and fibroblast cells are necrotic. We define *R*_m_(*t*) and *R*_f_ (*t*) through boundary constraints obtained by considering diffusion of oxygen within the spheroid, consumption of oxygen by living cells, and the oxygen thresholds *p*_m_ and *p*_f_ below which melanoma cells and fibroblasts undergo necrosis, respectively (Fig 3G). Further details on these boundary constraints are presented in Supplementary S3.6. All other model assumptions and mechanisms are unchanged. The prescribed functions for Co-culture Model 3 are

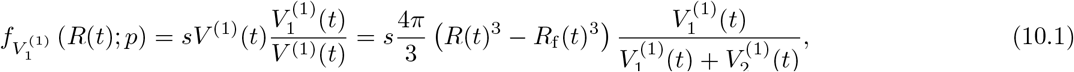

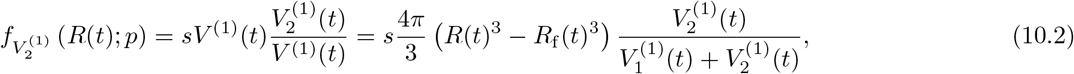

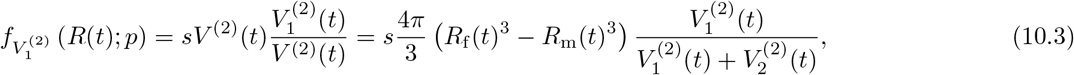

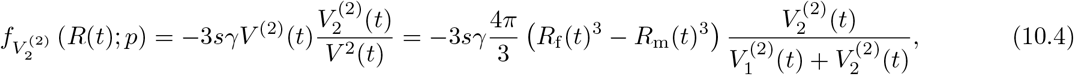

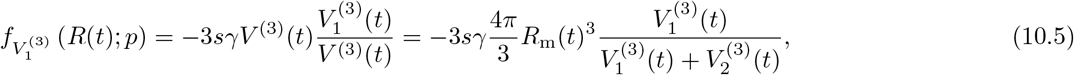

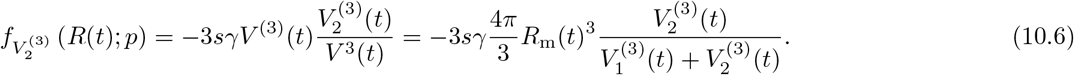

Substituting Eqs (10.1)-(10.6) into Eqs (4.1-4.4), and using the boundary constraint obtained by considering diffusion and consumption of oxygen, leads to a systems of differential equations and algebraic constraints that we solve numerically (Supplementary S3.6). This model is characterised by six parameters *θ* = (*R*(*T*), *V*_m_(*T*), *s, γ, ℛ*_m_, *ℛ*_f_).

### 4.6 Co-culture Model 4: Heterogeneous cell death oxygen thresholds and additional cell migration

To capture experimental results where fibroblasts are positioned throughout the spheroid at early times but are only observed in the central region of the spheroid at later times we extend Co-culture Model 3 (§4.5). We now include an additional cell migration mechanism that we denote *h*(*R*(*t*); *p*) ≥ 0 (Fig 3G,I). There are many biological reasons why this migration could occur and this model does not rule in or out any possibility. In general, we assume *h*(*R*(*t*); *p*) is known either from a modelling assumption or by considering additional biological mechanisms. Here, we assume that population 1 (melanoma) cells in region 2 migrate to region 1 while the same volume of population 2 (fibroblast) cells in region 1 migrate to region 2. Then we prescribe *h*(*R*(*t*); *p*) in Eq (11.7) by taking a simple approach and assuming that *h*(*R*(*t*); *p*) depends on the volume of population 2 (fibroblasts) cells in region 2, the volume of population 1 (melanoma) cells in region 1, and a new parameter *ω*. Therefore, *h*(*R*(*t*); *p*) = 0 if 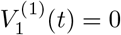 or 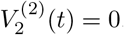. Alternative choices could be considered. Note that reversing the direction of this additional cell migration, by setting *h*(*R*(*t*); *p*) *<* 0, means that there is no mechanism for fibroblasts initially at the periphery to be lost from the periphery. Since we do not observe fibroblasts at the periphery at later times, we do not consider *h*(*R*(*t*); *p*) *<* 0. The prescribed functions for Co-culture Model 4 for *h*(*R*(*t*); *p*) ≥ 0 are

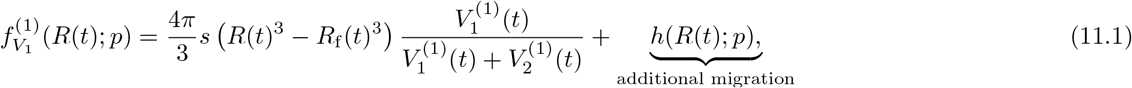

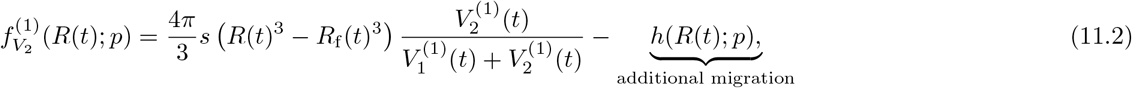

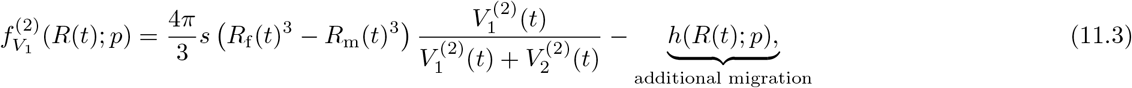

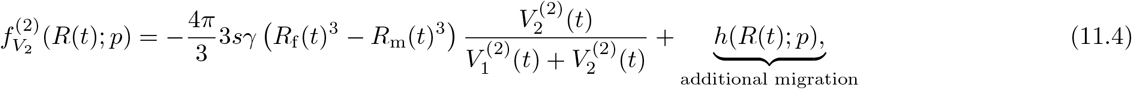

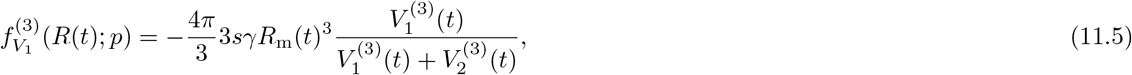

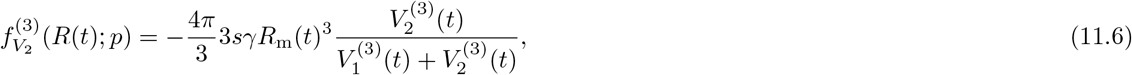

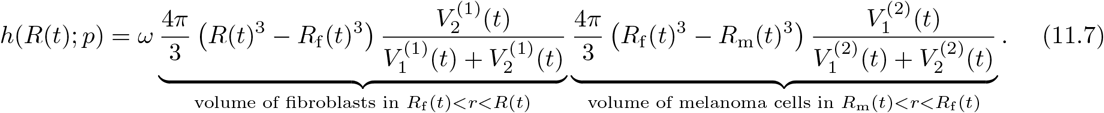

Substituting Eqs (11.1)-(11.7) into Eqs (4.1-4.4), and using the boundary constraint obtained by considering diffusion and consumption of oxygen, leads to a systems of differential equations and algebraic constraints that we solve numerically (Supplementary S3.6). This model is characterised by seven parameters *θ* = (*R*(*T*), *V*_m_(*T*), *s, γ, ℛ*_m_, *ℛ*_f_, *ω*).

## 5 Parameter estimation and identifiability analysis

For each deterministic ordinary differential equation-based mathematical model shown in §3-§4 we assess if the model parameters are practically identifiable, perform parameter estimation, and form approximate confidence intervals for model parameters using profile likelihood analysis [6, 14, 45, 46, 47, 48]. Practical identifiability assesses how well a parameter can be identified given a finite set of noisy experimental data. This is in contrast to *structural identifiability* studies that is more concerned with model structure and assesses whether parameters can be uniquely identified given a set of continuous noise-free observations [49]. For this purpose we assume that experimental measurements are noisy observations of the deterministic mathematical model. Therefore, we couple each deterministic mathematical model with a probabilistic observation model, also referred to as an error model. The probabilistic observation model accounts for experimental variability and measurement error. We assume that observation errors are independent, identically distributed, additive and normally distributed with zero mean and constant variance *σ*^2^. We estimate *σ* alongside other model parameters. For the biphasic model (Eq (1)) and linear model (Eq (2)) experimental measurements comprise of estimates of the radius of each spheroid, *R*(*t*). For the compartment-based models in §4 experimental measurements comprise of estimates of the radius of each spheroid, *R*(*t*), and the necrotic core of each spheroid, *R*_n_(*t*).

For this approach we use the log-likelihood function 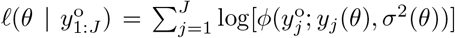, where *ϕ*(*x*; *μ, σ*^2^) denotes a Gaussian probability density function with mean *μ* and constant variance *σ*^2^; *θ* denotes model parameters; and 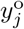 denotes the *j*^th^ experimental observation. For a best-fit to the data we compute the maximum likelihood estimate (MLE), 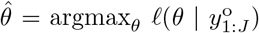, subject to bound constraints. Results are reported in terms of the normalised log-likelihood function, 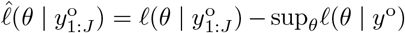. Note 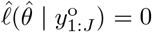. To compute profile likelihoods we assume that the full parameter *θ* can be partitioned into *θ* = (*ψ, λ*), where *ψ* is a scalar interest parameter and *λ* is a vector nuisance parameter. The profile likelihood for *ψ* is 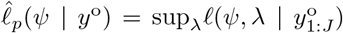. Bounds are chosen to ensure that approximate 95% confidence intervals, defined using a profile likelihood threshold value of −1.92 [50], are captured. Mathematical models are solved numerically using the open-source DifferentialEquations package in Julia (Supplementary S4). Numerical optimisations are performed using the Nelder–Mead routine in the open-source NLopt optimisation package in Julia [51], for further detail see [14].

## 6 Results and discussion

Here we analyse spheroid formation, geometry, and growth of overall size and structure in co-culture experiments. We seek to provide mechanistic insight by analysing experimental measurements of *R*(*t*) and *R*_n_(*t*) within objective mathematical and statistical modelling framework presented in §3-§5.

### 6.1 Spheroid formation and geometry

Spheroid experiments are routinely performed to characterise cancer growth and fibroblasts are commonly included in cancer cell line experiments to enhance spheroid formation and structure. To understand this approach, compare the size and densities of 1205Lu monoculture (M100:F0) spheroids and monoculture fibroblast (M0:F100) spheroids (Fig 5A). In both conditions cells initially placed in the well migrate and adhere. The M0:F100 spheroid forms a compact, dense, and approximately spherical structure by approximately 48 hours. In contrast, the M100:F0 spheroid does not (Fig 5B). Experimental images for intermediate conditions, namely M75:F25, M50:F50, and M25:F75, show that fibroblasts enhance formation (Fig 5A,C). We are interested in the contribution of fibroblasts to melanoma spheroids so we do not consider the M0:F100 condition in the remainder of the study.

**Figure 5:**
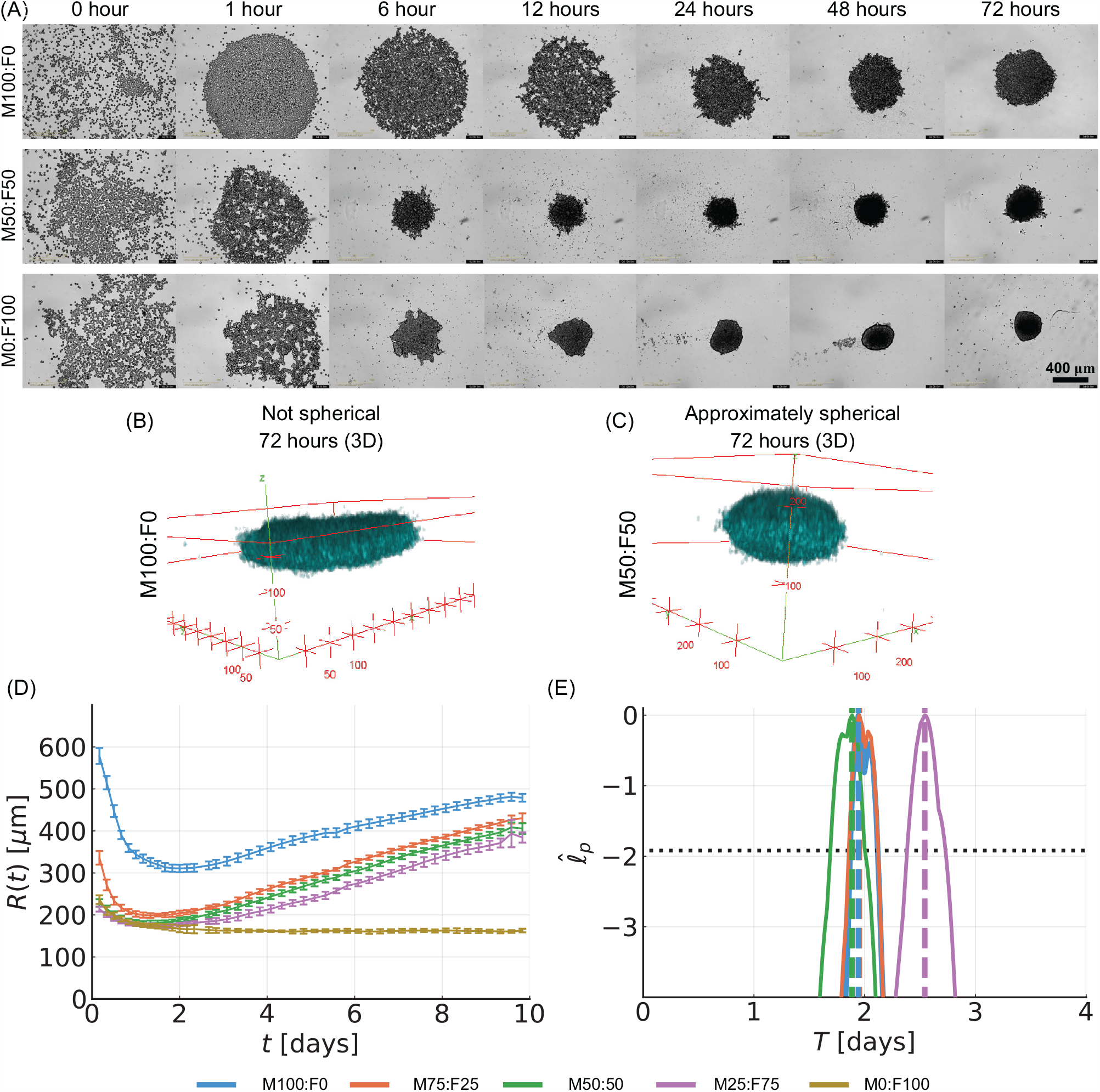
Spheroid formation analysis. Results shown for co-culture spheroids formed with the 1205Lu cell line and fibroblasts. (A) Brightfield experimental images captured from above each spheroid at 0, 1, 6, 24, 48, and 72 [hours]. (B,C) 3D rendering from confocal microscopy z-stack images at 72 hours for (B) M100:F0 and (C) M50:F50 spheroids. The spheroid in (B) is relatively flat while it is reasonable to approximate the spheroid in (C) as spherical. Scale in (B,C) is microns. Colour represents a cell nuclear stain (DAPI) for melanoma cells and fibroblasts. (D) Temporal evolution of measurements of spheroid size, *R*(*t*) [μm]. Results shown for sixteen spheroids for the M100:F0 condition, eight spheroids for the M75:F25 and M25:F75 conditions and seven spheroids for the M50:F50 and M0:F100 conditions. Error bars represent standard deviation about the mean at each time point. (E) Profile likelihoods for formation time, *T* [days], with approximate 95% confidence interval threshold (horizontal black-dashed). Throughout M100:F0 (blue), M75:F25 (orange), M50:F50 (green), M25:F75 (magenta), and M0:F100 (yellow).

We now explore the quantitative impact of fibroblasts on melanoma spheroid formation time, *T* [days], using the biphasic model (§3.1) and profile likelihood analysis (§5) to interpret the experimental measurements of spheroid size, *R*(*t*) (Fig 5D). For each condition, profile likelihoods for *T* are well-formed and relatively narrow about a single central peak suggesting *T* is practically identifiable. Three out of the four profiles appear to be consistent suggesting that spheroid formation time relative to the experimental duration is not drastically altered across the conditions (Fig 5E, Table 2). It is not clear if the profile for M25:F75 is different for a biologically meaningful reason or due to experimental noise (Fig 5E). In Supplementary S5.2 we show that, for each condition, experimental data are accurately described by simulating the biphasic mathematical model with the MLE and that the other parameters in the biphasic mathematical model are practically identifiable. In the remainder of this study we focus on the growth of WM983B co-culture spheroids because they form dense, compact, and approximately spherical spheroids across all conditions (M100:F0, M75:F25, M50:F50, M25:F75).

**Table 2:** Estimates of spheroid formation time, *T* [days] for 1205Lu co-culture spheroids. Results show MLE and 95% approximate confidence intervals obtained from the profile likelihoods presented in Fig 5E. We consider four conditions that correspond to the initial proportions of melanoma cells and fibroblasts at seeding.

| Condition | Formation time, $T$ [days] |
| --- | --- |
| M100:F0 | 1.94 (1.87, 2.12) |
| M75:F25 | 1.95 (1.86, 2.12) |
| M50:F50 | 1.89 (1.69, 2.01) |
| M25:F75 | 2.54 (2.38, 2.73) |

### 6.2 Temporal evolution of spheroid size

The temporal evolution of spheroid size, *R*(*t*), is approximately linear in all conditions (Fig 6B). Therefore, we analyse the data using a linear model (Eq (2). Profile likelihoods suggest that *R*(3) and *λ* are practically identifiable since the univariate profiles are each well-formed around a single peak (Figs 6C-D). Profile likelihoods for *R*(3) capture that spheroids seeded with a greater proportion of melanoma cells initially form larger spheroids (Fig 6C, Table 3). Profile likelihoods for *λ* suggest the growth rates are relatively consistent across conditions (Fig 6D, Table 3). Previous monoculture results suggest that increasing the experimental duration results in sigmoidal growth [5, 6, 14].

**Table 3:** Estimates of linear model parameters for WM983B co-culture spheroids. Parameters reported include the initial spheroid radius, *R*(3) [μm], and the growth rate, *λ* [day^−1^], obtained from the profile likelihoods presented in Fig 6C,D, respectively. Results show MLE and 95% approximate confidence intervals.

| Condition | Initial Radius, $R(3)$ [$\mu\text{m}$] | Growth rate, $\lambda$ [$\text{day}^{-1}$] |
| --- | --- | --- |
| M100:F0 | 181.69 (181.64, 181.82) | 0.0913 (0.0909, 0.0920) |
| M75:F25 | 174.47 (174.26, 174.80) | 0.0878 (0.0866, 0.0891) |
| M50:F50 | 172.46 (172.28, 172.70) | 0.0776 (0.0765, 0.0786) |
| M25:F75 | 158.38 (157.76, 158.66) | 0.0796 (0.0780, 0.0812) |

**Figure 6:**
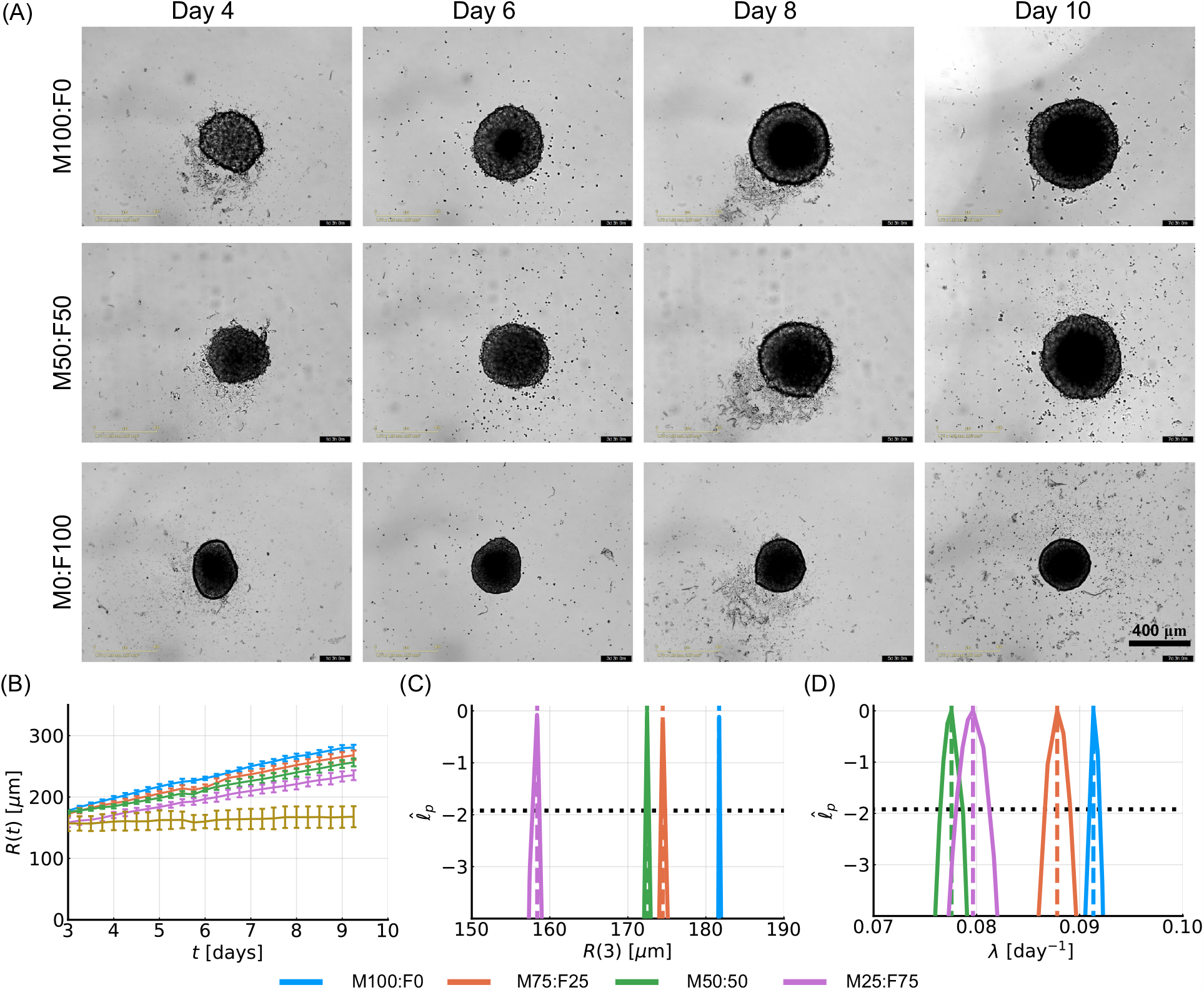
Temporal evolution of spheroid structure. Results shown for co-culture spheroid experiments formed with the WM983B cell line and fibroblasts. (A) Brightfield experimental images captured from above each spheroid. Exemplar image masks shown in Figure S1. (B) Temporal evolution of overall spheroid size *R*(*t*) [μm]. Results shown for 15, 7, 8, 7, and 12 spheroids for the M100:F0, M25:F75, M50:F50, M25:F75, and M0:F100 conditions. (C) Profile likelihoods for initial spheroid size, *R*(3) [μm].(D) Profile likelihoods for growth rate, *λ* [day^−1^]. In (C,D) approximate 95% confidence interval threshold shown with horizontal black-dashed line. Throughout conditions are M100:F0 (blue), M75:F25 (orange), M50:F50 (green), M25:F75 (magenta), and M0:F100 (yellow).

### 6.3 Temporal evolution of spheroid size and structure

Here we explore the role of fibroblasts play in influencing the size and structure of the co-culture spheroids. We analyse confocal microscopy images of each spheroid’s equatorial plane, measure *R*(*t*) and *R*_n_(*t*), and interpret these data using a series of mathematical models, parameter estimation, and profile likelihood analysis.

To identify living and proliferating regions within each spheroid we use FUCCI signals. For M100:F0 spheroids, on days 2 and 3 magenta and green FUCCI signals are present throughout the spheroid indicating all cells are living and proliferating (Fig 7A). From day 6 onwards the large central region without FUCCI signals indicates a necrotic core, previously confirmed by both confocal microscopy and flow cytometry using cell death markers [6, 13, 52, 53]. Outside of the necrotic core FUCCI signals indicate a proliferating region at the periphery (predominately green) and an intermediate region of living cells that are proliferation-inhibited (predominately magenta). We observe similar behaviour, albeit the necrotic core forms later, for M75:F25, M50:F50, and M25:F75 spheroids (Fig 7). Using standard image processing techniques we measure *R*(*t*) and *R*_n_(*t*) (Figs 8A-D, Supplementary S1 and S2.2). We focus on *R*(*t*) and *R*_n_(*t*) to present results in terms of typical measurements and this means that we group together the proliferating and proliferation-inhibited regions [6].

**Figure 7:**
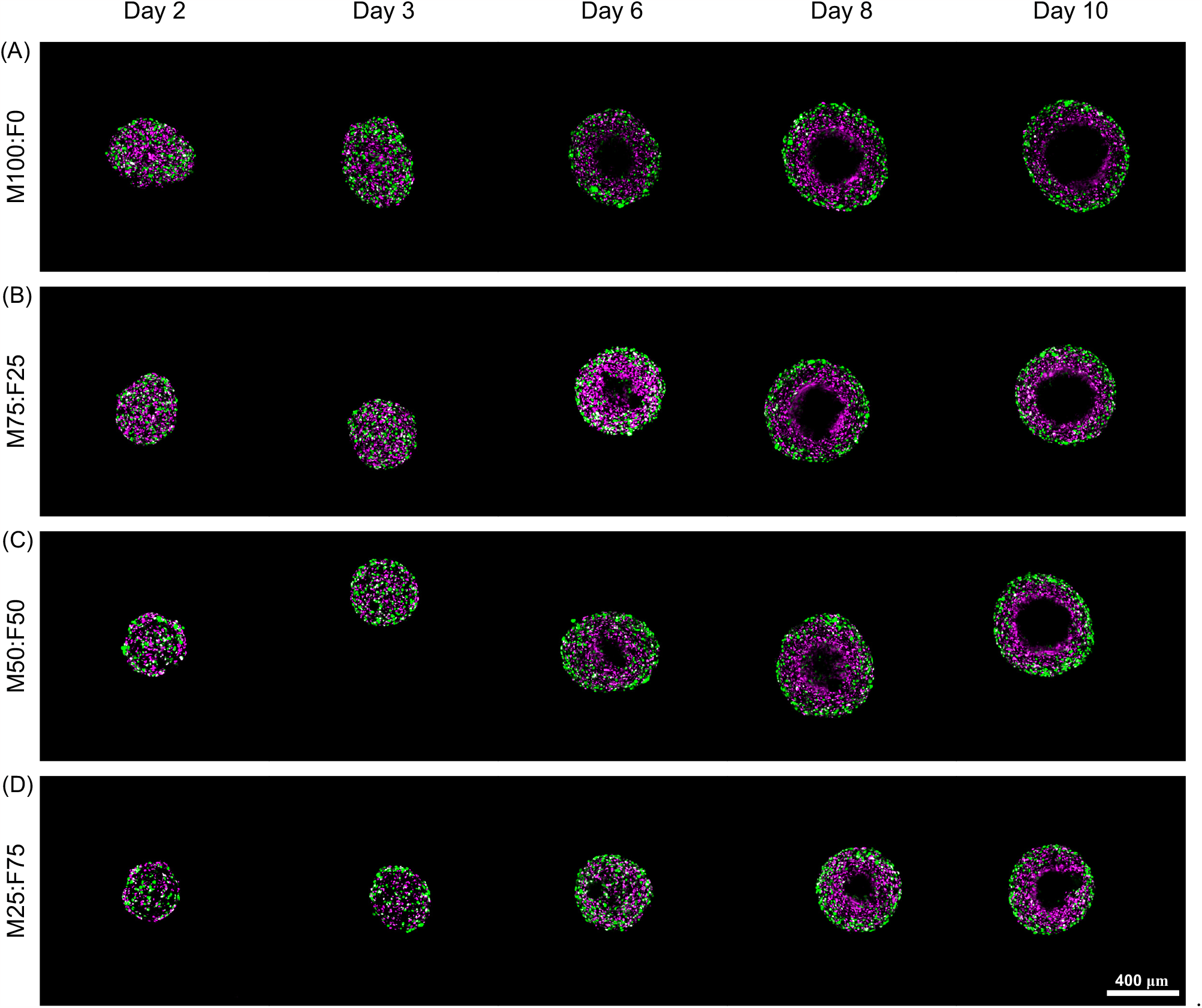
Spatio-temporal evolution of tumour spheroid structure. Confocal microscopy images of each spheroid’s equatorial plane for (A) M100:F0, (B) M75:F25, (C) M50:F50, and (D) M25:F75 spheroids. Results shown for the WM983B melanoma cell line. FUCCI signals are shown with magenta and green. Presence of FUCCI signal indicates living cells whereas large regions at the centre of spheroids that lack FUCCI signals indicate a necrotic core. Fibroblasts are not shown so that it is easier to visualise the necrotic core and overall size.

**Figure 8:**
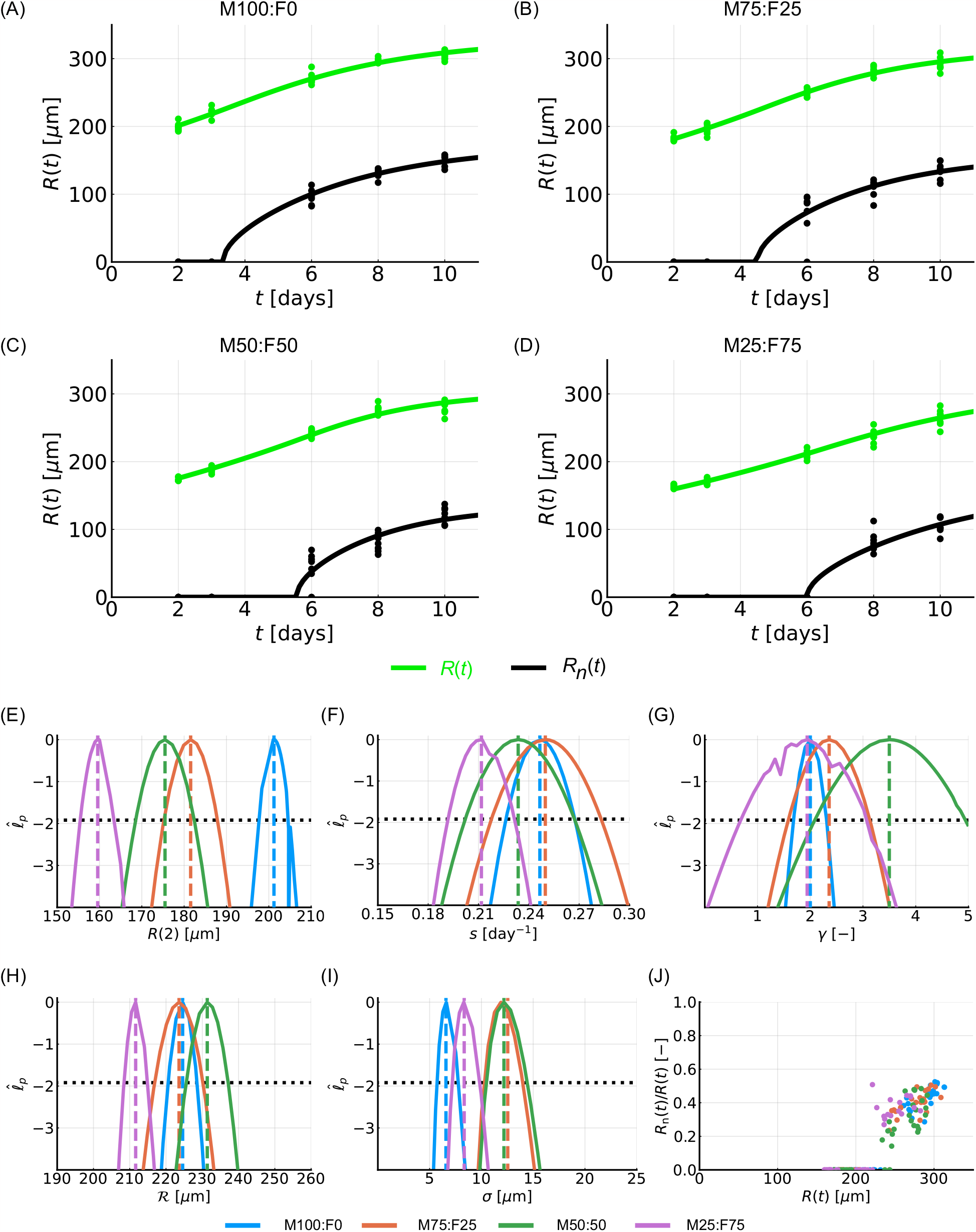
Temporal evolution of spheroid size and structure analysed using the reduced Greenspan mathematical model. (A-D) Comparison of experimental data and mathematical model simulated with the maximum likelihood estimate for: (A) M100:F0; (B) M75:F25; (V) M50:F50; (D) M25:F75 spheroids. In (A-D) experimental measurements of spheroid radius, *R*(*t*) [μm], and necrotic radius, *R*_n_(*t*) [μm], shown as green and black circles, respectively. (E-H) Profile likelihoods for (E) *R*(*T*) [μm]; (F) *s* [day^−1^]; (G) *γ* [-]; (H) *R* [μm], and (I) *σ* [μm]. In (E-I) approximate 95% confidence interval threshold shown with horizontal black-dashed line. (J) Measurements of the necrotic fraction, *R*_n_(*t*)*/R*(*t*) [-], against spheroid radius, *R*(*t*) [μm]. Throughout conditions are M100:F0 (blue), M75:F25 (orange), M50:F50 (green), and M25:F75 (magenta).

Seeking to characterise co-culture spheroid growth, we again turn to mathematical modelling and profile likelihood analysis. We consider a reduced version of Greenspan’s mathematical model [8], introduced by Burton [19], and focus on the temporal evolution of *R*(*t*) and *R*_n_(*t*) (§4.2). This model is relatively simple. Further, all mechanisms and parameters in the model are biologically interpretable. This is a great advantage over more complicated models where the biological meaning of parameters and mechanisms is not always clear. Note that this mathematical model assumes a monoculture spheroid whereas the experimental data comprise of measurements of co-culture spheroids. We take this approach deliberately. This approach allows us to explore whether estimates of parameters that characterise growth vary across experimental conditions. For each experimental condition, the monoculture model simulated with the MLE agrees with measurements of *R*(*t*) and *R*_n_(*t*) (Figs 8A-D). Profile likelihoods suggest that all model parameters are practically identifiable since the univariate profiles are each well-formed around a single peak (Figs 8E-I, Table 4). The profile for *R*(2) [μm] suggests the experimental condition influences the initial size of each spheroid (Fig 8E, Table 4). Profiles for the proliferation rate per unit volume of living cells, *s* [day^−1^], the dimensionless parameter related to the loss rate from the necrotic core, *γ* [-], and size of the spheroid when the necrotic region forms, *R* [μm], appear to be consistent across all conditions (Fig 8F,G,H, Table 4). Profiles for the standard deviation (Fig 8I, Table 4) capture variability in the experimental data (Fig 8A-D). These powerful quantitative insights suggest that growth characteristics are similar across conditions. Spheroid structure, assessed by measuring the necrotic fraction, *R*_n_(*t*)*/R*(*t*) [-], also appears to be consistent across conditions (Fig 8J). However, these results do not take into account the internal dynamics of the two populations within the growing spheroid.

**Table 4:** Estimates of monoculture reduced Greenspan model parameters for WM983B coculture spheroids. Parameters reported include *R*(2) [μm], *s* [day^−1^], *γ* [-], *R* [μm], and *σ* [μm]. Results show MLE and 95% approximate confidence intervals.

| Condition | $R(2)$ [$\mu\text{m}$] | $s$ [$\text{day}^{-1}$] | $\gamma$ [-] | $\mathcal{R}$ [$\mu\text{m}$] | $\sigma$ [$\mu\text{m}$] |
| --- | --- | --- | --- | --- | --- |
| M100:F0 | 201.2 (197.7, 205.1) | 0.247 (0.226, 0.267) | 2.00 (1.69, 2.31) | 224.6 (220.6, 228.5) | 7.5 (5.7,7.8) |
| M75:F25 | 181.5 (175.3, 187.9) | 0.250 (0.218, 0.283) | 2.36 (1.59, 3.13) | 223.6 (216.9, 230.2) | 12.5 (10.3,14.0) |
| M50:F50 | 175.5 (168.5, 182.4) | 0.234 (0.202, 0.267) | 3.50 (2.09, 4.95) | 231.4 (225.6, 237.2) | 12.2 (10.5,14.4) |
| M25:F75 | 159.6 (155.5, 163.4) | 0.212 (0.192, 0.231) | 1.94 (0.68, 3.07) | 211.6 (208.4, 214.9) | 9.1 (7.2,9.8) |

Thus far we have examined the experimental images using FUCCI signals to identify regions of living melanoma cells, that are proliferating or proliferation-inhibited, and necrotic matter. Here, we re-examine these experimental images and include the signal from the fibroblast marker (Fig 9, Supplementary S2.2). Focusing on the M75:F25, M50:F50, and M25:F75 spheroids, fibroblasts are present throughout each spheroid on days 2 and 3 (Fig 9B,C,D). At later times we only observe fibroblasts close to the centre of the spheroid. Therefore, assumptions that the two cell types are well mixed within compartments appears reasonable. The biological mechanisms driving the changing position of the fibroblasts within the spheroid are unclear. Seeking further insights we again turn to mathematical modelling and systematically extend the monoculture reduced Greenspan model.

**Figure 9:**
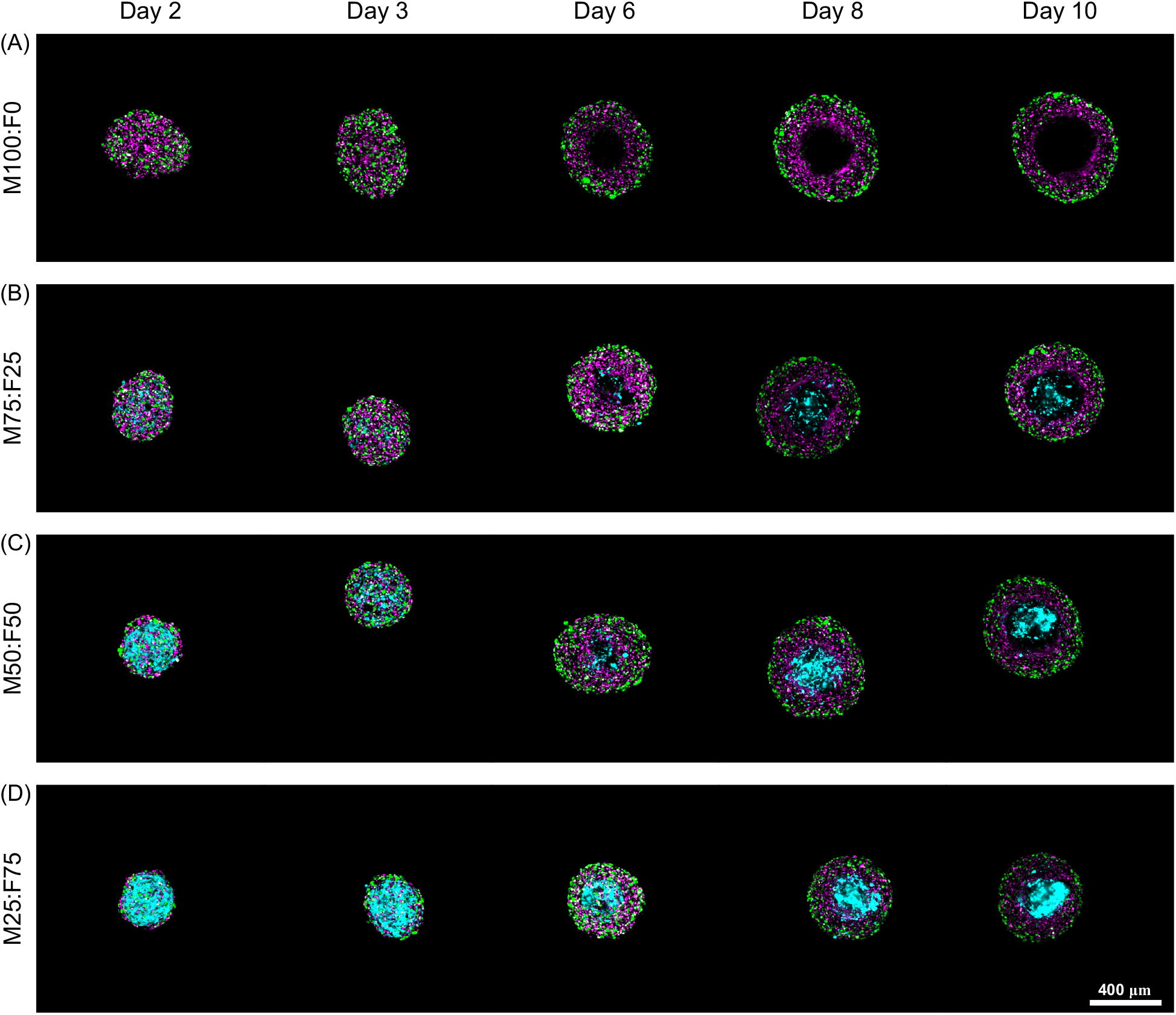
Spatio-temporal evolution of tumour spheroid structure including fibroblast marker. Confocal microscopy images of each spheroid’s equatorial plane for (A) M100:F0, (B) M75:F25, (C) M50:F50, and (D) M25:F75 spheroids. These are same spheroids as shown in Fig (7) now with the fibroblast marker shown in cyan. The fibroblast marker stains the entire cell whereas the FUCCI signal is only present at the cell nucleus.

We systematically extend the monoculture model by introducing heterogeneity independently and in turn for the proliferation rate *s*, the loss rate *γ*, and the cell death oxygen thresholds via *ℛ*. This results in four new co-culture models: (i) heterogeneity in *s* (§4.3); (ii) heterogeneity in *γ* (§4.4); (iii) heterogeneity in ℛ (§4.5); and (iv) heterogeneity in ℛ and an additional cell migration mechanism (§4.6). Parameter values are chosen to capture, where possible, loss of fibroblasts from the periphery and chosen so that model simulations agree with measurements of *R*(*t*) and *R*_*n*_(*t*) from the M50:F50 condition. Specifically, we simulated each model at a local estimate of the MLE, where the initial estimate of the MLE for the MLE search was chosen based on good qualitative agreement between the solution of the mathematical model and the data. Results for the M75:F25 and M25:F75 conditions are similar to those discussed in the following.

In Co-culture Model 1 (§4.3) we assume that melanoma cells and fibroblasts proliferate at different rates per unit volume, *s*_m_ [day^−1^] and *s*_f_ [day^−1^], respectively (Fig 1I). Simulating Model 1 with *s*_m_ *> s*_f_ we can capture the temporal evolution of measurements of *R*(*t*) and *R*_n_(*t*) (Fig 10A). For greater insight, we analyse the temporal evolution of the volume of melanoma cells and fibroblasts within each compartment *j* = 1, 2 of the spheroid, denoted 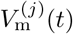 and 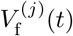, respectively. Here compartment one represents the periphery, *R*_n_(*t*) *< r < R*(*t*), and compartment two represents the necrotic core, 0 *< r < R*_n_(*t*). For each compartment the melanoma fraction is 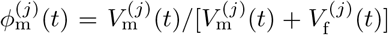. Since 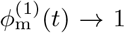as *t → ∞*, Co-culture Model 1 captures the loss of fibroblasts from the periphery (Fig 10B). Further, since 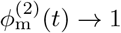as *t → ∞*, Co-culture Model 1 suggests that the necrotic core will eventually be composed only of matter from necrotic melanoma cells (Fig 10B). This is a useful insight. In experimental images we are unable to quantify the proportion of necrotic matter that is from melanoma cells and fibroblasts due to loss of signal during necrosis.

**Figure 10:**
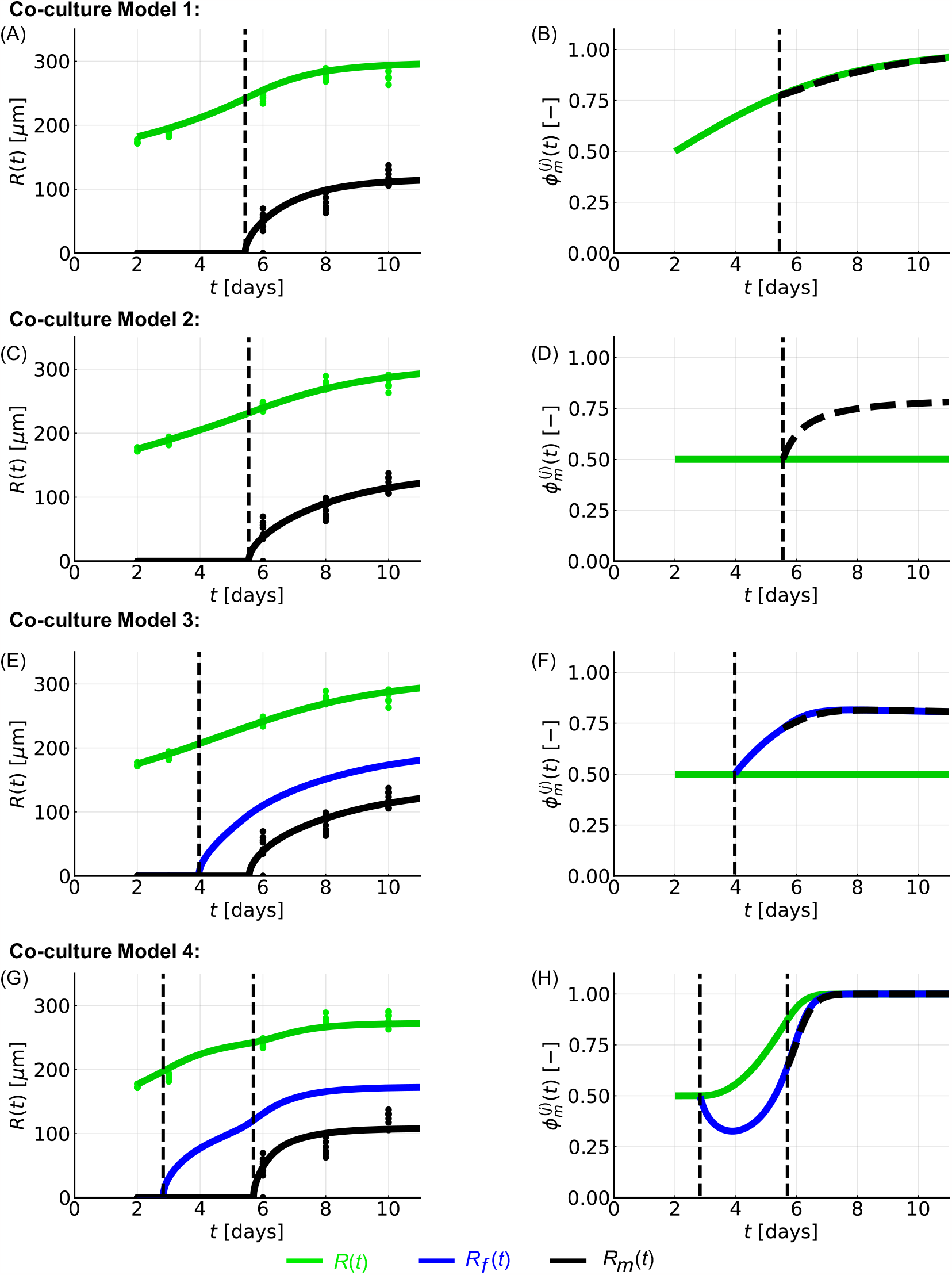
Spatio-temporal evolution of spheroid structure analysed using multiple co-culture reduced Greenspan models. (A,C,E,G) Simulations of (A) Co-culture Model 1, (C) Co-culture Model 2, (E) Co-culture Model 3, (G) Co-culture Model 4 show the temporal evolution of spheroid size *R*(*t*) (green). For Co-culture Models 1 and 2 we present the size of necrotic core *R*_n_(*t*) (black). For Co-culture Models 3 and 4 we present *R*_f_ (*t*) (blue) and *R*_m_(*t*) (black). Experimental measurements of *R*(*t*) and necrotic core from M50:F50 spheroids are shown as green and black circles, respectively. (B,D,F,H) Temporal evolution of melanoma fraction 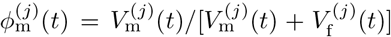 (green), 2 (black dashed) in Co-culture Models 1 and 2, and *j* = 1 (green), 2 (black dashed), 3 (blue) in Models 3 and 4. Parameter values and short descriptions are listed in Table 5.

In Co-culture Model 2 (§4.4), we assume that the rate at which mass in the necrotic core degrades and is lost from the spheroid depends on whether the necrotic matter is from melanoma cells or fibroblasts (Fig 1J). This is associated with two parameters *γ*_m_ [-] and *γ*_f_ [-]. Simulating Co-culture Model 2 we can capture the temporal evolution of measurements of *R*(*t*) and *R*_n_(*t*) but we cannot capture the loss of fibroblasts at the periphery (Fig 10C,D).

**Table 5:** Estimates of co-culture model parameters for WM983B M50:F50 co-culture spheroids. Results show a local estimate of the MLE, where initial guesses for the MLE search were chosen based on good qualitative agreement between the solution of the mathematical model and the data. Note that *V* (*T*) = 4*πR*(*T*)^3^*/*3.

| Parameter | Units | Short description | Co-culture<br>Model 1 | Co-culture<br>Model 2 | Co-culture<br>Model 3 | Co-culture<br>Model 4 |
| --- | --- | --- | --- | --- | --- | --- |
| $R(T)$ | $\mu\text{m}$ | Spheroid radius at formation | 177.00 | 175.35 | 176.00 | 177.00 |
| $V_m(T)$ | $\mu\text{m}^3$ | Volume of melanoma cells at formation | $0.50V(T)$ | $0.50V(T)$ | $0.50V(T)$ | $0.50V(T)$ |
| $s_m$ | $\text{day}^{-1}$ | Melanoma proliferation rate | 0.35 | 0.24 | 0.24 | 0.40 |
| $s_f$ | $\text{day}^{-1}$ | Fibroblast proliferation rate | 0.10 | 0.24 | 0.24 | 0.40 |
| $\gamma_m$ | - | Loss rate of melanoma cells from<br>necrotic core | 3.07 | 2.07 | 2.06 | 5.00 |
| $\gamma_f$ | - | Loss rate of fibroblasts from necrotic<br>core | 3.07 | 8.00 | 2.06 | 5.00 |
| $\mathcal{R}_m$ | $\mu\text{m}$ | Melanoma spheroid radius when<br>necrotic region forms. Associated with<br>oxygen threshold for melanoma cell<br>death, $p_m$ [%]. | 235.00 | 231.44 | 207.07 | 197.50 |
| $\mathcal{R}_f$ | $\mu\text{m}$ | Fibroblast spheroid radius when<br>necrotic region forms. Associated with<br>oxygen threshold for fibroblast cell<br>death, $p_f$ [%]. | 235.00 | 231.44 | 220.94 | 220.00 |
| $\omega$ | $\text{day}^{-1} \mu\text{m}^{-3}$ | Associated with rate of additional cell<br>migration | - | - | - | $9.75 \times 10^{-7}$ |

In Co-culture Models 3 (§4.5) and 4 (§4.6), we generalise the single oxygen partial pressure threshold, *p*_n_ [%], with two thresholds, *p*_f_ ≥ *p*_m_ [%], corresponding to the oxygen partial pressure below which fibroblasts and melanoma cells die, respectively (Fig 1K,L). These two thresholds define *R*_f_ (*t*) and *R*_m_(*t*), respectively, that satisfy *R*_f_ (*t*) ≥ *R*_m_(*t*) (Fig 1N), and two parameters (*R*_f_, *R*_m_). This gives rise to a model with three compartments (Fig 1I-N). Here we assume that measurements of the necrotic core correspond to measurements of *R*_m_(*t*) that is defined by *p*_m_. This is because we measure the size of the necrotic region using FUCCI signals from melanoma cells. Similarly to Co-culture Model 2, with Co-culture Model 3 we can capture the temporal evolution of measurements of *R*(*t*) and *R*_m_(*t*) (Fig 10E) but cannot capture the loss of fibroblasts at the periphery (Fig 10F).

Introducing additional cell migration into Co-culture Model 3 gives Co-culture Model 4. We assume that living fibroblasts at the periphery, *R*_f_ (*t*) ≤ *r* ≤ *R*(*t*), migrate towards the centre of the spheroid, *R*_m_(*t*) ≤ *r* ≤ *R*_f_ (*t*), while the same volume of melanoma cells migrate in the opposite direction (Fig 1N). This can give rise to transient dynamics where the proportion of fibroblasts at the core increases followed by long-term loss of fibroblasts throughout the spheroid (Fig 10G,H). There are many biological reasons why this migration could occur and this model does not rule in or out any possibility. Experimental observations for co-culture spheroids with the 1205Lu cell line suggest that fibroblasts can form the core of spheroids.

## 7 Conclusion

In this study we develop a general mathematical and statistical framework to characterise and interpret co-culture spheroid formation and growth. We perform co-culture spheroid experiments and find that introducing fibroblasts allows a cancer cell line to generate spheroids with approximately spherical structures that would not be possible otherwise. Using our mathematical and statistical modelling framework we then quantify spheroid formation time and estimate key parameters that characterise growth. We find that incorporating fibroblasts has limited impact on formation time, spheroid growth and structure. There are many possible reasons for these results. For example it could be that there are limited interactions, or cross-talk, between the melanoma cells and fibroblasts. As part of this analysis we quantitatively directly connect a version Greenspan’s seminal model for monoculture spheroids to co-culture spheroid data for the first time. We choose to use a monoculture model to describe co-culture data deliberately. This approach suggests that parameter estimates and growth characteristics are consistent between WM983b melanoma spheroids without fibroblasts and melanoma spheroids in co-culture with fibroblasts at initial percentages ranging from 25% to 75%. However, the internal dynamics of growing spheroids can be more complicated and these dynamics should be taken into account when designing experiments and interpreting measurements obtained from those experiments. By extending Greenspan’s model and a general class of monoculture compartment-based mathematical models to multiple populations, we reveal biological mechanisms that can describe the internal dynamics of co-culture spheroids and those that cannot.

Our mathematical and statistical modelling framework can be applied to different cell types and to spheroids grown in different conditions. Here we choose to perform experiments with melanoma cells and fibroblasts. This builds on our previous monoculture melanoma spheroid studies [5, 6, 14, 13]. Our results are consistent with analysis of two-dimensional experiments which suggest that fibroblasts have a limited impact on co-culture melanoma spreading and invasion [54]. However, we note that other spheroid studies suggest fibroblasts contribute to melanoma spheroid growth [12]. Interesting future work would be to explore these differences. Such differences could arise due to many reasons, for example choice of cell line, fibroblasts, and experimental conditions. Further, the framework is well-suited to diagnose cross-talk in other co-culture experiments.

In our WM983B co-culture experiments fibroblasts are present throughout the spheroid at early times but are only present in the central region of the spheroid at later times, also observed in [11, 55]. Using our new co-culture compartment-based mathematical models we identify that differences in proliferation rates (Co-culture Model 1), or differences in when each cell type undergoes necrosis due to lack of oxygen in combination with cell migration (Co-culture Model 4), could drive this behaviour. However, differences in loss rates from the necrotic core (Co-culture Model 2) and differences in when each cell type undergoes necrosis due to lack of oxygen (Co-culture Model 3) are on their own insufficient to capture this behaviour. We would not have been able to explore these biological differences without these new models and experimental data. Performing additional experiments to inhibit proliferation, or growing co-culture spheroids in different oxygen conditions [14], would be interesting and may provide useful insights to improve understanding of the mechanisms driving internal dynamics of growing co-culture spheroids. In general, additional data would be helpful to assess whether parameter identifiability in the co-culture mathematical models. Our framework is well-suited to incorporate such additional data and, if appropriate, different error models [45, 48].

The co-culture mathematical modelling framework is general. We use this framework to extend the monoculture reduced Greenspan mathematical model to two populations. This allows us to explore the temporal evolution of co-culture spheroid size and structure. In Supplementary S5.1 we present further examples of the general model, including the seminal Greenspan model and the radial-death model [17]. The radial-death model captures key features of Greenspan’s model, but rather than considering oxygen mechanisms prescribes a fixed size for the proliferating region. Many other models could be considered with additional compartments and/or mechanisms such as nutrient/waste diffusion and consumption/production [19, 20, 21, 22, 23, 24, 25, 26, 27, 28, 29, 30, 31]. For example, we could use our experimental images to identify regions where melanoma cells are proliferation-inhibited and incorporate this into mathematical modelling [5, 6, 8, 14]. As a further example, we could relax the assumption of a single compartment at formation and explore co-culture spheroid data where one cell type is located at the core of the spheroid throughout (Supplementary S.5.4). In general, the additional complexity of these models may result in parameter identifiability challenges with currently available data [17, 46]. Incorporating mechanical effects [56, 57] and extracellular material as an additional population are also of interest. The model can also be adapted to consider cell types with different densities, or compartments with different densities, by considering the total cell number as well as the volume. The framework could also be extended to characterise the effectiveness of chemotherapies and radiotherapies [9, 10, 11, 12, 58, 59]. If compartment-based modelling assumptions, such as assumptions of spherical symmetry or well-mixed compartments, break down then alternative models should be considered [32, 33, 34, 35]. Throughout the study we have focused on parameter estimation. It would be interesting future work to explore the predictive capability of the models, for example by using profile likelihood-based prediction tools [47, 48]. Overall, this study presents a quantitative approach to characterise the formation of co-culture spheroids and the internal dynamics of growing co-culture spheroids.

## Code and data availability

Key computer code used to generate computational results and datasets generated and analysed during this study are summarised in the electronic supplementary material and are available on a GitHub repository (https://github.com/ryanmurphy42/Murphy2022CoCulture).

## Author’s contributions

All authors conceived and designed the study. RJM performed the mathematical and statistical modelling, and drafted the article. GG performed all experimental work. All authors provided comments and approved the final version of the manuscript. NKH and MJS contributed equally.

## Competing interests

We declare we have no competing interest.

## Funding

MJS and NKH are supported by the Australian Research Council (DP200100177).

## Acknowledgements

We thank Associate Professor Rick Sturm, Frazer Institute (University of Queensland) for providing the fibroblasts. We thank John Blake for guidance using IncuCyte. This research was carried out at the Translational Research Institute (TRI), Woolloongabba, QLD. TRI is supported by a grant from the Australian Government. We thank the staff in the microscopy core facility at TRI for their technical support. We thank Professor Atsushi Miyawaki, RIKEN, Wako-city, Japan, for providing the FUCCI constructs, and Professor Meenhard Herlyn, The Wistar Institute, Philadelphia, PA, for providing the cell lines.

## Notes

https://github.com/ryanmurphy42/Murphy2022CoCulture

### Competing Interest Statement

The authors have declared no competing interest.

### Summary of Updates

Changes have been made to the presentation of the manuscript, results are unchanged. Specifically, Fig 1 has been replaced with Fig 1, 2 and 3 describing workflow, and mathematical model schematics. Details of experimental methods and additional mathematical modelling equations have been moved to supplementary.

