## Supplementary Material for "Formation and growth of co-culture tumour spheroids: new compartment-based mathematical models and experiments"

**S1 Experimental methods** **3**

### S1 Experimental methods

We perform two co-culture experiments (Fig 1). Experiment 1 focuses on spheroid formation and growth of overall spheroid size. Experiment 2 focuses on the size, internal structure, and geometry of growing spheroids after formation. In the following we detail the experimental methods for cell culture; spheroid generation, culture, and experiments; imaging and image processing.

#### S1.1 Cell culture

Methods for cell culture are identical to that described in our previous tumour spheroid studies [1, 2]. For clarity we outline the method here. The human melanoma cell lines WM983B and 1205Lu were provided by Professor Meenhard Herlyn, The Wistar Institute, Philadelphia, PA, [3], previously genotypically characterised [4, 5, 6, 7], grown as described in [8], and previously authenticated by short tandem repeat fingerprinting (QIMR Berghofer Medical Research Institute, Herston, Australia). Both melanoma cell lines were previously transduced with fluorescent ubiquitination-based cell cycle indicator (FUCCI) constructs as described in [5, 8]. Using FUCCI-transduced cell lines allows us to visualise the cell-cycle status of each melanoma cell continuously throughout time without loss of signal [5, 9]. Human primary fibroblast QF1696 were a gift from Associate Professor Rick Sturm (Frazer Institute, University of Queensland) [10].

#### S1.2 Spheroid generation, culture, and experiments.

Spheroids were generated in 96-well cell culture flat-bottomed plates (3599, Corning), with a total seeding density of 5000 total cells/well, using 50 µL total/well non-adherent 1.5% agarose to promote formation of a single spheroid per well [1, 5, 8, 11]. For each experiment, we mix one human melanoma cell line (WM983B or 1205Lu) with human primary fibroblasts (QF1696) at five different initial compositions (M100:F0, M75:F25, M50:F50, M25:F75, M0:F100, where M$x$:F$y$ is the initial proportion, measured as a percentage, of melanoma cells $x$ [%] and fibroblasts $y = 100 - x$ [%]). Fibroblasts are included to enhance spheroid formation. We do not mix the two melanoma cell lines. Experiments 1 and 2 are performed for ten days after seeding, replacing 50% of the medium in each well (200 µL total/well) on days 3, 5, 7, and 9.

In Experiment 1, for each melanoma cell line, we seed sixteen spheroids for the M100:F0 condition, and eight spheroids for the M75:F25, M50:F50, M25:F75, and M0:F100 conditions into a plate (§S1.2). The plate is placed inside the IncuCyte S3 live cell imaging incubator (37 °C, 5% $CO_2$) immediately after seeding to the end of the experiment.

For Experiment 2 fibroblasts were stained with CellTracker™ Deep Red dye as per manufacturer's instructions prior to seeding. The fibroblast marker stains the entire cell whereas the FUCCI signal is only present at the cell nucleus. Incubation and culture conditions were as described in §S1.1.

#### S1.3 Experiment 1: Imaging and image processing

In Experiment 1 to estimate the radius of each spheroid at time $t$, $R(t)$, we use the IncuCyte S3 live cell imaging system (Sartorius, Goettingen, Germany). IncuCyte S3 settings were chosen to image hourly for the duration of the experiment with the 10× objective. Brightfield images captured from above each spheroid with the IncuCyte S3 are processed using the accompanying IncuCyte 2020C Rev1 software (spheroid analysis type, brightfield image channel, largest object area, `hole fill` $= 1 \times 10^4$ µm$^2$). Area masks were visually compared with the IncuCyte brightfield images to confirm accuracy (Figure S1). For each spheroid the mask area, $A(t)$, at time $t$ was converted to an equivalent spheroid radius measurement $R(t) = \sqrt{A(t)/\pi}$. These measurements are used for analysis. At later times cellular debris can form outside of the main spheroid mass, but this is limited and so it is not included in the analysis. Some measurements could not be obtained, primarily due to blurring of the automated imaging.

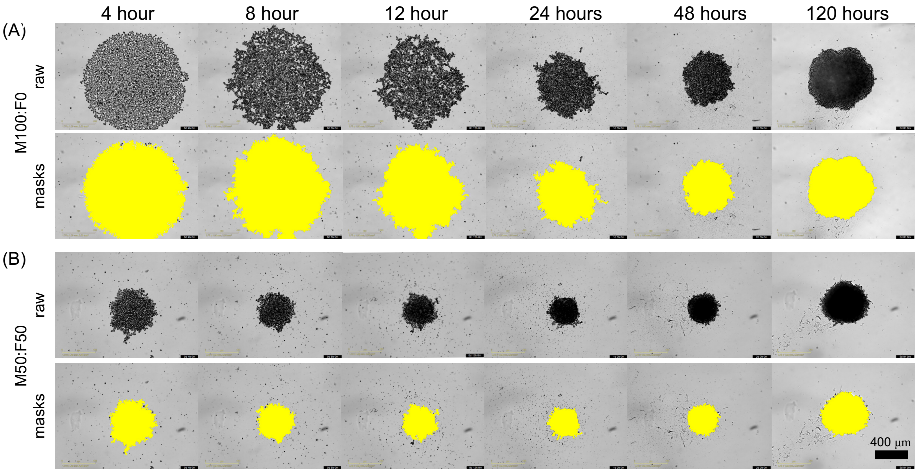

**Figure S1: Brightfield images processing for spheroid formation and growth.** Results shown for (A) M100:F0 and (B) M50:F50 1205Lu co-culture spheroids. In (A,B) top rows show raw images and bottom rows masks (yellow). Top and bottom rows show the same spheroids. Scale bar is 400 µm.

#### S1.4 Experiment 2: Imaging

In Experiment 2, to capture spheroid structure we use confocal microscopy and follow the experimental procedure described in [12] which means that each image is an end-point measurement for that particular spheroid. Spheroids maintained in the incubator were harvested, fixed with 4% paraformaldehyde (PFA), and stored in phosphate buffered saline solution, sodium azide (0.02%), Tween-20 (0.1%), and DAPI (1:2500) at 4 °C, on days 2, 3, 6, 8 and 10 after seeding. Fixed spheroids were set in place using low melting 2% agarose and optically cleared in 500 µL total/well high refractive index mounting solution (Quadrol 9 % wt/wt, Urea 22 % wt/wt, Sucrose 44 % wt/wt, Triton X-100 0.1 % wt/wt, water) for 2 days in a 24-well glass bottom plate (Cellvis, P24-1.5H-N) before imaging to ensure accurate measurements. Images were then captured using an Olympus FV3000 confocal microscope with the 10× objective focused on the equatorial plane of each spheroid.

#### S1.5 Experiment 2: Image processing

We analyse confocal images using standard tools in a CellProfiler pipeline [13, 14] to estimate the spheroid radius, $R(t)$, and necrotic core radius, $R_{\mathrm{n}}(t)$. Methods are similar to [15]. In brief, we use four channels from the raw confocal microscopy images. These channels correspond to: (i) DAPI, a stain of the nucleus of each melanoma and fibroblast cell; (ii) FUCCI green, which identifies melanoma cells in S/G2/M phases of the cell cycle (shown in green); (iii) FUCCI red, which identifies melanoma cells in G1 phase of the cell cycle (shown in magenta), and (iv) a fibroblast marker, which identifies fibroblast cells (shown in cyan).

*Spheroid radius.* To estimate the area covered by each spheroid we use the DAPI signals; binarise, using the `threshold` function; fill gaps between cells, using the `dilate` and `erode` pixels functions with a disk of the same magnitude and then the `RemoveHoles` function; and finally use the `IdentifyPrimaryObjects` function. For each spheroid, the spheroid mask area, $A(t)$, at time $t$ was converted to equivalent radii measurements, $R(t) = \sqrt{A(t)/\pi}$ and used for analysis.

*Necrotic core radius.* To estimate the area of the necrotic core we identify regions that lack FUCCI signals, that this corresponds to the necrotic core has previously been confirmed by both confocal microscopy and flow cytometry using cell death markers [1, 5, 13, 16]. Then the necrotic region is obtained by the following: merge the signals from the FUCCI red and FUCCI green channels using the `GrayToColor` function; convert the merged signals to a grayscale image, using the `ColorToGray` function; binarise the image, using the `threshold` function; remove gaps between cells in the periphery region, using the `dilate`, `RemoveHoles`, and `erode` functions; invert the binary image, using the `ImageMath` function; and then use the `IdentifyPrimaryObjects` function with an upper bound on the maximum size expected to capture the necrotic core rather than the area external to the spheroid. Area masks for the spheroid and necrotic core were visually compared to confirm accuracy (§S2.2). For each spheroid, the necrotic core mask area, $A_{\mathrm{n}}(t)$, at time $t$ was converted to equivalent radii measurements, $R_{\mathrm{n}}(t) = \sqrt{A_{\mathrm{n}}(t)/\pi}$ and used for analysis.

### S2 Experimental data

Here we summarise the experimental data and present additional experimental images.

#### S2.1 Summary

In Table S1 we present the number of spheroids imaged and measured in Experiment 2. These images are used to analyse the temporal evolution of spheroid structure, namely spheroid radius, $R(t)$, and necrotic radius, $R_{\text{n}}(t)$. Each measurement is an end-point measurement since we harvest, fix, and mount each spheroid before imaging. Day 0 corresponds to the start of the experiment when spheroids were seeded.

| Condition | Day | WM983B |
|---|---|---|
| M100:F0 | 2 | 8 |
| | 3 | 7 |
| | 6 | 8 |
| | 8 | 8 |
| | 10 | 10 |
| M75:F25 | 2 | 8 |
| | 3 | 8 |
| | 6 | 8 |
| | 8 | 8 |
| | 10 | 8 |
| M50:F50 | 2 | 7 |
| | 3 | 6 |
| | 6 | 8 |
| | 8 | 8 |
| | 10 | 8 |
| M25:F75 | 2 | 8 |
| | 3 | 6 |
| | 6 | 8 |
| | 8 | 8 |
| | 10 | 8 |

**Table S1: Number of spheroids imaged and measured in Experiment 2.** Results shown for the co-culture spheroid experiments performed with the WM983B cell line and fibroblasts.

#### S2.2 Experimental images

Here we present experimental images from Experiment 2. Images capture co-culture experiments performed with the WM983B melanoma cell line and fibroblasts (Figs S2-S5). FUCCI signals indicate different stages of the cell cycle: cells in gap 1 (G1) phase fluoresce red, shown in magenta for clarity; and cells in synthesis, gap 2, and mitotic (S/G2/M) phases fluoresce green. At late times the large central regions without FUCCI signals indicate a necrotic core. Outside of the necrotic core FUCCI signals indicate a proliferating region at the periphery (predominately green) and an intermediate region of living cells that are proliferation-inhibited (predominately magenta). Fibroblasts are identified using CellTracker™ Deep Red dye shown in cyan. The fibroblast marker stains the entire cell whereas the FUCCI signal is only present at the cell nucleus.

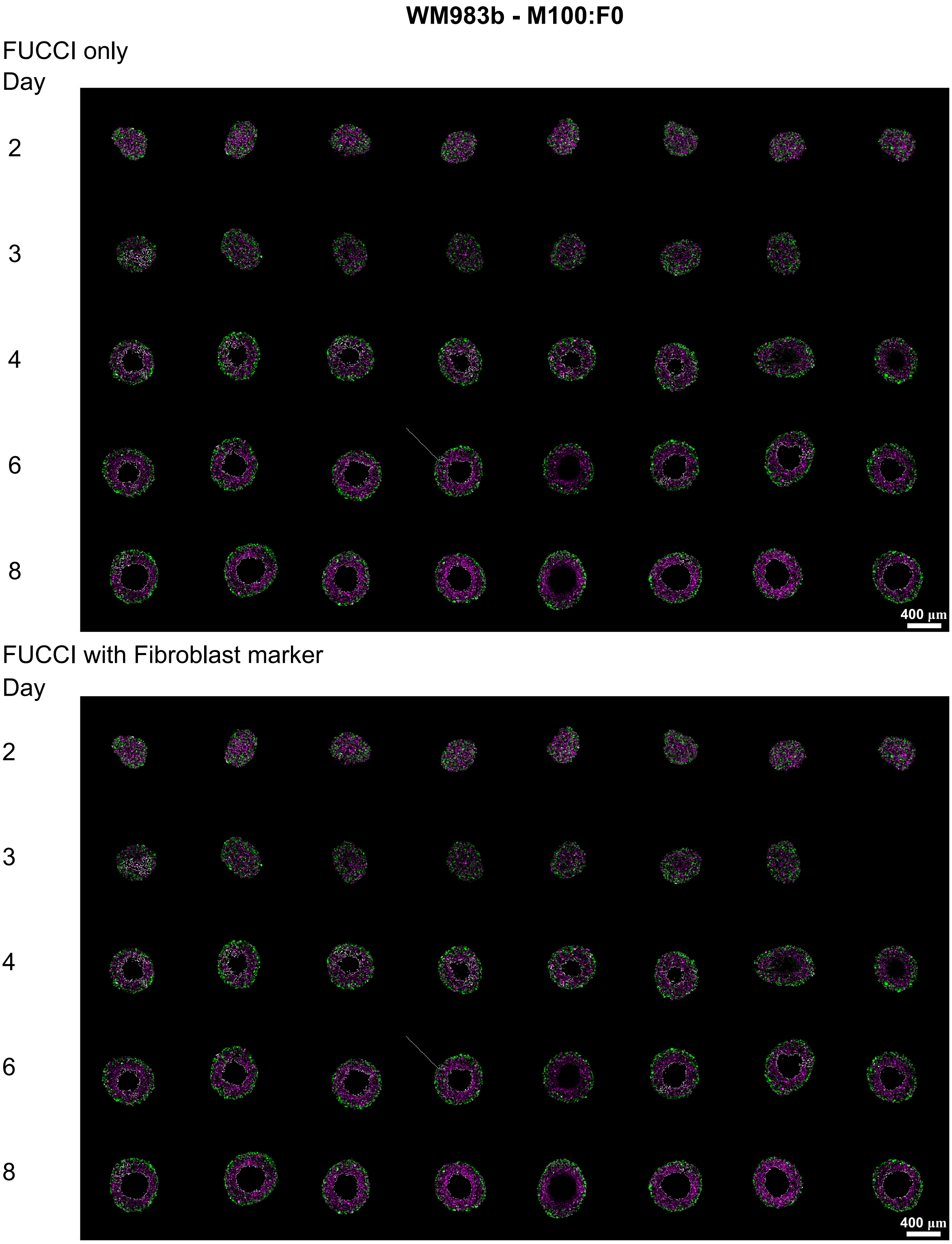

**Figure S2: Confocal experimental images of M100:F0 spheroids.** Top images show FUCCI signals (magenta, green) that we use to identify melanoma cells. Bottom images show signals from FUCCI and the fibroblast marker (cyan). There are no fibroblasts in this condition therefore there are no signals from the fibroblast marker. Scale bar is 400µm.

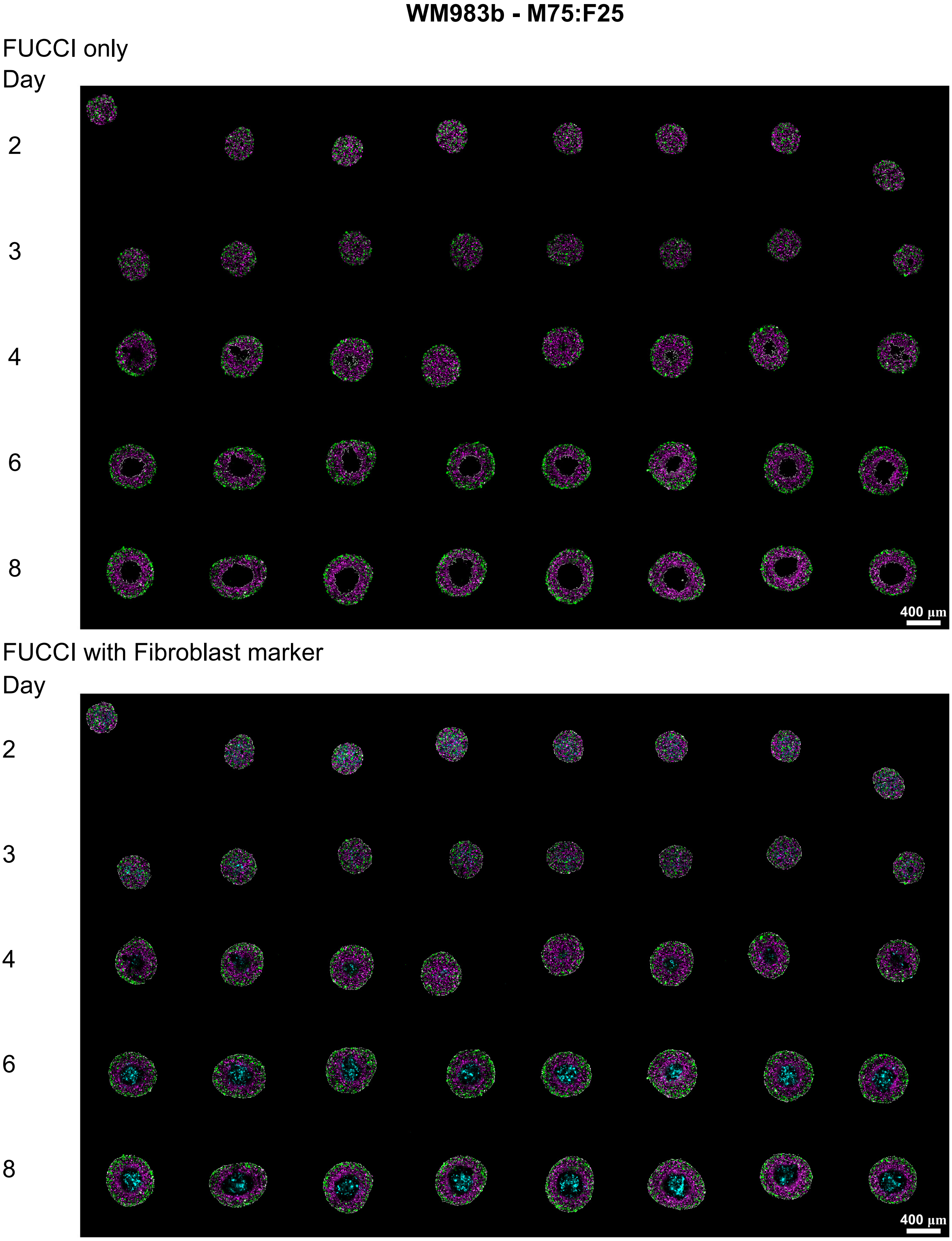

**Figure S3: Confocal experimental images of M75:F25 spheroids.** Top images show FUCCI signals (magenta, green) that we use to identify melanoma cells. Bottom images show signals from FUCCI and the fibroblast marker (cyan). Scale bars are 400µm.

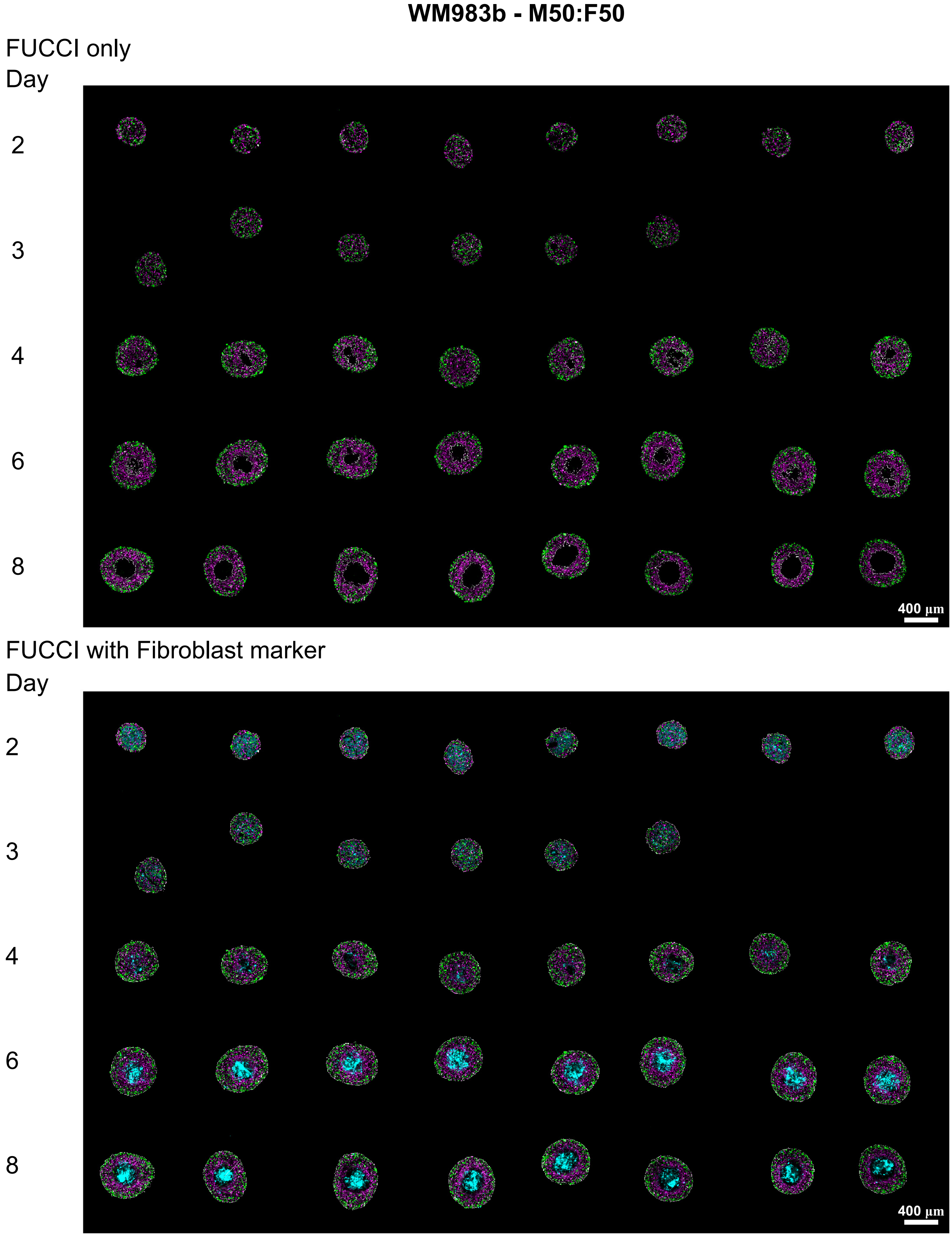

**Figure S4: Confocal experimental images of M50:F50 spheroids.** Top images show FUCCI signals (magenta, green) that we use to identify melanoma cells. Bottom images show signals from FUCCI and the fibroblast marker (cyan). Scale bars are 400µm.

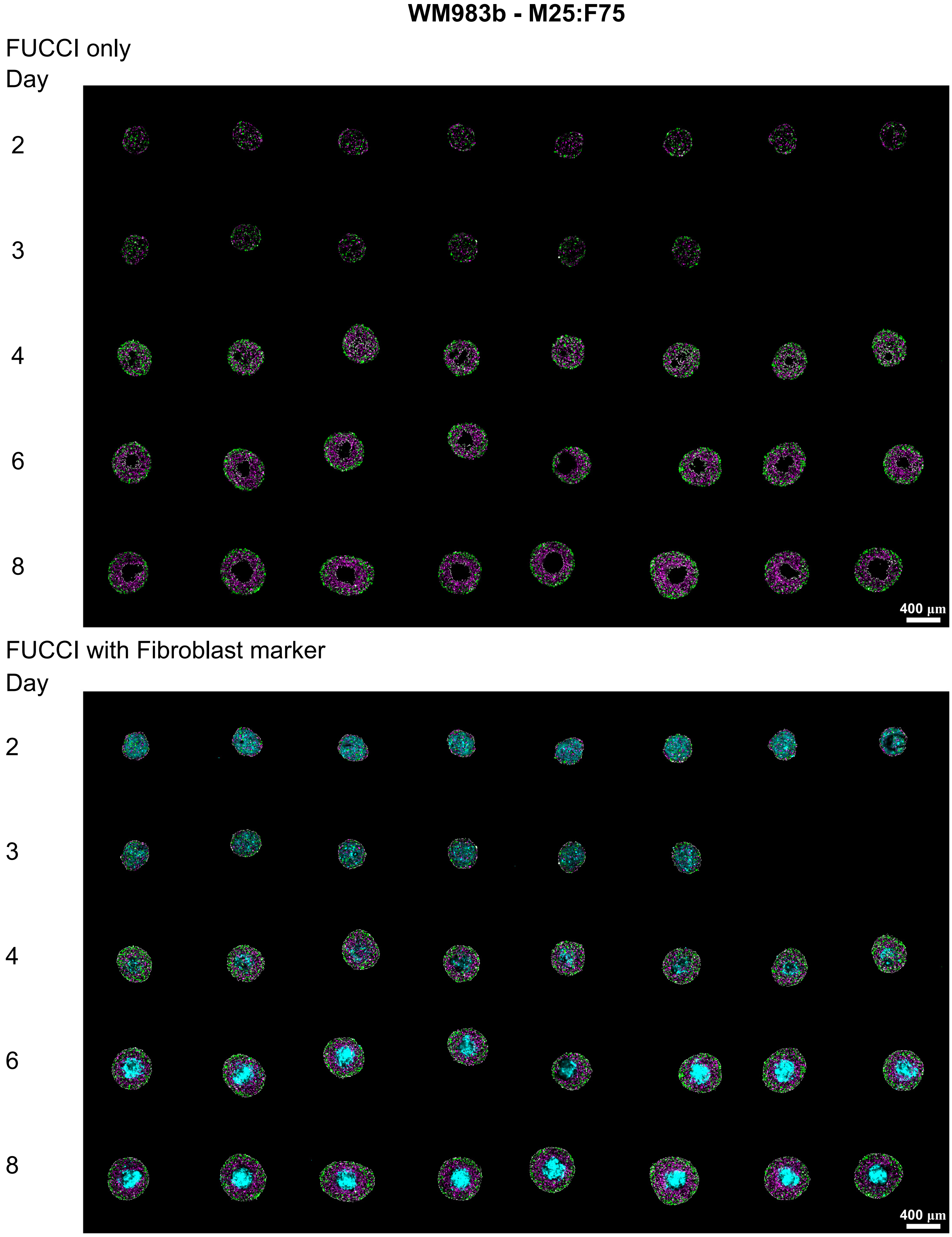

**Figure S5: Confocal experimental images of M25:F75 spheroids.** Top images show FUCCI signals (magenta, green) that we use to identify melanoma cells. Bottom images show signals from FUCCI and the fibroblast marker (cyan). Scale bars are 400µm.

### S3 Additional mathematical methods

#### S3.1 Linear approximation to the biphasic model

Here we present the linear approximation to early time growth dynamics in the biphasic model. Focusing on $t \geq T$ Eq. (1) is

$$\frac{\mathrm{d}R(t)}{\mathrm{d}t} = r_2 R(t) \left(1 - \frac{R(t)}{\mathcal{R}_2}\right). \tag{S.1}$$

Assuming that the size of the spheroid during the experiment is much smaller than the long-time maximum spheroid size, $\mathcal{R}_2$, i.e. $R(t) \ll \mathcal{R}_2$ for $T \leq t \leq 10$ [days]. Therefore, $1 - R(t)/\mathcal{R}_2 \approx 1$ and Eq. (S.1) can be written as

$$\frac{\mathrm{d}R(\tilde{t})}{\mathrm{d}\tilde{t}} = r_2 R(\tilde{t}), \tag{S.2}$$

where $\tilde{t} = t - T \geq 0$. We solve Eq (S.2) analytically to obtain

$$R(\tilde{t}) = R(T)\exp(r_2\tilde{t}). \tag{S.3}$$

Assuming $r_2\tilde{t}$ is small gives

$$R(T)\exp(\lambda\tilde{t}) = R(0)[1 + \lambda\tilde{t} + \mathcal{O}((\lambda\tilde{t})^2)]. \tag{S.4}$$

where we have relabelled $r_2$ with $\lambda$. Finally, neglecting higher order terms and rewriting in terms of $t$ instead of $\tilde{t}$ we obtain the linear model presented in Eq. (2).

#### S3.2 Solving the general compartment model

In the main manuscript we state the general compartment based model for tumour spheroid growth (Eqs 3.1-3.6) and transform this to a reduced system (Eq 4.1-4.4) that we solve numerically. In this section we present how to obtain Eq (4.2) and Eq (4.3). Note that Eq (4.1) is identical to Eq (3.1) and Eq (4.4) is identical to Eq (3.6). We start by rewriting Eqs 3.1-3.6,

$$\frac{\mathrm{d}V_i^{(j)}(t)}{\mathrm{d}t} = \begin{cases} \underbrace{f_{V_i^{(j)}}\left(R(t);p\right)}_{\text{source, sink, reaction, migration}} - \underbrace{g^{(j)}(R(t);p)}_{\substack{\text{migration due to}\\ \text{boundary constraint}}} \underbrace{g_i^{(j)}(R(t);p),}_{\text{proportion}\in[0,1]} \quad j=1,\ i=1,2,\dots,I, \\ f_{V_i^{(j)}}\left(R(t);p\right) + g^{(j-1)}(R(t);p)g_i^{(j-1)}(R(t);p) - g^{(j)}(R(t);p)g_i^{(j)}(R(t);p), \\ \qquad\qquad j=1,\dots,J-1,\ i=1,2,\dots,I, \\ f_{V_i^{(j)}}\left(R(t);p\right) + g^{(j-1)}(R(t);p)g_i^{(j-1)}(R(t);p), \quad j=J,\ i=1,2,\dots,I, \end{cases} \tag{S.5.1}$$

$$\sum_{i=1}^{I} g_i^{(j)}(t) = 1, \quad j=1,2,\dots,J, \text{ with } g_i^{(j)}(t)\geq 0, \text{ for } \quad i=1,2,\dots,I,\ j=1,2,\dots,J-1, \tag{S.5.2}$$

$$V^{(j)}(t) = \sum_{i=1}^{I} V_i^{(j)}(t), \quad j=1,2,\dots,J, \tag{S.5.3}$$

$$V(t) = \sum_{j=1}^{J} V^{(j)}(t), \tag{S.5.4}$$

$$V^{(j)}(t) = \begin{cases} \frac{4\pi}{3}\left(R_{j-1}(t)^3 - R_j(t)^3\right), \quad j=1,2,\dots,J-1, \\ \frac{4\pi}{3}R_j(t)^3, \quad j=J \end{cases} \tag{S.5.5}$$

$$R_j(t) = \underbrace{f_j\left(R(t);p\right),}_{\text{boundary constraint}} \quad j=1,2,\dots.J-1. \tag{S.5.6}$$

To obtain Eq (4.2), we use Eqs (S.5.5), (S.5.3), and (S.5.1)

$$4\pi R_0(t)^2\frac{\mathrm{d}R_0(t)}{\mathrm{d}t} = \frac{\mathrm{d}}{\mathrm{d}t}\left(\frac{4\pi}{3}R_0(t)^3\right) \tag{S.6.1}$$

$$= \frac{\mathrm{d}}{\mathrm{d}t}\left(\sum_{j=1}^{J} V^{(j)}(t)\right) \tag{S.6.2}$$

$$= \frac{\mathrm{d}}{\mathrm{d}t}\left(\sum_{j=1}^{J}\sum_{i=1}^{I} V_i^{(j)}(t)\right) \tag{S.6.3}$$

$$= \sum_{j=1}^{J}\sum_{i=1}^{I}\frac{\mathrm{d}V_i^{(j)}(t)}{\mathrm{d}t} \tag{S.6.4}$$

$$= \sum_{j=1}^{J}\sum_{i=1}^{I} f_{V_i^{(j)}}\left(R(t);p\right). \tag{S.6.5}$$

To obtain Eq (4.3), we start by considering $j = 1$ in Eq (S.5.1) and sum over $i$ to obtain

$$\sum_{i=1}^{I} \frac{\mathrm{d}V_i^{(1)}(t)}{\mathrm{d}t} = \sum_{i=1}^{I} f_{V_i^{(1)}}(R(t);p) - \sum_{i=1}^{I} g^{(1)}(R(t);p) g_i^{(1)}(R(t);p)) \tag{S.7.1}$$

$$= \sum_{i=1}^{I} f_{V_i^{(1)}}(R(t);p) - g^{(1)}(R(t);p) \tag{S.7.2}$$

where we have used Eq (S.5.2) to simplify the summation. Rearranging Eq (S.7.2), using Eqs (S.5.3), (S.5.5), and (S.5.6), and recalling that $R(t) = R_0(t)$ gives,

$$g^{(1)}(R(t);p) = \sum_{i=1}^{I} f_{V_i^{(1)}}(R(t);p) - \sum_{i=1}^{I} \frac{\mathrm{d}V_i^{(1)}(t)}{\mathrm{d}t} \tag{S.8.1}$$

$$= \sum_{i=1}^{I} f_{V_i^{(1)}}(R(t);p) - \frac{\mathrm{d}V^{(1)}(t)}{\mathrm{d}t} \tag{S.8.2}$$

$$= \sum_{i=1}^{I} f_{V_i^{(1)}}(R(t);p) - \frac{\mathrm{d}}{\mathrm{d}t}\left(\frac{4\pi}{3}\left(R_0(t)^3 - R_1(t)^3\right)\right) \tag{S.8.3}$$

$$= \sum_{i=1}^{I} f_{V_i^{(1)}}(R(t);p) - 4\pi\left(R_0(t)^2\frac{\mathrm{d}R_0(t)}{\mathrm{d}t} - R_1(t)^2\frac{\mathrm{d}R_1(t)}{\mathrm{d}t}\right) \tag{S.8.4}$$

$$= \sum_{i=1}^{I} f_{V_i^{(1)}}(R(t);p) - 4\pi\left(R_0(t)^2\frac{\mathrm{d}R_0(t)}{\mathrm{d}t} - R_1(t)^2\frac{\mathrm{d}f_j(R_0(t);p)}{\mathrm{d}t}\right) \tag{S.8.5}$$

$$= \sum_{i=1}^{I} f_{V_i^{(1)}}(R(t);p) - 4\pi\left(R_0(t)^2\frac{\mathrm{d}R_0(t)}{\mathrm{d}t} - R_1(t)^2\frac{\mathrm{d}f_j(R_0(t);p)}{\mathrm{d}R_0(t)}\frac{\mathrm{d}R_0(t)}{\mathrm{d}t}\right) \tag{S.8.6}$$

$$= \sum_{i=1}^{I} f_{V_i^{(1)}}(R(t);p) - 4\pi\left(R_0(t)^2\frac{\mathrm{d}R_0(t)}{\mathrm{d}t} - R_1(t)^2\frac{\mathrm{d}R_1(t)}{\mathrm{d}R(t)}\frac{\mathrm{d}R_0(t);p)}{\mathrm{d}t}\right) \tag{S.8.7}$$

$$= \sum_{i=1}^{I} f_{V_i^{(1)}}(R(t);p) - 4\pi\frac{\mathrm{d}R_0(t)}{\mathrm{d}t}\left(R_0(t)^2 - R_1(t)^2\frac{\mathrm{d}R_1(t)}{\mathrm{d}R_0(t)}\right) \tag{S.8.8}$$

For $j = 2$ we proceed similarly and use Eq (S.8.8) to obtain,

$$g^{(2)}(R(t);p) = \sum_{i=1}^{I} f_{V_i^{(2)}}(R(t);p) + \sum_{i=1}^{I} g^{(1)}(R(t);p) g_i^{(1)}(R(t);p) - \sum_{i=1}^{I} \frac{\mathrm{d}V_i^{(2)}(t)}{\mathrm{d}t} \tag{S.9.1}$$

$$= \sum_{i=1}^{I} f_{V_i^{(2)}}(R(t);p) + g^{(1)}(R(t);p) - \frac{\mathrm{d}V^{(2)}(t)}{\mathrm{d}t} \tag{S.9.2}$$

$$= \sum_{i=1}^{I} f_{V_i^{(2)}}(R(t);p) + g^{(1)}(R(t);p) - \frac{\mathrm{d}}{\mathrm{d}t}\left(\frac{4\pi}{3}\left(R_1(t)^3 - R_2(t)^3\right)\right) \tag{S.9.3}$$

$$= \sum_{i=1}^{I} f_{V_i^{(2)}}(R(t);p) + g^{(1)}(R(t);p) - 4\pi\left(R_1(t)^2\frac{\mathrm{d}R_1(t)}{\mathrm{d}t} - R_2(t)^2\frac{\mathrm{d}R_2(t)}{\mathrm{d}t}\right) \tag{S.9.4}$$

$$= \sum_{i=1}^{I} f_{V_i^{(2)}}(R(t);p) + g^{(1)}(R(t);p) - 4\pi\left(R_1(t)^2\frac{\mathrm{d}R_1(t)}{\mathrm{d}R_0(t)}\frac{\mathrm{d}R_0(t)}{\mathrm{d}t} - R_2(t)^2\frac{\mathrm{d}R_2(t)}{\mathrm{d}R_0(t)}\frac{\mathrm{d}R_0(t)}{\mathrm{d}t}\right) \tag{S.9.5}$$

$$= \sum_{k=1}^{2}\sum_{i=1}^{I} f_{V_i^{(k)}}(R(t);p) - 4\pi\frac{\mathrm{d}R_0(t)}{\mathrm{d}t}\left(R_0(t)^2 - R_2(t)^2\frac{\mathrm{d}R_2(t)}{\mathrm{d}R_0(t)}\right). \tag{S.9.6}$$

It is then straightforward to obtain the general formula presented in Eq (4.3).

#### S3.3 Monoculture reduced Greenspan Model: Deriving the constraint from oxygen mechanisms

In main manuscript §4.2 we present the monoculture reduced Greenspan model. Here we detail how we define $R_\mathrm{n}(t)$ using a boundary constraint obtained by considering oxygen diffusion and consumption.

To define $R_\mathrm{n}(t)$ we consider the oxygen partial pressure within the spheroid. Spatial and temporal differences within spheroids in this study, with respect to cell survival, are reasonably explained by availability of oxygen [2, 17]. Following [17] and our previous study [2], we report oxygen at a radial distance $r$ from the centre of the spheroid and time $t$, in terms of the oxygen partial pressure $p(r,t)$ [%] for $0 \leq r \leq R(t)$. Oxygen diffuses within the spheroid with diffusivity $k$ [m$^2$ s$^{-1}$] and the rate of volume of oxygen gas per unit tumour mass that is consumed by living cells is a constant $\alpha$ [m$^3$ kg$^{-1}$ s$^{-1}$]. The external oxygen partial pressure is $p_\infty$ [%]. Since oxygen takes approximately 10 seconds to diffuse across a distance of 100 µm [18], and this timescale is much faster than the timescales of cell proliferation and spheroid growth, we assume that oxygen within the spheroid is at diffusive equilibrium. Therefore, at any instant in time we have $p(r,t) = p(r)$ due to fast diffusion of oxygen. However, as the spheroid is growing with time, diffusion of oxygen occurs on a growing domain and we write $p(r) = p(r(t))$. The governing equation for the oxygen partial pressure within the spheroid, $0 \leq r \leq R(t)$, is,

$$\underbrace{\frac{1}{r^2}\frac{\partial}{\partial r}\left(r^2\frac{\partial}{\partial r}p(r(t))\right)}_{\text{diffusion}} = \underbrace{\frac{\Omega\alpha}{k}\mathrm{H}\left(r - R_\mathrm{n}(t)\right)\mathrm{H}\left(R(t) - r\right),}_{\text{consumption in } R_\mathrm{n}(t)<r<R(t)} \tag{S.10}$$

where $\mathrm{H}(\cdot)$ denotes the Heaviside function and $\Omega = 3.0318 \times 10^7$ [mmHg kg m$^{-3}$] is a conversion constant from volume of oxygen gas per unit tumour mass to partial pressure. Since we assume that oxygen is only consumed by living cells there is no oxygen consumption in $0 \leq r \leq R_\mathrm{n}(t)$. Solving Eq (S.10) for the oxygen partial pressure in each region gives

$$p(r,t) = \begin{cases} p_\infty - \dfrac{\Omega\alpha}{6k}\left[R(t)^2 - r^2\right] + \dfrac{\Omega\alpha}{3k}R_\mathrm{n}(t)^3\left[\dfrac{1}{r} - \dfrac{1}{R(t)}\right], & R_\mathrm{n}(t) \leq r \leq R(t), \\ p_\infty - \dfrac{\Omega\alpha}{6k}\left[R(t)^2 - R_\mathrm{n}(t)^2 - \dfrac{2R_\mathrm{n}(t)^2}{R(t)}\left(R(t) - R_\mathrm{n}(t)\right)\right], & 0 \leq r \leq R_\mathrm{n}(t). \end{cases} \tag{S.11}$$

where we have assumed $p(r,t)$ and $\partial p(r,t)/\partial r$ are continuous at $r = R_\mathrm{n}(t)$, and $p(r,t)$ is bounded at $r = 0$. These assumptions determine constants of integration that arise when solving Eq (S.10).

We then define $R_\mathrm{n}(t)$ implicitly via

$$p(R_\mathrm{n}(t), t) = p_\mathrm{n}, \tag{S.12}$$

provided the oxygen partial pressure is sufficiently small, otherwise $R_\mathrm{n}(t) = 0$. Here, $p_\mathrm{n}$ [%] corresponds to the oxygen partial pressure threshold below which cells die (Fig 1E,F).

The oxygen partial pressure first reaches $p_\mathrm{n}$ at the centre of the spheroid at the end of phase (i) and the start of phase (ii), that we refer to as time $T_1$. By definition $R_\mathrm{n}(T_1) = 0$. Setting $r = 0$ in the solution of the oxygen partial pressure in Eq (S.11), corresponding to the region $R_\mathrm{n}(t) < r < R(t)$, and using the oxygen threshold in Eq (S.12) we obtain that the radius of the spheroid when the necrotic region first forms, $\mathcal{R}$ [µm], is

$$\mathcal{R}^2 = \frac{6k}{\Omega\alpha}\left(p_\infty - p_\mathrm{n}\right). \tag{S.13}$$

To obtain an expression for $\mathcal{R}$ in terms of radial measurements $R(t)$ and $R_{\mathrm{n}}(t)$, and valid for $R(t) > \mathcal{R}$, we evaluate the oxygen partial pressure at $r = R_{\mathrm{n}}(t)$ in Eq (S.11) to obtain

$$\mathcal{R}^2 = R(t)^2 - R_{\mathrm{n}}(t)^2 - \frac{2R_{\mathrm{n}}(t)^2}{R(t)}\left[R(t) - R_{\mathrm{n}}(t)\right]. \tag{S.14}$$

#### S3.4 Co-culture Model 1: Governing equations

In main manuscript §4.3 we present Eqs (8.1)-(8.5) that define Co-culture Model 1 describing heterogeneous proliferation rates. Here we detail the governing equations. Substituting Eqs (8.1)-(8.5) into Eqs (4.1-4.4) gives equations for the temporal evolution the volume occupied by each population in each compartment (Eqs (S.15.1)-(S.15.4)), the temporal evolution of overall spheroid size (Eq (S.15.5)), and an equation for the rate of transfer of living cells to the necrotic core (Eq (S.15.6),

$$\frac{\mathrm{d}V_1^{(1)}(t)}{\mathrm{d}t} = s_{\mathrm{m}}\frac{4\pi}{3}\left(R(t)^3 - R_{\mathrm{n}}(t)^3\right)\frac{V_1^{(1)}(t)}{V_1^{(1)}(t)+V_2^{(1)}(t)} - g^{(1)}(R(t);p)\frac{V_1^{(1)}(t)}{V_1^{(1)}(t)+V_2^{(1)}(t)}, \tag{S.15.1}$$

$$\frac{\mathrm{d}V_2^{(1)}(t)}{\mathrm{d}t} = s_{\mathrm{f}}\frac{4\pi}{3}\left(R(t)^3 - R_{\mathrm{n}}(t)^3\right)\frac{V_2^{(1)}(t)}{V_1^{(1)}(t)+V_2^{(1)}(t)} - g^{(1)}(R(t);p)\frac{V_2^{(1)}(t)}{V_1^{(1)}(t)+V_2^{(1)}(t)}, \tag{S.15.2}$$

$$\frac{\mathrm{d}V_1^{(2)}(t)}{\mathrm{d}t} = -3s_{\mathrm{m}}\gamma\frac{4\pi}{3}R_{\mathrm{n}}(t)^3\frac{V_2^{(2)}(t)}{V_1^{(2)}(t)+V_2^{(2)}(t)} + g^{(1)}(R(t);p)\frac{V_1^{(1)}(t)}{V_1^{(1)}(t)+V_2^{(1)}(t)}, \tag{S.15.3}$$

$$\frac{\mathrm{d}V_2^{(2)}(t)}{\mathrm{d}t} = -3s_{\mathrm{m}}\gamma\frac{4\pi}{3}R_{\mathrm{n}}(t)^3\frac{V_2^{(2)}(t)}{V_1^{(2)}(t)+V_2^{(2)}(t)} + g^{(1)}(R(t);p)\frac{V_2^{(1)}(t)}{V_1^{(1)}(t)+V_2^{(1)}(t)}, \tag{S.15.4}$$

$$\frac{\mathrm{d}R(t)}{\mathrm{d}t} = \frac{1}{R(t)^2}\left[\left(\frac{s_{\mathrm{m}}}{3}\frac{V_1^{(1)}(t)}{V_1^{(1)}(t)+V_2^{(1)}(t)} + \frac{s_2}{3}\frac{V_2^{(1)}(t)}{V_1^{(1)}(t)+V_2^{(1)}(t)}\right)\left(R(t)^3 - R_{\mathrm{n}}(t)^3\right) - s_1\gamma R_{\mathrm{n}}(t)^3\right], \tag{S.15.5}$$

$$\begin{aligned} g^{(1)}(R(t);p) = {} & \frac{4\pi}{3}\left[\left(\frac{s_{\mathrm{m}}}{3}\frac{V_1^{(1)}(t)}{V_1^{(1)}(t)+V_2^{(1)}(t)} + \frac{s_2}{3}\frac{V_2^{(1)}(t)}{V_1^{(1)}(t)+V_2^{(1)}(t)}\right)\left(R(t)^3 - R_{\mathrm{n}}(t)^3\right)\right] \\ & - 4\pi\frac{\mathrm{d}R(t)}{\mathrm{d}t}\left(R(t)^2 - R_{\mathrm{n}}(t)^2\frac{\mathrm{d}R_{\mathrm{n}}(t)}{\mathrm{d}R(t)}\right). \end{aligned} \tag{S.15.6}$$

For Eq (S.15.6) we obtain $\mathrm{d}R_{\mathrm{n}}(t)/\mathrm{d}R(t)$ by differentiating Eq (8.5) with respect to $R(t)$ and rearranging

$$\frac{\mathrm{d}R_{\mathrm{n}}(t)}{\mathrm{d}R(t)} = \begin{cases} 0, & \text{for } R(t) \leq \mathcal{R}, \\ \dfrac{2R(t) + \dfrac{2R_{\mathrm{n}}(t)^2}{R(t)^2}\left(R(t) - R_{\mathrm{n}}(t)\right) - \dfrac{2R_{\mathrm{n}}(t)^2}{R(t)}}{2R_{\mathrm{n}}(t) + \dfrac{4R_{\mathrm{n}}(t)}{R(t)}\left(R(t) - R_{\mathrm{n}}(t)\right) - \dfrac{2R_{\mathrm{n}}(t)^2}{R(t)}}, & \text{for } R(t) > \mathcal{R}. \end{cases} \tag{S.16}$$

#### S3.5 Co-culture Model 2: Governing equations

In main manuscript §4.4 we present Eqs (9.1-9.5) that define Co-culture Model 2 describing heterogeneous loss rates. Here we detail the governing equations. Substituting Eqs (9.1-9.5) into Eqs (4.1-4.4) gives equations for the temporal evolution of the volume occupied by each population in each compartment (Eqs (S.17.1)-(S.17.4)), the temporal evolution of overall spheroid size (Eq (S.17.5)), and an equation for the rate of transfer of living cells to the necrotic core (Eq (S.15.6)),

$$\frac{\mathrm{d}V_1^{(1)}(t)}{\mathrm{d}t} = s_\mathrm{m}\frac{4\pi}{3}\left(R(t)^3 - R_\mathrm{n}(t)^3\right)\frac{V_1^{(1)}(t)}{V_1^{(1)}(t)+V_2^{(1)}(t)} - g^{(1)}(R(t);p)\frac{V_1^{(1)}(t)}{V_1^{(1)}(t)+V_2^{(1)}(t)}, \tag{S.17.1}$$

$$\frac{\mathrm{d}V_2^{(1)}(t)}{\mathrm{d}t} = s_\mathrm{f}\frac{4\pi}{3}\left(R(t)^3 - R_\mathrm{n}(t)^3\right)\frac{V_2^{(1)}(t)}{V_1^{(1)}(t)+V_2^{(1)}(t)} - g^{(1)}(R(t);p)\frac{V_2^{(1)}(t)}{V_1^{(1)}(t)+V_2^{(1)}(t)}, \tag{S.17.2}$$

$$\frac{\mathrm{d}V_1^{(2)}(t)}{\mathrm{d}t} = -3s_\mathrm{m}\gamma\frac{4\pi}{3}R_\mathrm{n}(t)^3\frac{V_2^{(2)}(t)}{V_1^{(2)}(t)+V_2^{(2)}(t)} + g^{(1)}(R(t);p)\frac{V_1^{(1)}(t)}{V_1^{(1)}(t)+V_2^{(1)}(t)}, \tag{S.17.3}$$

$$\frac{\mathrm{d}V_2^{(2)}(t)}{\mathrm{d}t} = -3s_\mathrm{m}\gamma\frac{4\pi}{3}R_\mathrm{n}(t)^3\frac{V_2^{(2)}(t)}{V_1^{(2)}(t)+V_2^{(2)}(t)} + g^{(1)}(R(t);p)\frac{V_2^{(1)}(t)}{V_1^{(1)}(t)+V_2^{(1)}(t)}, \tag{S.17.4}$$

$$\frac{\mathrm{d}R(t)}{\mathrm{d}t} = \frac{1}{R(t)^2}\left[\left(\frac{s_\mathrm{m}}{3}\frac{V_1^{(1)}(t)}{V_1^{(1)}(t)+V_2^{(1)}(t)} + \frac{s_2}{3}\frac{V_2^{(1)}(t)}{V_1^{(1)}(t)+V_2^{(1)}(t)}\right)\left(R(t)^3 - R_\mathrm{n}(t)^3\right) - s_1\gamma R_\mathrm{n}(t)^3\right], \tag{S.17.5}$$

$$g^{(1)}(R(t);p) = \frac{4\pi}{3}\left[\left(\frac{s_\mathrm{m}}{3}\frac{V_1^{(1)}(t)}{V_1^{(1)}(t)+V_2^{(1)}(t)} + \frac{s_2}{3}\frac{V_2^{(1)}(t)}{V_1^{(1)}(t)+V_2^{(1)}(t)}\right)\left(R(t)^3 - R_\mathrm{n}(t)^3\right)\right] - 4\pi\frac{\mathrm{d}R(t)}{\mathrm{d}t}\left(R(t)^2 - R_\mathrm{n}(t)^2\frac{\mathrm{d}R_\mathrm{n}(t)}{\mathrm{d}R(t)}\right). \tag{S.17.6}$$

For Eq (S.17.6) $\mathrm{d}R_\mathrm{n}(t)/\mathrm{d}R(t)$ is given by Eq (S.16).

#### S3.6 Co-culture Model 3: Deriving the constraint from oxygen mechanisms

In main manuscript §4.2 we present the Co-culture Model 3. Here we detail how we define $R_\mathrm{m}(t)$ and $R_\mathrm{n}(t)$ by considering oxygen diffusion and consumption.

To define $R_\mathrm{m}(t)$ and $R_\mathrm{f}(t)$ we consider the oxygen partial pressure within the spheroid. In agreement with the two compartment model, spatial and temporal differences within spheroids in this study, with respect to cell survival, are thought to arise due to availability of oxygen. Considering the proportion of each population in each region of the spheroid and recalling that only living cells consume oxygen, the governing equation for the oxygen partial pressure within the spheroid is

$$\underbrace{\frac{1}{r^2}\frac{\partial}{\partial r}\left(r^2\frac{\partial}{\partial r}p(r(t))\right)}_{\text{diffusion}} = \begin{cases} \dfrac{\Omega\alpha}{k}, & R_\mathrm{f}(t) \le r \le R(t), \\ \dfrac{\Omega\alpha}{k}\dfrac{V_1^{(2)}(t)}{V_1^{(2)}(t)+V_2^{(2)}(t)}, & R_\mathrm{m}(t) \le r \le R_\mathrm{f}(t), \\ 0, & 0 \le r \le R_\mathrm{m}(t). \end{cases} \tag{S.18}$$

Oxygen within the spheroid is assumed to be at diffusive equilibrium and we assume that oxygen diffusion is fast relative to the migration of cells across the compartment boundaries. Under these assumptions we treat $\phi = V_1^{(2)}(t)/\left(V_1^{(2)}(t)+V_2^{(2)}(t)\right)$ as constant at each time point and solve Eq (S.18) analytically

$$p(r,t) = \begin{cases} p_\infty + \dfrac{\Omega\alpha}{6k}\dfrac{1}{rR(t)}\Big[-(rR(t)^2 + R(t)r^2 + \left(2R_\mathrm{f}(t)^3 - 2R_\mathrm{m}(t)^3\right)\phi \\ \quad -2R_\mathrm{f}(t)^3)(-r+R(t))\Big], & R_\mathrm{f}(t) \le r \le R(t), \\ p_\infty + \dfrac{\Omega\alpha}{6k}\dfrac{1}{rR(t)}\Big[\phi r^3 R(t) + \Big(-R(t)^3 - 3R_\mathrm{f}(t)^2(\phi-1)R(t) \\ \quad + \left(2R_\mathrm{f}(t)^3 - 2R_\mathrm{m}(t)^3\right)\phi - 2R_\mathrm{f}(t)^3\Big)r + 2R_\mathrm{m}(t)^3\phi R(t)\Big], & R_\mathrm{m}(t) \le r \le R_\mathrm{f}(t), \\ p_\infty + \dfrac{\Omega\alpha}{6k}\dfrac{1}{R(t)}\Big[-R(t)^3 + \left(\left(-3R_\mathrm{f}(t)^2 + 3R_\mathrm{m}(t)^2\right)\phi + 3R_\mathrm{f}(t)^2\right)R(t) \\ \quad + \left(2R_\mathrm{f}(t)^3 - 2R_\mathrm{m}(t)^3\right)\phi - 2R_\mathrm{f}(t)^3\Big], & 0 \le r \le R_\mathrm{m}(t). \end{cases} \tag{S.19}$$

where we have assumed $p(r,t)$ and $\partial p(r,t)/\partial r$ are continuous at $r = R_\mathrm{m}(t)$ and $r = R_\mathrm{f}(t)$, and $p(r,t)$ is bounded at $r = 0$ to determine the constants of integration that arise when solving Eq (S.19).

The two oxygen partial pressure thresholds, $p_\mathrm{f}$ and $p_\mathrm{m}$, define $R_\mathrm{f}(t)$ and $R_\mathrm{m}(t)$ via (Fig 1K,L)

$$p(R_\mathrm{f}(t),t) = p_\mathrm{f}, \tag{S.20.1}$$

$$p(R_\mathrm{m}(t),t) = p_\mathrm{m}. \tag{S.20.2}$$

The oxygen partial pressure first reaches the oxygen partial pressure threshold $p_\mathrm{f}$ at $t = T_1$, corresponding to the end of phase (i) and the start of phase (ii). By definition $R_\mathrm{f}(T_1) = 0$ and $R_\mathrm{m}(T_1) = 0$. Setting $r = 0$ in the solution of the oxygen partial pressure for the region $R_\mathrm{f}(t) < r < R(t)$ of Eq (S.19) and using the oxygen threshold in Eq (S.20.1) we obtain that the radius of the spheroid when fibroblasts first undergo necrosis, $\mathcal{R}_\mathrm{f}$ [µm], is

$$\mathcal{R}_\mathrm{f}^2 = \frac{6k}{\Omega\alpha}\left(p_\infty - p_\mathrm{f}\right). \tag{S.21}$$

The oxygen partial pressure first reaches $p_\mathrm{m}$ at the centre of the spheroid at $t = T_2$, corresponding to the end of phase (ii) and start of phase (iii). By definition $R_\mathrm{f}(T_3) > 0$ and $R_\mathrm{m}(T_2) = 0$. Setting $r = 0$ in solution of the oxygen partial pressure for the region $R_{\mathrm{m}}(t) < r < R_{\mathrm{f}}(t)$ of Eq (S.19) and using the oxygen threshold in Eq (S.20.2) we obtain,

$$\mathcal{R}_{\mathrm{m}}^2 = -\frac{1}{R(t)}\left[-R(t)^3 - 3R_{\mathrm{f}}(t)^2\,(\phi-1)\,R(t) + 2R_{\mathrm{f}}(t)^3\,(\phi-1)\right], \tag{S.22}$$

where

$$\mathcal{R}_{\mathrm{m}}^2 = \frac{6k}{\Omega\alpha}\,(p_\infty - p_{\mathrm{m}})\,. \tag{S.23}$$

Setting $\phi = 1$ in Eq (S.22) gives $\mathcal{R}_{\mathrm{m}} = R(t)$. Hence, $\mathcal{R}_{\mathrm{m}}$ is the radius of a monoculture melanoma spheroid when the necrotic region first forms. Note Eq (S.22) is only valid at $t = T_2$.

To obtain expressions for $\mathcal{R}_{\mathrm{f}}$ and $\mathcal{R}_{\mathrm{m}}$ throughout time and in terms of radial measurements, $R(t)$, $R_{\mathrm{f}}(t)$, and $R_{\mathrm{m}}(t)$, we evaluate the solution of the oxygen partial pressure given in Eq (S.19) at $r = R_{\mathrm{f}}(t)$ and $r = R_{\mathrm{m}}(t)$, respectively, and use the oxygen thresholds in Eqs (S.20.1) and (S.20.2) to obtain Eqs (S.24)-(S.25) included on the next page for ease of presentation.

$$\mathcal{R}_{\mathrm{f}}^2 = -\frac{1}{R_{\mathrm{f}}(t)R(t)}\left[-\Big(R_{\mathrm{f}}(t)R(t)^2 + R(t)R_{\mathrm{f}}(t)^2 + \left(2R_{\mathrm{f}}(t)^3 - 2R_{\mathrm{m}}(t)^3\right)\phi - 2R_{\mathrm{f}}(t)^3\Big)\Big(-R_{\mathrm{f}}(t) + R(t)\Big)\right], \qquad t > T_1, \tag{S.24}$$

$$\mathcal{R}_{\mathrm{m}}^2 = -\frac{1}{R_{\mathrm{m}}(t)R(t)}\left[3\phi R_{\mathrm{m}}(t)^3R(t) + \Big(-R(t)^3 - 3\,R_{\mathrm{f}}(t)^2\,(\phi-1)\,R(t) + \left(2\,R_{\mathrm{f}}(t)^3 - 2\,R_{\mathrm{m}}(t)^3\right)\phi - 2\,R_{\mathrm{f}}(t)^3\Big)R_{\mathrm{m}}(t)\right], \quad t > T_2. \tag{S.25}$$

Assuming $R(t)$ and $\mathcal{R}_{\mathrm{f}}$ and $\mathcal{R}_{\mathrm{f}}$ are known, we determine the boundary constraints that define $R_{\mathrm{f}}(t)$ and $R_{\mathrm{m}}(t)$, respectively, through

$$0 = \begin{cases} R_{\mathrm{f}}(t) - 0, & 0 \le t \le T_1, \\ -\mathcal{R}_{\mathrm{f}}^2 - \dfrac{1}{R_{\mathrm{f}}(t)R(t)}\left[-\Big(R_{\mathrm{f}}(t)R(t)^2 + R(t)R_{\mathrm{f}}(t)^2 + 2R_{\mathrm{f}}(t)^3\,(\phi-1)\Big)\Big(-R_{\mathrm{f}}(t) + R(t)\Big)\right], & T_1 < t \le T_2, \\ -\mathcal{R}_{\mathrm{f}}^2 - \dfrac{1}{R_{\mathrm{f}}(t)R(t)}\left[-\Big(R_{\mathrm{f}}(t)R(t)^2 + R(t)R_{\mathrm{f}}(t)^2 + \left(2R_{\mathrm{f}}(t)^3 - 2R_{\mathrm{m}}(t)^3\right)\phi - 2R_{\mathrm{f}}(t)^3\Big)\Big(-R_{\mathrm{f}}(t) + R(t)\Big)\right], & t > T_2, \end{cases} \tag{S.26}$$

$$0 = \begin{cases} R_{\mathrm{m}}(t) - 0, & 0 \le t \le T_2, \\ -\mathcal{R}_{\mathrm{m}}^2 - \dfrac{1}{R_{\mathrm{m}}(t)R(t)}\left[3\phi R_{\mathrm{m}}(t)^3R(t) + \Big(-R(t)^3 - 3\,R_{\mathrm{f}}(t)^2\,(\phi-1)\,R(t) + \left(2\,R_{\mathrm{f}}(t)^3 - 2\,R_{\mathrm{m}}(t)^3\right)\phi - 2\,R_{\mathrm{f}}(t)^3\Big)R_{\mathrm{m}}(t)\right], & t > T_2, \end{cases} \tag{S.27}$$

Given these expressions we determine $\mathrm{d}R_{\mathrm{f}}(t)/\mathrm{d}R(t)$ and $\mathrm{d}R_{\mathrm{m}}(t)/\mathrm{d}R(t)$ by considering phase (ii) and phase (iii) separately. For phase (ii), we implicitly differentiate Eq (S.26) with respect to $R(t)$ to obtain

$$\frac{\mathrm{d}R_{\mathrm{f}}(t)}{\mathrm{d}R(t)} = \frac{(\phi-1)\,R_{\mathrm{f}}(t)^3 + R(t)^3}{R(t)R_{\mathrm{f}}(t)\left[3R_{\mathrm{f}}(t)\phi - 2R(t)\phi - 3R_{\mathrm{f}}(t) + 3R(t)\right]}, \tag{S.28}$$

where as before we set $\phi = V_1^{(2)}(t)/\left(V_1^{(2)}(t) + V_2^{(2)}(t)\right)$. For phase (iii) we implicitly differentiate Eqs (S.26) and (S.27) with respect to $R(t)$, and solve the resulting two coupled differential equations for $\mathrm{d}R_{\mathrm{f}}(t)/\mathrm{d}R(t)$ and $\mathrm{d}R_{\mathrm{m}}(t)/\mathrm{d}R(t)$,

$$\frac{\mathrm{d}R_{\mathrm{f}}(t)}{\mathrm{d}R(t)} = \frac{-R_{\mathrm{f}}(t)\left((\phi-1)\,R_1(t)^3 - R_{\mathrm{m}}(t)^3\phi + R(t)^3\right)}{2R(t)\left[\frac{3}{2}\,(1-\phi)\,R_{\mathrm{f}}(t)^3 + \left(\frac{R_{\mathrm{m}}(t)\phi}{2} + R(t)\left(\phi - \frac{3}{2}\right)\right)R_{\mathrm{f}}(t)^2 - \frac{R_{\mathrm{m}}(t)\phi(R(t)-R_{\mathrm{m}}(t))R_{\mathrm{f}}(t)}{2} - \frac{\phi R_{\mathrm{m}}(t)^2(R(t)-R_{\mathrm{m}}(t))}{2}\right]}, \tag{S.29.1}$$

$$\frac{\mathrm{d}R_{\mathrm{m}}(t)}{\mathrm{d}R(t)} = \frac{-\left(R_{\mathrm{f}}(t)^2 + R_{\mathrm{m}}(t)R_{\mathrm{f}}(t) + R_{\mathrm{m}}(t)^2\right)\left((\phi-1)\,R_{\mathrm{f}}(t)^3 - R_{\mathrm{m}}(t)^3\phi + R(t)^3\right)}{6R(t)\left[\frac{3}{2}\,(1-\phi)\,R_{\mathrm{f}}(t)^3 + \left(\frac{R_{\mathrm{m}}(t)\phi}{2} + R(t)\left(\phi - \frac{3}{2}\right)\right)R_{\mathrm{f}}(t)^2 - \frac{R_{\mathrm{m}}(t)\phi(R(t)-R_{\mathrm{m}}(t))R_{\mathrm{f}}(t)}{2} - \frac{\phi R_{\mathrm{m}}(t)^2(R(t)-R_{\mathrm{m}}(t))}{2}\right]R_{\mathrm{m}}(t)}. \tag{S.29.2}$$

Substituting Eqs (10.1)-(10.6), (S.26), (S.27), S.28, and (S.29) into Eqs (4.1-4.4) gives the final system that we solve numerically.

### S4 Numerical methods

In brief, we solve each mathematical model numerically in Julia. Key algorithms are available on a GitHub repository (https://github.com/ryanmurphy42/Murphy2022CoCulture). For the biphasic model we solve Eq (1) using a second-order explicit Runge–Kutta method, further details in [19]. For the linear model we use the analytical solution (Eq 2).

For compartment-based models, which typically take the form of a system of differential-algebraic equations, we consider each phase in turn. For each phase, we write the governing equations in a mass-matrix differential-algebraic equation form, i.e. $M\mathrm{d}u/\mathrm{d}t = f(u, t; p)$ where $M$ is known as the mass matrix, $u$ is a vector of time-dependent functions, $t$ denotes time which is the independent variable, and $p$ is a vector of parameters. To solve these systems we use the `Rodas(5)` solver, a 5th order A-stable stiffly stable Rosenbrock method with a stiff-aware 3rd order interpolant, that is included in open-source `DifferentialEquations` package [20]. We terminate each phase using a stopping condition that incorporates a small error tolerance denoted by $\epsilon$. For example, we set the terminating condition for phase (i) in the model presented in §4.5 to be $R(t) > \mathcal{R}_\mathrm{f} + \epsilon$. For Greenspan-type models we set $\epsilon = 1 \times 10^{-4}$ to overcome numerical challenges at the start of phase (ii) where $\mathrm{d}R_\mathrm{f}(t)/\mathrm{d}R(t)$ is not defined for $R_\mathrm{f}(t) = 0$ and is very large for small $R_\mathrm{f}(t)$ (Eq S.28). Similarly, at the start of phase (iii) where $\mathrm{d}R_2(t)/\mathrm{d}R(t)$ is not defined for $R_\mathrm{m}(t) = 0$ and is very large for small $R_\mathrm{m}(t)$ (Eq S.29). Initial conditions in phase (ii) and (iii) are determined using the solution at the end of the phase (i) and (ii), respectively. For other models, such as the radial-death model (§S5.1.2) we can set $\epsilon = 0$.

### S5 Additional results

Here we present additional results to support results in the main manuscript. This includes how other mathematical models in the literature are captured as part of our general compartment based tumour modelling framework (§S5.1); comparisons of experimental measurements of spheroid size and simulations of the biphasic model at the MLE for each 1205Lu co-culture condition (§S5.2); comparisons of experimental measurements of spheroid size and simulations of the linear model at the MLE for each WM983B co-culture condition §S5.3); and experimental images that show the temporal evolution of spheroid size and structure for co-culture experiments performed with the 1205Lu melanoma cell line (§S5.4)

#### S5.1 Additional examples using general compartment based modelling framework

In the main manuscript we demonstrate how the reduced Greenspan model is captured within our general framework. Here we present two further examples (i) Greenspan's seminal model [18] and (ii) the radial-death model [21].

##### S5.1.1 Greenspan's model

Greenspan's seminal monoculture model [18] considers one population and three compartments ($I = 1$, $J = 3$) and describes growth with three phases. In phase (i) all cells within the spheroid can proliferate and the spheroid grows exponentially. In phase (ii) the spheroid is composed of a proliferating region at the periphery, $R_1(t) < r < R(t)$, and a proliferation-inhibited or quiescent region, $0 \leq r \leq R_1(t)$. In phase (iii) the spheroid is composed of three compartments: compartment one which is a proliferating region at the periphery, $R_1(t) < r < R(t)$; compartment two which is the proliferation-inhibited or quiescent region, $R_2(t) \leq r \leq R_1(t)$; and compartment three which is the necrotic core, $0 \leq r \leq R_2(t)$. At later times proliferation at the periphery balances mass loss from the necrotic core resulting in a limiting spheroid structure. For consistency with the literature we write $R_0(t)$ as $R(t)$, $R_1(t)$ as $R_\mathrm{i}(t)$, and $R_2(t)$ as $R_\mathrm{n}(t)$ [1, 2]. We define $R_\mathrm{i}(t)$ and $R_\mathrm{n}(t)$ using boundary constraints obtained by considering additional biological mechanisms [2]. Note that here we consider the version of Greenspan's model where the proliferation-inhibited or quiescent region forms before the necrotic region [1, 2, 21]. The alternative formulation, where the necrotic region forms before the proliferation-inhibited region, can also be captured in this general framework.

We assume that in the proliferating region the rate at which cell volume is produced by mitosis per unit volume of living cells is $s$ [day$^{-1}$]. In the necrotic region we assume that the rate at which cell volume is lost from the necrotic core per unit volume of necrotic material is $3s\gamma$ [day$^{-1}$] where $\gamma$ [-] is a dimensionless parameter and the three is included for mathematical convenience and consistency with the literature [18].

Therefore, we prescribe the following functions

$$f_{V_1^{(1)}}\left(R(t);p\right) = sV_1^{(1)}(t) \qquad = s\frac{4\pi}{3}\left(R(t)^3 - R_{\mathrm{i}}(t)^3\right), \tag{S.30.1}$$

$$f_{V_1^{(2)}}\left(R(t);p\right) = 0 \tag{S.30.2}$$

$$f_{V_1^{(3)}}\left(R(t);p\right) = -3s\gamma V_1^{(3)}(t) = -3s\gamma\frac{4\pi}{3}R_{\mathrm{n}}(t)^3. \tag{S.30.3}$$

As described in §S3.3 to define $R_{\mathrm{n}}(t)$ we consider the oxygen partial pressure within the spheroid which gives rise to the boundary constraint in Eq (S.30), rewritten here for clarity,

$$0 = \begin{cases} R_{\mathrm{n}}(t) - 0, & R(t) \leq \mathcal{R}, \\ -\mathcal{R}^2 + R(t)^2 - R_{\mathrm{n}}(t)^2 - \dfrac{2R_{\mathrm{n}}(t)^2}{R(t)}\left(R(t) - R_{\mathrm{n}}(t)\right), & R(t) > \mathcal{R}. \end{cases} \tag{S.31}$$

Similarly, to define $R_{\mathrm{i}}(t)$ we consider the oxygen partial pressure within the spheroid and an oxygen partial pressure threshold $p_{\mathrm{i}}$ [%]. For the proliferation-inhibited region to form $p_{\mathrm{i}} > p_{\mathrm{n}}$. This gives rise to the following boundary constraint

$$0 = \begin{cases} R_{\mathrm{i}}(t) - 0, & R(t) \leq \mathcal{R}_{\mathrm{i}}, \\ -\mathcal{R}_{\mathrm{i}}^2 + R(t)^2 - R_{\mathrm{i}}(t)^2 - 2R_{\mathrm{n}}(t)^3\left(\dfrac{1}{R_{\mathrm{i}}(t)} - \dfrac{1}{R(t)}\right), & R(t) > \mathcal{R}_{\mathrm{i}}, \end{cases} \tag{S.32}$$

where $\mathcal{R} = [6k(p_\infty - p_{\mathrm{i}})/(\alpha\Omega)]^{1/2}$ is the size of the spheroid when the proliferation-inhibited region first forms. Alternatively, one can define $R_{\mathrm{i}}(t)$ by considering production and diffusion of diffusible waste. These waste mechanisms gives rise to the same boundary constraint as in Eq (S.32) but with $\mathcal{R}_{\mathrm{i}} = 6\beta_{\mathrm{i}}\kappa/P$, where $\beta_{\mathrm{i}}$ is a waste threshold, $\kappa$ is the diffusivity of the diffusible waste, and $P$ is rate of production of waste per unit volume [2].

Substituting Eqs (S.30.1)- (S.30.3) and Eqs S.31-S.31) into Eqs (4.1-4.4) and simplifying we can solve for $R(t)$ directly using,

$$R(t)^2\frac{\mathrm{d}R(t)}{\mathrm{d}t} = \frac{s}{3}\left(R(t)^3 - R_{\mathrm{i}}(t)^3\right) - \lambda R_{\mathrm{n}}(t)^3, \tag{S.33}$$

$$0 = \begin{cases} R_{\mathrm{i}}(t) - 0, & R(t) \leq \mathcal{R}_{\mathrm{i}}, \\ -\mathcal{R}_{\mathrm{i}}^2 + R(t)^2 - R_{\mathrm{i}}(t)^2 - 2R_{\mathrm{n}}(t)^3\left(\dfrac{1}{R_{\mathrm{i}}(t)} - \dfrac{1}{R(t)}\right), & R(t) > \mathcal{R}_{\mathrm{i}}, \end{cases} \tag{S.34}$$

$$0 = \begin{cases} R_{\mathrm{n}}(t) - 0, & R(t) \leq \mathcal{R}, \\ -\mathcal{R}^2 + R(t)^2 - R_{\mathrm{n}}(t)^2 - \dfrac{2R_{\mathrm{n}}(t)^2}{R(t)}\left(R(t) - R_{\mathrm{n}}(t)\right), & R(t) > \mathcal{R}. \end{cases} \tag{S.35}$$

In this model there are five parameters $\theta = (R(T), s, \gamma, \mathcal{R}, \mathcal{R}_{\mathrm{i}})$, where $R(T)$ [µm] is the initial radius of the spheroid at formation that we treat as a constant parameter.

##### S5.1.2 Radial-death model

The radial-death model introduced in [21] describes one population with two compartments ($I = 1$, $J = 2$), captures key characteristics of Greenspan's model, and describes spheroid growth with two phases. In phase (i) all cells within the spheroid proliferate and the spheroid grows exponentially. In phase (ii) the spheroid is composed of two compartments: compartment one which is the proliferating region at the periphery, $R_\mathrm{n}(t) < r < R(t)$; and compartment two which is the necrotic core, $0 \leq r \leq R_\mathrm{n}(t)$. In phase (ii) the size of the proliferating region, $R(t) - R_\mathrm{n}(t)$, is assumed to be constant and denoted $R_\mathrm{d}$ [µm]. This defines a boundary constraint on the size of the proliferating region, or equivalently of the necrotic region.

Identical to the monoculture reduced Greenspan model (§4.2), we assume that in the proliferating region the rate at which cell volume is produced by mitosis per unit volume of living cells is $s$ [day$^{-1}$]. In the necrotic region we assume that the rate at which cell volume is lost from the necrotic core per unit volume of necrotic material is $3s\gamma$ [day$^{-1}$] where $\gamma$ [-] is a dimensionless parameter and the three is included for mathematical convenience and consistency with the literature [18]. The prescribed functions for this model are

$$f_{V_1^{(1)}}\left(R(t); p\right) = sV_1^{(1)}(t) = s\frac{4\pi}{3}\left(R(t)^3 - R_\mathrm{n}(t)^3\right) \tag{S.36.1}$$

$$f_{V_1^{(2)}}\left(R(t); p\right) = -3s\gamma V_1^{(2)}(t) = -3s\gamma\frac{4\pi}{3}R_\mathrm{n}(t)^3 \tag{S.36.2}$$

$$R_\mathrm{n}(t) = \max\left(0, R(t) - R_\mathrm{d}\right) = \begin{cases} 0, & R(t) < R_\mathrm{d}, \\ R(t) - R_\mathrm{d}, & R(t) \geq R_\mathrm{d}. \end{cases} \tag{S.36.3}$$

In this model there are four parameters, $\theta = (R(T), s, \gamma, R_\mathrm{d})$, where $R(T)$ [µm] is the initial radius of the spheroid at formation that we treat as a constant parameter.

Substituting the prescribed functions (Eqs (S.36.1)-(S.36.3) and (S.14)) into general model Eqs (4.1-4.4) and simplifying we can solve for $R(t)$ directly using,

$$\frac{\mathrm{d}R(t)}{\mathrm{d}t} = \frac{1}{3R(t)^2}\left[s\left(R(t)^3 - \max\left(0, R(t) - R_\mathrm{d}\right)^3\right) - 3s\gamma\max\left(0, R(t) - R_\mathrm{d}\right)^3\right]. \tag{S.37}$$

#### S5.2 Spheroid formation and growth (1205Lu)

In the main manuscript we present profile likelihoods for the formation time, $T$ [days], of 1205Lu co-culture spheroids (Fig 5). In the main manuscript we state that, for each 1205Lu co-culture condition, we observe excellent agreement between experimental measurements of spheroid size, $R(t)$, and the biphasic model simulated at the MLE. We now show this excellent agreement in Fig S6. Here we also show profile likelihoods for the other model parameters. These profiles suggest that each parameter is practically identifiable for each condition since the univariate profiles are each well-formed around a single peak (Fig S7).

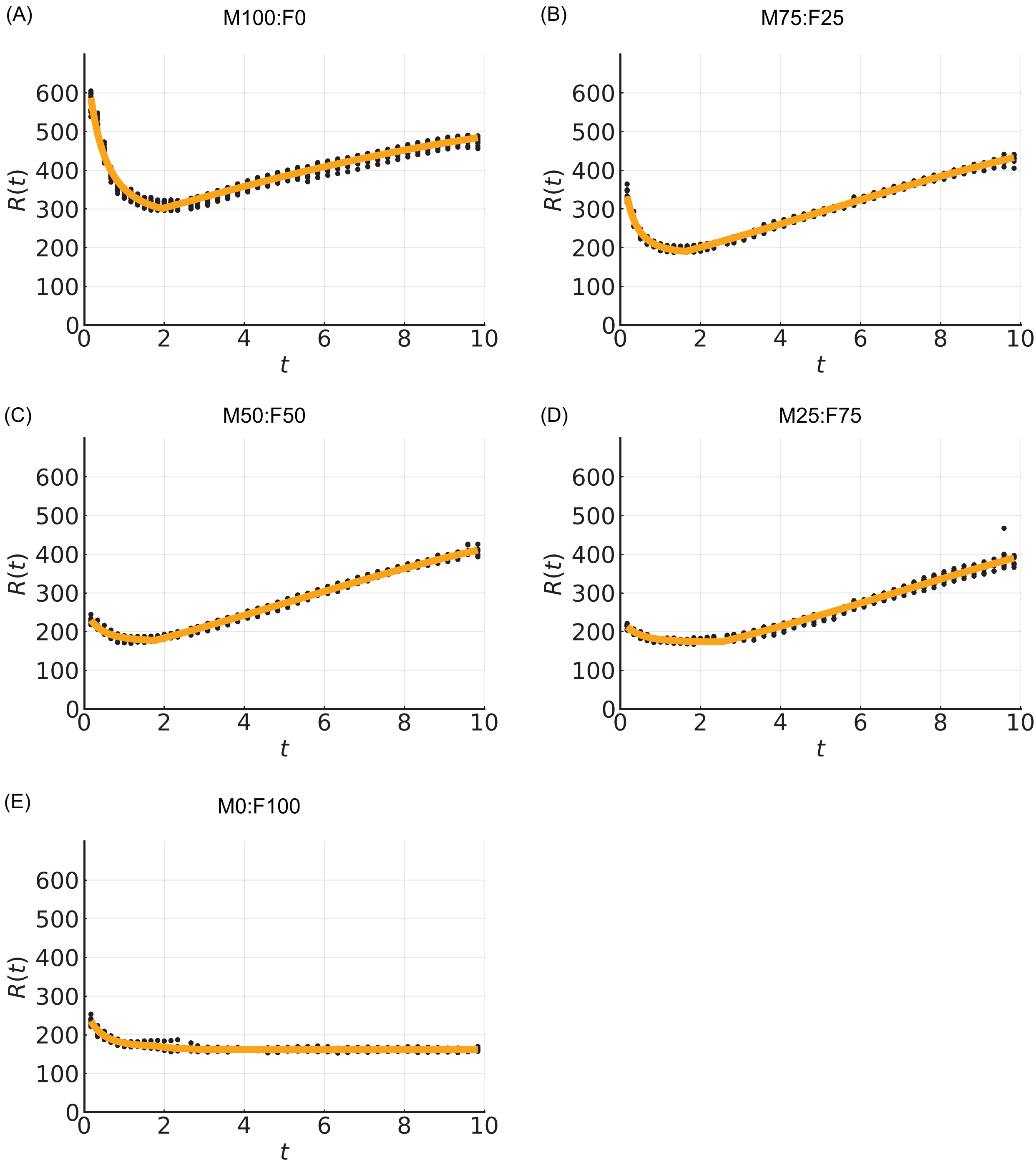

**Figure S6: Comparison of experimental measurements of spheroid size, $R(t)$, (black circles) and the biphasic model simulated at the MLE (orange line)**. Results shown for 1205Lu co-culture spheroids.

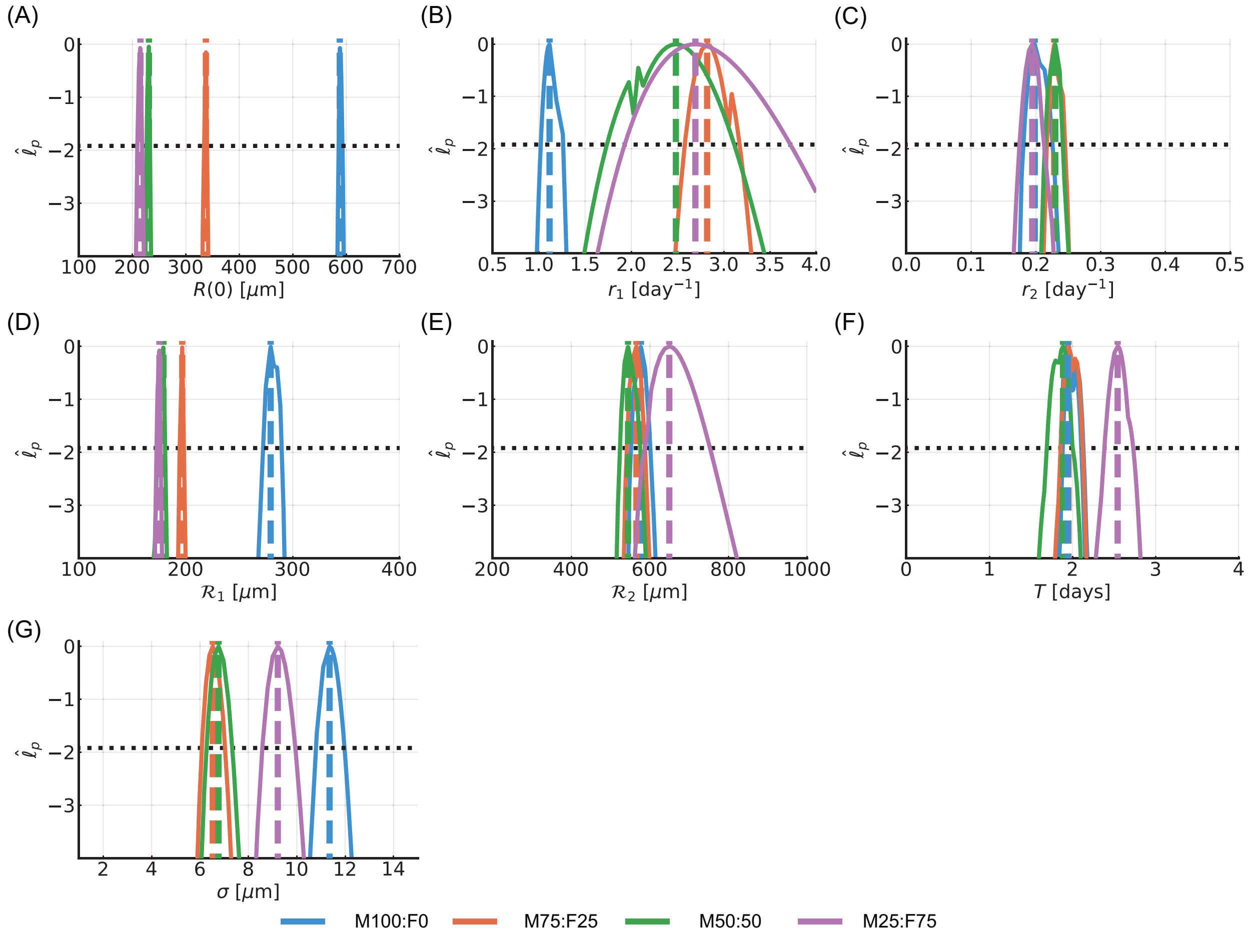

**Figure S7: Profile likelihoods for the biphasic model.** Results shown for 1205Lu co-culture spheroids. (A-F) Profile likelihoods for (A) $R(0)$ [µm], (B) $r_1$ [days$^{-1}$], (C) $r_2$ [days$^{-1}$], (D) $\mathcal{R}_1$ [µm], (E) $\mathcal{R}_2$ [µm], (F) $T$ [days]. The approximate 95% confidence interval threshold shown with horizontal black-dashed line. Throughout conditions are M100:F0 (blue), M75:F25 (orange), M50:F50 (green), and M25:F75 (magenta).

##### S5.3 Temporal evolution of spheroid size and structure (WM983B)

In the main manuscript, for each WM983B co-culture condition, we use the linear model. Here we show that we observe excellent agreement between experimental measurements of spheroid size, $R(t)$, and the linear model simulated at the MLE (Fig S8).

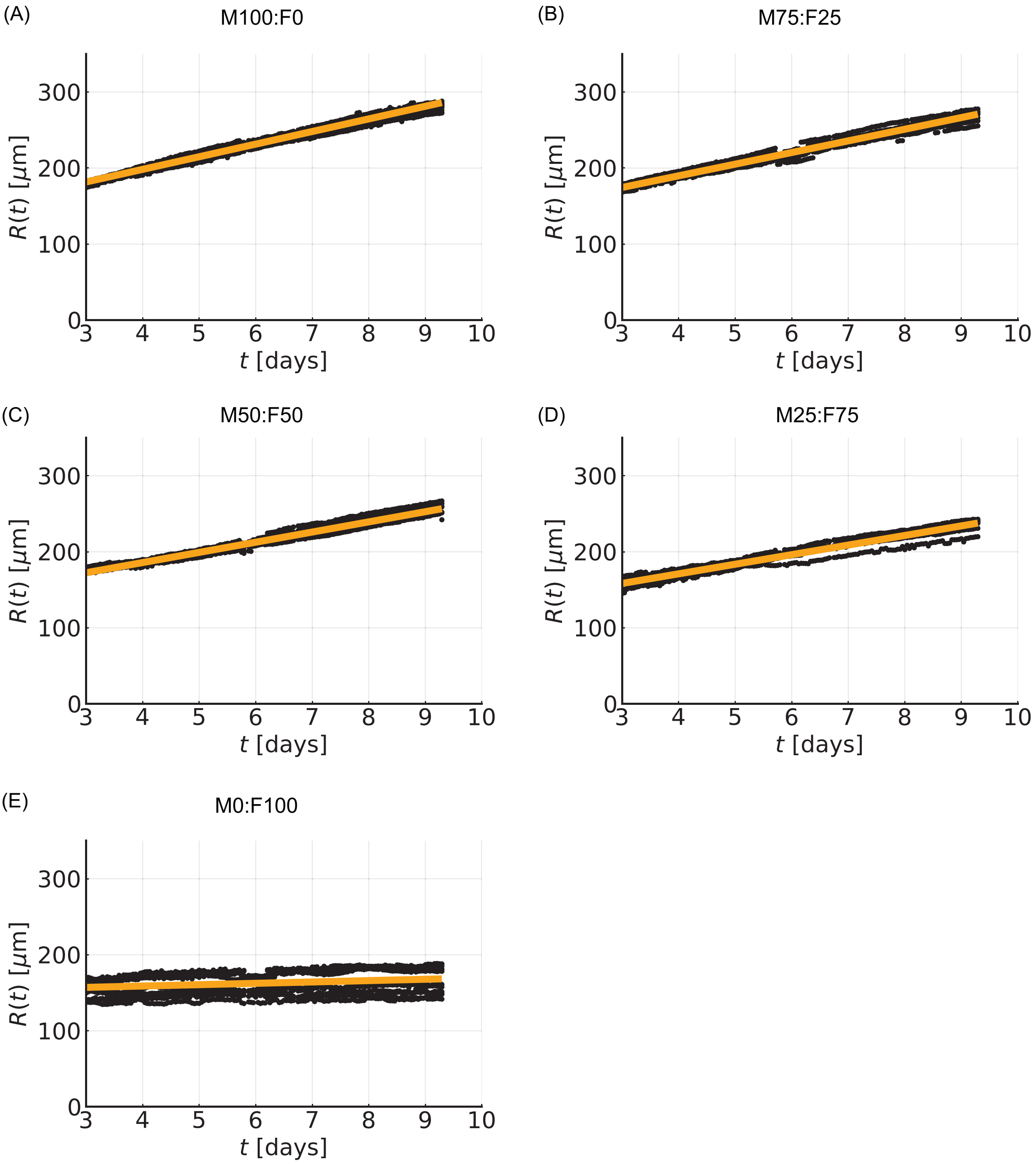

**Figure S8: Comparison of experimental measurements of spheroid size, $R(t)$, (black circles) and the linear model simulated at the MLE (orange line)**. Results shown for WM983B co-culture spheroids.

#### S5.4 Temporal evolution of spheroid size and structure (1205Lu)

Here we present experimental images that show the temporal evolution of spheroid size and structure for co-culture experiments performed with the 1205Lu melanoma cell line (Figs S9, S10). For 1205Lu co-culture spheroids we observe that fibroblasts are primarily located at the central region of the spheroid for the entire experimental duration. This is in contrast to results for WM983B co-culture spheroids where fibroblasts are present throughout the spheroid at early times but are only present at the central region at later times.

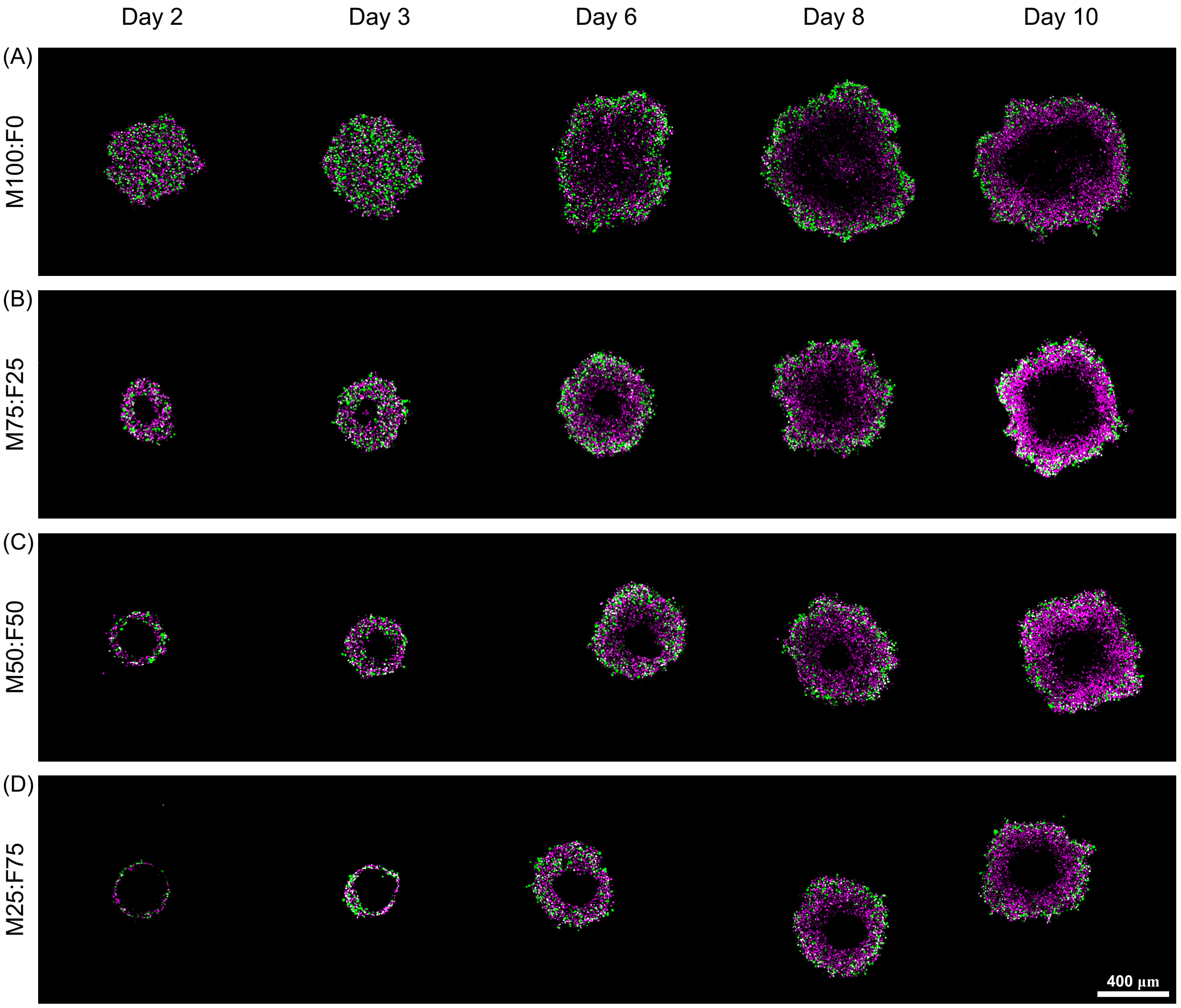

**Figure S9: Spatio-temporal evolution of tumour spheroid structure.** Confocal microscopy images of each spheroid's equatorial plane for (A) M100:F0, (B) M75:F25, (C) M50:F50, and (D) M25:F75 spheroids. Results shown for the 1205Lu melanoma cell line. FUCCI signals are shown with magenta and green. Presence of FUCCI signal indicates living cells whereas large regions at the centre of spheroids that lack FUCCI signals indicate a necrotic core. Fibroblasts are not shown so that it is easier to visualise the necrotic core and overall size.

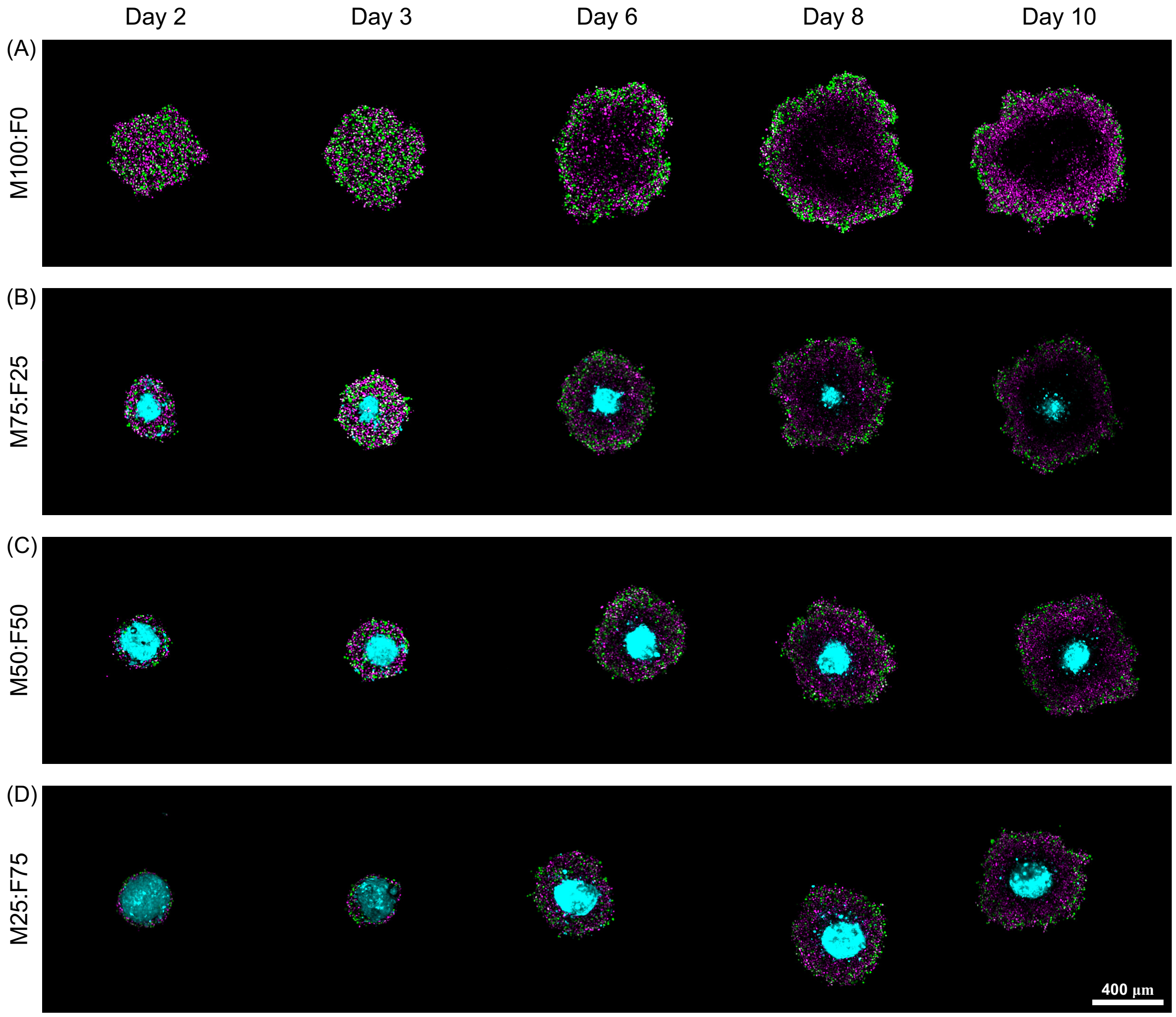

**Figure S10: Spatio-temporal evolution of tumour spheroid structure including fibroblast marker.** Confocal microscopy images of each spheroid's equatorial plane for (A) M100:F0, (B) M75:F25, (C) M50:F50, and (D) M25:F75 spheroids. Results shown for the 1205Lu melanoma cell line. These are same spheroids as shown in Fig (S9) now with the fibroblast marker shown in cyan. The fibroblast marker stains the entire cell whereas the FUCCI signal is only present at the cell nucleus.
